# Proteomic identification of the interactome of stalled ribosome nascent chain complexes translating the thylakoid membrane protein D1

**DOI:** 10.1101/2022.03.18.484870

**Authors:** Dominique S. Stolle, Paul Treimer, Jan Lambertz, Lena Osterhoff, Annika Bischoff, Beatrix Dünschede, Anja Rödiger, Christian Herrmann, Sacha Baginsky, Marc M. Nowaczyk, Danja Schünemann

## Abstract

The synthesis of multi-span thylakoid membrane proteins initiates at ribosomes off the membrane. Subsequently, the ribosome nascent chain complexes (RNCs) are transferred to the translocase machinery in the thylakoid membrane for cotranslational protein insertion. These steps require finely tuned mechanisms for protein processing, quality control, and targeting to prevent misfolding or aggregation and to ensure efficient transfer of the nascent chain to the insertion machinery. However, little is known about the regulatory network underlying these processes. To identify factors specifically involved in the cotranslational biogenesis of the reaction center protein D1 of photosystem II we established a chloroplast-derived *in vitro* translation method that allows the production and affinity purification of stalled RNCs bearing nascent chains of D1 of different defined lengths. Stalled RNCs translating the soluble ribosomal subunit uS2c were affinity-purified for comparison. Quantitative tandem-mass spectrometry revealed a set of about 120 proteins specifically associated with D1 RNCs. The interactome includes proteins with broad functions in protein processing, biogenesis and metabolic pathways, such as chlorophyll biosynthesis. We identified STIC2 as a new factor specifically associated with D1 RNCs. Furthermore, our results demonstrated that the interaction of STIC2 with the thylakoid insertase Alb3 and its homologue Alb4 is mediated by the conserved motif III within the C-terminal regions of Alb3 and Alb4. Our data suggest that STIC2 is involved in cotranslational substrate delivery at the thylakoid membrane by coordinating the binding of the D1 RNCs to the insertase machinery.

## Introduction

The biogenesis and maintenance of the photosynthetically active thylakoid membrane of chloroplasts requires the cotranslational targeting of plastid-encoded thylakoid membrane proteins, which mediate electron transport and ATP synthesis during photosynthesis (Jagendorf and Michaels, 1990; Kim et al., 1991; Zhang et al., 1999; Zoschke and Barkan, 2015). The synthesis of these proteins begins at soluble ribosomes that can be indirectly tethered to the thylakoid membrane by mRNA-associated factors. With the emergence of the first transmembrane helix (TMH) of the nascent peptide, a tight nuclease-resistent association between the translating ribosome and the thylakoid membrane is established (Zoschke and Barkan, 2015). This is most likely due to insertion of the nascent chain into the cpSec1/Alb3 membrane insertase machinery and contacts with the lipid bilayer. The transition from soluble to membrane-bound translating ribosomes needs to be orchestrated by many factors that are involved in nascent chain processing, insertion-competent folding, quality control, and sorting of the ribosome nascent chain complexes (RNCs) to the target membrane. Chloroplasts harbor an extensive network of processing enzymes, molecular chaperones and proteases which contributes to protein modification and maturation (Trösch et al., 2015a; van Wijk, 2015; Breiman et al., 2016; Nishimura et al., 2017; Ries et al., 2020; Sun et al., 2021). However, the question of which factors are specifically required for cotranslational thylakoid membrane protein biogenesis and how they exert their molecular function remains largely unexplored. Recently, the chloroplast trigger factor-like protein 1 (TIG1), a homolog of the bacterial chaperone trigger factor, that binds at the bacterial ribosomal polypeptide tunnel exit site and promotes folding of a broad subset of nascent chains, was shown to be partially associated with chloroplast ribosomes but its potential role as chaperone and its substrate specificity has not been defined yet (Olinares et al., 2010; Rohr et al., 2019). A recent proteome analysis of affinity-purified chloroplast ribosomes from the green alga *Chlamydomonas reinhardtii*, demonstrated that Hsp70B, Hsp90C and the chaperonin Cpn60 are associated with translating ribosomes (Westrich et al., 2021). Factors involved in sorting of RNCs to the thylakoid membrane comprise the chloroplast 54 kDa signal recognition particle subunit (cpSRP54), which is homologous to cytosolic SRP54 targeting factors in eukaryotes and prokaryotes (Akopian et al., 2013; Ziehe et al., 2018). Substrate proteins that depend on cpSRP54 for efficient cotranslational targeting comprise central photosynthetic proteins, such as the core proteins of the photosystems I and II (PsaA, PsaB, D1, D2) or the cytochrome *b*_*6*_*f* complex subunit PetB (Hristou et al., 2019). CpSRP54 can bind directly to the ribosome via the ribosomal subunit uL4c and initiates targeting at early stages of translation (Hristou et al., 2019). CpSRP54 is also able to contact the nascent chain of D1 after its emergence from the ribosomal exit tunnel (Nilsson et al., 1999; Nilsson and van Wijk, 2002). An alternative cotranslational targeting mechanism was described for the chloroplast-encoded cytochrome *b*_*6*_*f* subunit cytochrome *f*. Cytochrome *f* is synthesized with a cleavable N-terminal signal peptide and is targeted by chloroplast SecA (cpSecA), a homologue of the bacterial ATPase SecA, which mediates targeting of preproteins to the plasma membrane Sec machinery for translocation (Voelker and Barkan, 1995; Voelker et al., 1997; Röhl and van Wijk, 2001; Zoschke and Barkan, 2015; Fernandez, 2018). Consistently, cpSecA was identified in the ribosome-associated proteome of *Chlamydomonas* chloroplasts (Westrich et al., 2021).

In addition to folding, quality control, and sorting, ribosome-associated factors are also important for the precise regulation of protein synthesis in response to changing environmental and internal cellular conditions. They include translation factors, ribosome recycling, hibernation and biogenesis factors and transcript-specific RNA binding proteins (Sun and Zerges, 2015; Zoschke and Bock, 2018; Trösch and Willmund, 2019). Furthermore, there is growing evidence that metabolic enzymes can physically interact with components of the translation machinery. In a recent study, it was found that various enzymes, e.g. of carbon, amino acid or chlorophyll metabolism, are associated with isolated chloroplast ribosomes (Westrich et al., 2021; Trösch et al., 2022). In line with this, previous studies have demonstrated a direct association of an enzyme of chlorophyll biosynthesis with chloroplast ribosomes and an enzyme of fatty acid biosynthesis with D1-encoding mRNA, affecting its localized translation (Kannangara et al., 1997; Bohne et al., 2013).

Yet even though previous studies made progress in deciphering the ribosome interactome, the proteome specifically associated with a translating ribosome depending on the nature of the nascent chain and the question of chloroplast ribosome heterogeneity remain unexplored. Here, we describe a method for the *in vitro* generation and affinity purification of stalled chloroplast ribosomes translating either different chain lengths of the thylakoid membrane protein D1 or the soluble ribosomal subunit uS2c using a homologous translation system. Mass spectrometry-based proteome analysis of the purified RNCs revealed a set of proteins specifically associated with D1-translating ribosomes including factors involved in nascent chain modifications and sorting as well as metabolic enzymes. We also identified STIC2 as a novel factor specifically associated with D1-translating ribosomes and analyzed its interaction with the insertase machinery in the thylakoid membrane.

## Results

### Production and purification of stalled ribosome complexes translating D1 and uS2c nascent chains

The production and isolation of stalled chloroplast RNCs containing varying chains builds on a *Pisum sativum* chloroplast-derived *in vitro* translation system using truncated mRNAs lacking a stop codon, which was previously used to generate stable RNC translation intermediates of D1 (Nilsson et al., 1999; Nilsson and van Wijk, 2002; Walter et al., 2015). Here, we adapted the method by incorporating a twin-strep-tag (TST) coding sequence (CDS) into the D1-encoding *psbA* mRNA allowing specific isolation of D1 RNCs by affinity purification. The TST was positioned between amino acids 25 and 26 of the D1 N-terminus (Figure 1A) as it was described that the integrity of the *psbA* 5’ untranslated region (UTR) and *psbA* coding region is crucial for translational initiation (Nakamura et al., 2016). Furthermore, initial attempts to translate D1 with an N-terminal TST, positioned directly downstream the start codon were unsuccessful (data not shown). To identify factors specifically associated with D1-translating ribosomes, ribosome complexes translating the soluble 30S ribosomal subunit uS2c were generated for comparison. Here, the TST was positioned between amino acids 17 and 18 of the uS2c peptide (Figure 1B). To characterize possible differences in the ribosome interactome as a function of different translational stages, we used truncated *psbA* mRNAs of various lengths resulting in short, medium or long nascent TST-D1 peptides, with a length of 56 and 69, 108 and 136, or 195 and 291 amino acid residues, respectively. The numbers correspond to the D1 amino acid sequence and the peptides are hereafter referred to as TST-D1 (56) to TST-D1 (291). As the ribosomal exit tunnel accommodates approximately 40 residues, the short nascent chain constructs represent translational stages in which the first TMH of D1 is expected to be fully buried inside the tunnel, while the medium-length constructs are designed to expose the first TMH outside of the ribosomal tunnel. The long nascent chain constructs represent translational stages in which at least two TMHs are exposed from the ribosome (Figure 1C). For uS2c, a translation intermediate encoding 158 amino acids of uS2c was generated, referred to as TST-uS2c (158) (Figure 1D).

**Figure 1.**
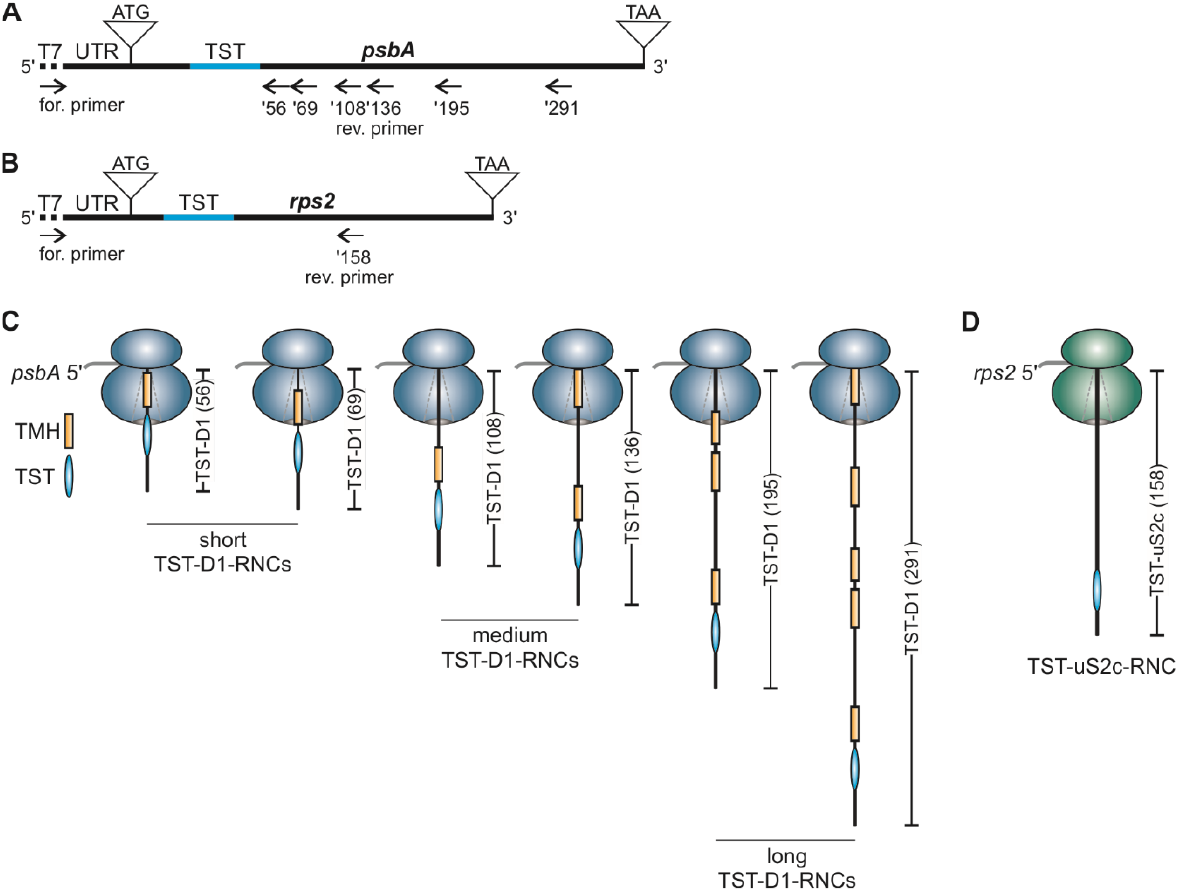
Generation of stalled affinity-tagged ribosome nascent chain complexes (**A**) Schematic representation of the psbA cDNA with T7 promotor sequence (dotted line) at the 5’ end and the endogenous psbA 5’ UTR (87 nt) used for PCR production of truncated templates for *in vitro* transcription into mRNA. A 90 nt twin-strep-tag (TST) coding sequence (blue line) was inserted at position +76 of the psbA sequence. The reverse primers (‘56-’291), which lack a stop codon, determine the length of the PCR product and are named according to the number of D1 amino acids encoded by the corresponding mRNA. (**B**) As in A, the rps2 cDNA possesses a T7 promotor sequence and the endogenous 5’ UTR (90 nt). The TST coding sequence was inserted at position +52. The reverse primer lacks a stop codon and results in a truncated PCR product that corresponds to 158 amino acids of the uS2c protein. (**C&D**) Schematic ribosome nascent chain (RNC) complexes generated by *in vitro* translation of mRNA from truncated templates as shown in A and B. An internal TST is used for the affinity purification of the complexes. The nascent peptides of the D1 protein comprise at least one transmembrane helix (TMH) that is buried in the ribosome peptide tunnel or is exposed to the surrounding environment depending on the nascent peptide length (**C**). The nascent peptide of the soluble uS2c protein lacks any hydrophobic TMH (**D**).

Previously, we demonstrated that D1 truncations generated by the homologous translation system are stably associated with ribosomes (Walter et al., 2015). To verify that the D1 truncations containing a TST and the newly generated uS2c truncation are also ribosome-associated, translation reactions of TST-D1 (136), TST-D1 (195) and TST-uS2c (158) were spun on 1 M sucrose cushions. Western blot analyses of the ultracentrifugation pellets using α-Strep antibodies and antibodies against the ribosomal subunit uL4 demonstrated that the nascent chains of D1 and uS2c cosedimented with the ribosomes through the sucrose cushion (Figure 2A). Next, we examined, whether the TST-tagged RNCs could be purified using streptactin-coupled magnetic beads. To this end, radiolabeled *in vitro* translation reactions of various TST-D1 truncations (56, 108,136,195 and 291) and TST-uS2c (158) were subjected to affinity-purification. The untagged truncation D1 (195) was used as a control. As expected, the TST-tagged D1 truncations and TST-uS2c (158) were detected in the eluates, while the untagged D1 truncation remained in the supernatant (Figure 2B).

**Figure 2.**
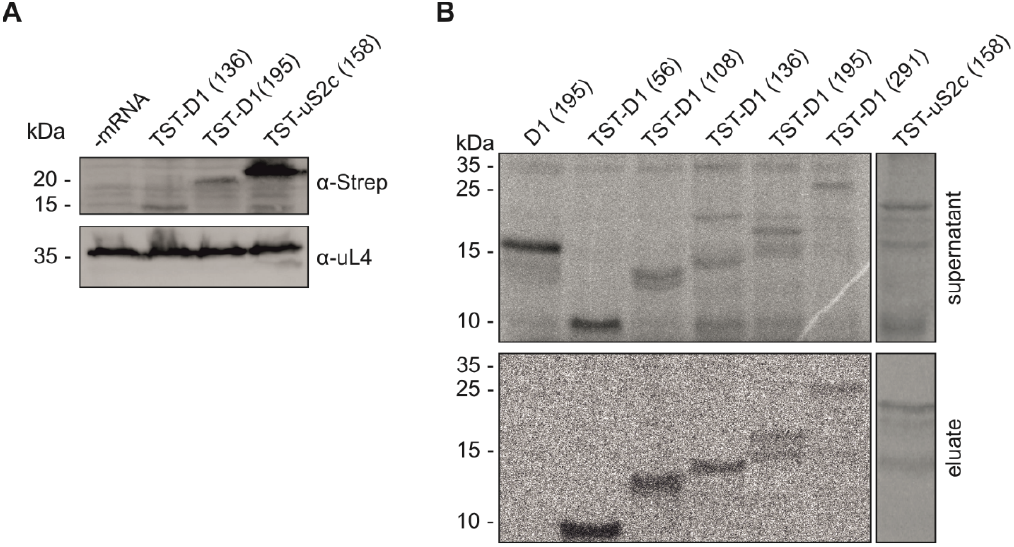
Enrichment and affinity purification of ribosomes translating TST-tagged peptides Generation of stalled RNCs with nascent TST-D1 peptides of different lengths and TST-uS2c (158) using the pea chloroplast-derived *in vitro* translation system. (**A**) Enrichment of TST-D1 (136), TST-D1 (195), and TST-uS2c (158) RNCs via 1 M sucrose cushion centrifugation. A translation sample lacking mRNA was used as a control. The sedimented protein complexes were separated with SDS-PAGE and applied to Western blot. RNCs with 15 kDa and 20 kDa TST-D1 or 22 kDa TST-uS2c peptides were detected with antibody against the TST and the ribosomal protein uL4. (**B**) In the presence of radiolabeled methionine, truncated D1 and uS2c peptides were synthesized *in vitro* and subsequently incubated with MagStrep XT magnetic beads for affinity based purification. A truncated D1 (195) peptide lacking the TST was synthesized as a control for the affinity purification (first lane). Unbound RNCs were removed with the supernatant in a magnetic separator. The magnetic beads were washed and remaining RNCs were subsequently released with native elution. Supernatant (upper panel) and eluate (lower panel) were subjected to SDS-PAGE and synthesized peptides were detected by autoradiography.

### Mass spectrometric analysis of the purified RNCs

To identify factors associated with ribosomes translating D1, affinity-purified RNCs of the short, medium and long TST-D1 intermediates and TST-uS2c (158) were applied to tandem mass spectrometry. Data were analyzed using MaxQuant and a *Pisum sativum* database (see Materials and Methods). *Arabidopsis thaliana* homologs were assigned using the best match in Blast. A total of 242 proteins were identified. Of these proteins, 119 showed no significant quantitative differences in any of the short, medium and long TST-D1 samples compared to the TST-uS2 (158) sample, whereas the remaining 123 proteins were significantly enriched (63 proteins) or exclusively detected (60 proteins) in at least one of the different chain length samples of D1 (Figure 3A, Supplemental Figure 1, Supplemental Table 1 and Supplemental Table 2). Of the 123 putatively D1 RNCs-associated proteins, a core set of 25 proteins were enriched in all D1 chain length samples, while the remaining 98 proteins were assigned to either the short, medium or long chain lengths or an overlap between two groups (Figure 3B, Supplemental Table 2). As expected, due to the TST-D1 nascent chains in the short, medium and long D1 samples peptides of D1 were exclusively detected in the D1 samples (Supplemental Table 2). As uS2 is a core ribosomal subunit, the detection of uS2c peptides in the uS2c sample as well as in D1 samples was also anticipated (Supplemental Table 1). Notably, peptides belonging to the twin-strep-tagged version of uS2c could only be detected in the corresponding sample (data not shown). Chloroplast ribosomes consist of 57 core proteins, of which 24 compose the small 30S subunit and 33 the large 50S subunit (Ban et al., 2014; Bieri et al., 2017). Successful purification and comparative quantitative mass spectrometric analysis of the D1 and uS2c RNCs should therefore result in a large number of ribosomal proteins in the set of proteins showing no enrichment in any of the RNCs. In line with this, 34 core ribosomal proteins (13 subunits of the small subunit and 21 of the large subunit) were detected in the set of the 119 unenriched proteins (Figure 3A, Supplemental Figure 1, Supplemental Table 3). Together with 3 ribosomal proteins that were enriched in the long TST-D1 chains, 65 % of the core ribosomal proteins were covered in the mass spectrometry data set.

**Figure 3.**
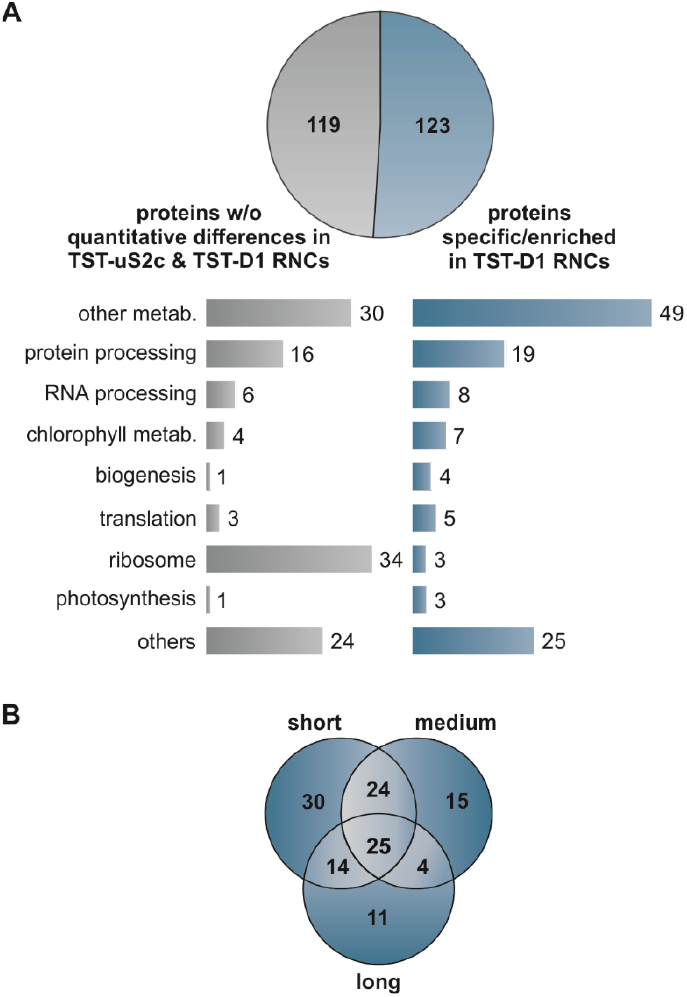
RNC associated proteins identified by MS/MS (**A**) Affinity purification and label-free quantitative tandem mass spectrometry identified 242 proteins in RNC complexes translating TST-uS2c (158) and TST-D1 of different lengths (TST-D1 (56), (69), (108), (136) and (195)). Of these proteins, 123 were significantly enriched or exclusively present in TST-D1 RNCs. The other 119 proteins showed no significant quantitative differences between the uS2c and D1 samples. Identified proteins were functionally categorized. (**B**) The enriched and exclusively assigned proteins of the TST-D1 RNCs were visualized in a Venn diagram showing their distribution among the RNCs with short (56, 69), medium (108, 136), and long (195, 291) nascent TST-D1 peptides or overlaps of these groups.

The 123 D1 RNCs-associated factors include 10 factors with uncharacterized function and 113 factors which can be classified in different categories covering protein/RNA processing, translation, biogenesis and metabolisms (Figure 3A).

Four factors were classified as biogenesis factors (Table 1). As expected, this group includes cpSRP54, whose ribosome association and function in cotranslational targeting has been demonstrated (Nilsson et al., 1999; Nilsson and van Wijk, 2002; Piskozub et al., 2015; Walter et al., 2015; Hristou et al., 2019). CpSRP54 was enriched in all nascent chain lengths of TST-D1-RNCs indicating that cpSRP54 associates with D1 translating ribosomes independent of the D1 chain length. Another biogenesis factor, that was specifically detected in the short, medium and long TST-D1 RNCs is the stromal suppressor of *tic40* protein 2 (STIC2) (Table 1). Recent data indicated that STIC2 and the thylakoid membrane protein Alb4, a homolog of the Alb3 insertase, cooperate in a common cpSRP-dependent pathway in thylakoid membrane biogenesis (Bédard et al., 2017). Furthermore, we detected the fuzzy onions-like (FZL) protein, and the fructokinase-like 1 (FLN1) protein in two (small/medium) and all chain length samples, respectively. Both proteins have been previously characterized as thylakoid membrane or chloroplast biogenesis factors (Gao et al., 2006; Gilkerson et al., 2012; Börner et al., 2015; Liang et al., 2018).

**Table 1.** Chloroplast proteins associated with D1 RNCs and functional in chloroplast or thylakoid biogenesis Mass spectrometry analyses and label-free quantification of purified RNCs showed a D1 specific enrichment of chloroplast biogenesis factors. Asterisks mark proteins that were exclusively found in TST-D1 RNCs. Peptide chain length of TST-D1 RNCs: short (s), medium (m), long (l).

| Uniprot ID | Uniprot Name | Function | Razor Psat | Potential AT | enriched in D1 RNC |
| --- | --- | --- | --- | --- | --- |
| P37107 | Signal recognition particle 54 kDa protein (cpSRP54) | Protein targeting | Psat3g032320.1 | AT5G03940.1 | s, m, l |
| O82230 * | Suppressor of tic40 protein 2 (STIC2) | Thylakoid biogenesis | Psat1g022480.1 | AT2G24020.1 | s, m, l |
| Q1KPV0 * | Probable transmembrane GTPase FZO-like (FZL) | Thylakoid biogenesis | Psat6g060000.3 | AT1G03160.1 | s, m |
| Q9M394 * | Fructokinase-like 1 (FLN1/SCKL1) | Chloroplast biogenesis | Psat7g243520.1 | AT3G54090.1 | s, m, l |

To verify that cpSRP54, STIC2, FZL and FLN1 are specifically associated with D1 translating ribosomes stalled translation products of TST-D1 (136) and as a control TST-uS2 (158) were affinity purified and subjected to immunoblot analysis. In both samples, the equal abundance of ribosomes was demonstrated using antibodies directed against the ribosomal subunits uL4 and uL24 (Figure 4). The twin-strep-tagged D1 and uS2 nascent chains were immunologically detected using *α*-Strep antibodies. Here, however, a longer chain length variant of D1 was used because the *α*-Strep antibody showed a nonspecific signal that overlapped with TST-D1 (136). Notably, cpSRP54 and STIC2 were exclusively detected and FZL and FLN1 were clearly enriched in the TST-D1 (136) sample supporting the mass spectrometry data (Figure 4).

**Figure 4.**
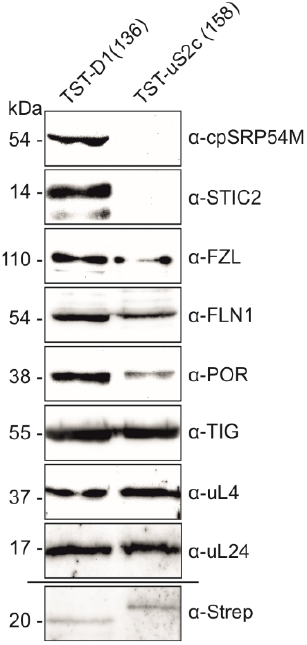
Immunological analyses of purified ribosome nascent chain complexes Stalled RNCs of TST-D1 (136) and TST-uS2c (158) were generated using the chloroplast-derived *in vitro* translation system and purification was performed with StrepTrap XT columns. For immunological analyses equal amounts of purified complexes were separated by SDS-PAGE and transferred onto nitrocellulose membrane. Nonspecific binding of the α-Strep antibody at the size of TST-D1 (136) prevented the detection of the twin-strep-tagged nascent peptide. The purification of RNCs with a longer TST-D1 (195) peptide allowed the immunological detection of the tagged nascent chains with α-Strep antibody.

Recent reports indicated that chloroplast ribosomes exhibit a large interaction network that connects the protein synthesis machinery with various metabolic pathways (Westrich et al., 2021; Trösch et al., 2022). Consistently, we identified among the 123 factors enriched in D1 RNCs 55 proteins that could be assigned to different metabolic pathways (Supplemental Table 2). As we considered an association of metabolic enzymes with ribosomes as a function of nascent chain length rather unlikely and to apply stringent parameters, we restricted this group to factors that were enriched in all chain length samples. This group comprises 10 proteins belonging to carbon, amino acid, chlorophyll, terpenoid and purine metabolism (Table 2). Because of a potential link between the chlorophyll metabolism and the biogenesis of chlorophyll-binding thylakoid membrane proteins such as D1 (Wang and Grimm, 2021) we focused on the chlorophyll biosynthesis enzymes, magnesium protoporphyrin IX methyltransferase (CHLM) and protochlorophyllide reductase (POR) and confirmed the enrichment of POR in stalled D1-RNCs compared to uS2-RNCs using immunoblot analysis (Figure 4).

**Table 2.** Chloroplast proteins associated with D1 RNCs and functional in metabolic pathways Mass spectrometry analyses revealed the enrichment of chloroplast metabolic proteins with the RNCs of short (s), medium (m), and long (l) TST-D1 peptides in comparison to the TST-uS2c RNCs. Asterisks mark proteins that were exclusively found in the TST-D1 RNCs.

| Uniprot ID | Uniprot Name | Function | Razor Psat | Potential AT | enriched in D1 RNC |
| --- | --- | --- | --- | --- | --- |
| Q9FLW9 * | Plastidial pyruvate kinase 2 (PKP2) | Carbon metabolism | Psat1g096960.1<br>Psat6g194760.1 | AT5G52920.1 | s, m, l |
| Q43117 | Pyruvate kinase isozyme A (KPYA) | Carbon metabolism | Psat6g113960.1 | AT3G22960.1 | s, m, l |
| Q9ZU52 * | Fructose-bisphosphate aldolase 3 (ALFP3) | Carbon metabolism | Psat6g223200.1 | AT2G01140.1 | s, m, l |
| Q9SN86 * | Malate dehydrogenase (MDHP) | Carbon metabolism | Psat4g051000.1 | AT3G47520.1 | s, m, l |
| Q9SZX3 * | Argininosuccinate synthase (ASSY) | Amino acid metabolism | Psat0s3176g0120.1*<br>Psat4g065240.1 | AT4G24830.1 | s, m, l<br>m |
| P37142 | Bifunctional aspartokinase/homoserine dehydrogenase (AKH) | Amino acid metabolism | Psat7g075960.1 | AT4G19710.1 | s, m, l |
| Q9SW18 | Magnesium protoporphyrin IX methyltransferase (CHLM) | Chlorophyll metabolism | Psat7g039320.1 | AT4G25080.1 | s, m, l |
| Q01289 | Protochlorophyllide reductase (POR) | Chlorophyll metabolism | Psat5g245240.1 | AT5G54190 | s, m, l |
| Q8W250 * | 1-deoxy-D-xylulose 5-phosphate reductoisomerase (DXR) | Terpenoid metabolism | Psat4g058000.1 | AT5G62790.1 | s, m, l |
| P52418 * | Amidophosphoribosyltransferase (PUR1) | Purine metabolism | Psat7g181880.1 | AT2G16570.1 | s, m, l |

A large functional D1-RNCs associated group is formed by factors involved in protein processing like maturation, folding or degradation (Table 3). Notably, among these are two factors of D1 nascent peptide maturation: the peptide-deformylase DEF1B, which was exclusively found in short TST-D1 translation intermediates, and the subsequent acting methionine aminopeptidase MAP1B. Furthermore, the plastidic trigger factor (TIG1), a homolog of the bacterial trigger factor located close to the exit of the ribosomal polypeptide tunnel and acting as a molecular chaperone on emerging nascent chains, was exclusively enriched in the short TST-D1 RNCs (Table 3, Supplemental Table 4). However, immunoblot analysis indicates that TIG1 is also present in the middle chain length of TST-D1-RNCs and the TST-uS2-RNCs (Figure 4). Consistent with the mass spectrometry data these samples contained a similar amount of TIG1 (Figure 4, Supplemental Table 4). Our data suggest that TIG1 might preferentially bind to D1 translating ribosomes at early translation stages before exposure of the first TMH from the ribosomal peptide tunnel and indicate that the ribosome-associated protein biogenesis factors DEF1B, MAP1B, TIG1 and cpSRP54 can bind simultaneously to D1 translating ribosomes with nascent chain of up to about 69 amino acids. Moreover, members of the chloroplast chaperonin system, CPN60*α* and CPN60*α*, were enriched in all D1-chain length RNCs (Table 3). The cochaperonin CPN20 was also detected, but this finding was limited to the medium chain length samples. Chloroplast chaperonin CPN60 was originally described as a Rubisco-binding protein and its function assigned to a posttranslationally acting chaperone (Zhao and Liu, 2017). However, it has also been described that CPN60 and its bacterial counterpart, the GroEL/GroES system, bind ribosomes in a puromycin-dependent manner pointing towards a cotranslational function (Kramer et al., 2019; Ries et al., 2020; Westrich et al., 2021). Additional chaperones that were enriched in one or two chain length samples are members and cochaperones of the heat shock protein family and the 43 kDa subunit of the chloroplast SRP (cpSRP43) (Table 3). As factors that might be involved in degradation of an aberrantly folded nascent D1-chains we detected the protease CLPP5 to be associated with all D1 chain length samples. Additional proteases were detected in samples translating short and medium D1 variants (DEG2, PREP2 and CGEP) (Table 3).

**Table 3.** Chloroplast proteins associated with D1 RNCs and functional in protein processing List of proteins associated with TST-D1 RNCs and functionally categorized in protein processing. Asterisks mark proteins that were exclusively found in TST-D1 RNCs. Peptide chain length of TST-D1 RNCs: short (s), medium (m), long (l).

| Uniprot ID | Uniprot Name | Function | Razor Psat | Potential AT | enriched in D1 RNC |
| --- | --- | --- | --- | --- | --- |
| Q9FUZ2 * | Peptide deformylase 1B (DEF1B) | Deformylation, specificity to PS II D1 polypeptide | Psat3g166600.1 | AT5G14660.1 | s |
| Q9FV52 | Methionine aminopeptidase 1B (MAP1B) | N-terminal methionine excision | Psat2g085800.1 | AT1G13270.1 | s, l |
| Q8S9L5 | Trigger factor-like protein TIG | Chaperone | Psat2g009800.1 | AT5G55220.1 | s |
| P08926 | 60 kDa chaperonin subunit alpha (CPN60A) | Chaperone | Psat7g144320.1 | AT2G28000.1 | s, m, l |
| P08927 | 60 kDa chaperonin subunit beta (CPN60B) | Chaperone | Psat1g001680.1<br>Psat6g016920.1 | AT1G55490 | s, m, l<br>s, m, l |
| O65282 | 20 kDa chaperonin (CH20) | Chaperone | Psat1g135800.1 | AT5G20720.1 | m |
| Q9SIF2 | Heat shock protein 90-5 (HS905) | Chaperone | Psat2g102720.1 | AT2G04030.1 | s, l |
| Q9LF37 * | Chaperone protein ClpB3 | Chaperone | Psat1g069480.1 | AT5G15450.1 | s, m |
| G7L2M8 | GrpE protein homolog | Chaperone | Psat3g085320.3 | AT5G17710.1 | s |
| O22265 | Signal recognition particle 43 kDa protein (cpSRP43) | Chaperone | Psat7g178840.1 | AT2G47450.1 | s, m |
| Q9S834 | Clp protease proteolytic subunit 5 (CLPP5) | Protease | Psat6g161200.1 | AT1G02560.1 | s, m, l |
| O82261 * | Protease Do-like 2 (DEGP2) | Protease, primary cleavage of photodamaged D1 protein | Psat6g167960.1 | AT2G47940.1 | s, m |
| Q8VY06 * | Presequence protease 2 (PREP2) | Protease | Psat4g213560.1 | AT1G49630.1 | s, m |
| Q8VZF3 * | Probable glutamyl endopeptidase (CGEP) | Protease | Psat3g019800.3 | AT2G47390.1 | s |
| O64730 * | Probable protein phosphatase 2C 26 (P2C26) | PSII phosphatase | Psat5g153360.1 | AT2G30170.1 | s, m, l |

### STIC2 is partially associated with a stromal higher molecular weight complex and with thylakoid membranes

As described above, we identified STIC2 as a factor specifically associated with TST-D1 RNCs. To analyze the association of STIC2 with stromal ribosomes a stromal extract of pea chloroplasts was applied to size exclusion chromatography and the corresponding fractions were subjected to immunoblot analyses using antibodies against STIC2 and the ribosomal subunit uL4. While a markedly different elution behavior was observed between ribosomes and most of the endogenous STIC2, a small amount of STIC2 coeluted together with fractions containing ribosomes (Figure 5A). As ribosomes prepared from a stromal extract are probably largely translationally inactive or predominantly translate soluble proteins, these data are consistent with our finding that STIC2 is only recruited to ribosomes engaged in translating D1 or possibly other thylakoid membrane proteins.

**Figure 5.**
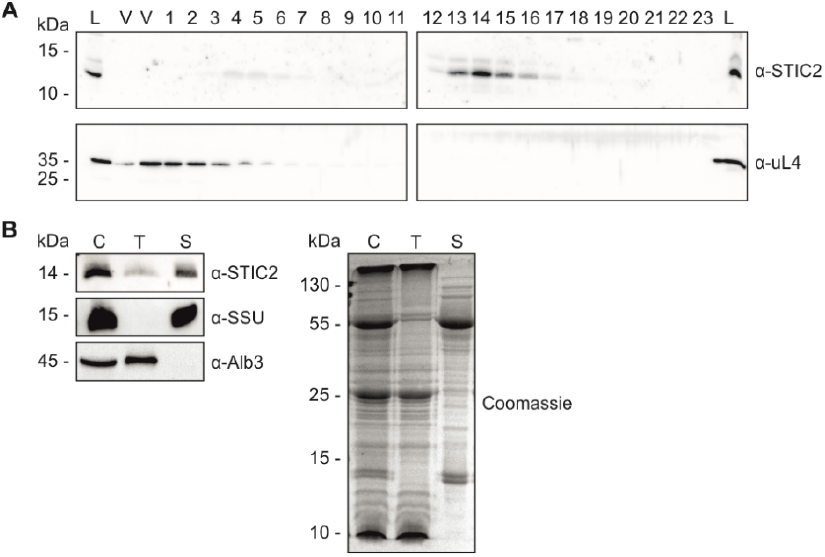
Analyses of STIC2 interaction with stromal ribosomes and thylakoid membranes (**A**) The interaction of endogenous STIC2 and ribosomes was tested by size exclusion chromatography (SEC) of chloroplast stroma from *Pisum sativum*. A stromal loading samples (L) was taken prior to SEC. The SEC fractions were separated by SDS-PAGE and applied to Western blot. Immunodetection was performed with α-STIC2 and α-uL4 antibodies. V: void volume. (B) Isolated chloroplasts from Arabidopsis thaliana (Col0) were lysed and separated into stromal and thylakoid fractions by centrifugation. Samples equivalent to 5 µg chlorophyll of chloroplast extract (C), washed thylakoids (T), and stroma (S) were separated on SDS-PAGE and blotted for immunodetection using specific antisera raised against STIC2, Alb3 and the small subunit of Rubisco (SSU). The Alb3 insertase and SSU were used as controls for successful fractionation. A Coomassie blue stained SDS gel served as loading control.

STIC2 was described to be exclusively located in the chloroplast stroma (Bédard et al., 2017). However, prior chloroplast proteome studies (Peltier et al., 2004), the ability of STIC2 to interact with Alb3 and Alb4 (Bédard et al., 2017) and our finding that STIC2 is a D1-RNC associated factor point to an at least partial or transient association with the thylakoids. To reexamine the subchloroplast localization of STIC2, *Arabidopsis* chloroplasts were fractionated into stroma and thylakoids. In the subsequent immunoblot, STIC2 was detected predominantly in the stromal fraction. However, a significant amount of the protein was also present in the thylakoid fraction (Figure 5B) suggesting that thylakoid membrane-associated STIC2 functions in cotranslational insertion at the thylakoid membrane.

### The C-terminal motif III in Alb3 and Alb4 is crucial for the interaction with STIC2

To get further insight into the molecular function of STIC2, we examined its interaction with Alb3 and Alb4. Both thylakoid membrane proteins functionally contribute to the folding, insertion and assembly of thylakoid membrane proteins (Benz et al., 2009; Wang and Dalbey, 2011; Trösch et al., 2015b) and are characterized by a conserved hydrophobic core region comprising five TMHs and a stromal exposed positively charged extension at their C-termini (Hennon et al., 2015; Ackermann et al., 2021; McDowell et al., 2021). We employed pepspot analyses to characterize the binding interfaces. For mature Alb3 (aa 56-462), the designed pepspots covered the first stromal loop (C1) (aa 155-208) and transmembrane domains (TMH 3-5), stromal loop C2, luminal loop L2, and the whole C-terminus within residues 282-462 in 15mer peptides with an overlap of 11 aa. Corresponding homologous structures of Alb4 were covered with equally designed pepspots ranging from aa 139-192 and aa 266-499 (Figure 6 and Supplemental Figure 2). Incubation with recombinant His-mSTIC2 and subsequent detection with His-tag specific antibodies revealed that STIC2 interacted with one peptide spot of Alb3 (aa 386-400) and one peptide spot of Alb4 (aa 394-408). These two peptides largely match to motif III, a conserved sequence within the C-terminal regions of both proteins (Falk et al., 2010).

**Figure 6.**
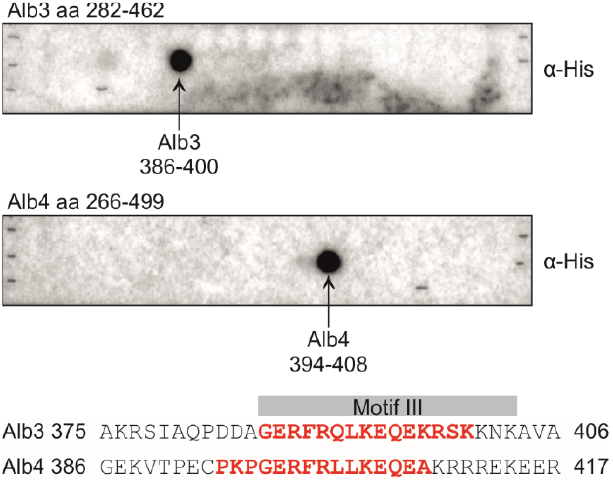
Recombinant STIC2 binds C-terminal motif III in Alb3 and Alb4 pepspot analyses The interaction of His-mSTIC2 with Alb3 (aa 282-462, upper panel) and Alb4 (aa 266-499, lower panel) was analyzed using pepspot-labeled nitrocellulose membranes. Recombinant His-mSTIC2 was incubated in a final concentration of 5 µg/ml with the pepspot membranes. Bound His-mSTIC2 was detected with antisera directed against the His-tag. Detected spots correspond to amino acids 386-400 of Alb3 and amino acids 394-408 of Alb4. These residues are indicated in red in an alignment of a C-terminal region of Alb3 and Alb4 comprising the conserved motif III (grey box).

*In vitro* pulldown assays using the recombinant His-tagged C-terminus of Alb3 (Alb3C-His; aa 350-462) or Alb4 (Alb4C-His; aa 334-499) or the corresponding motif III deletion constructs together with GST-mSTIC2 confirmed the role of motif III in Alb3 and Alb4 binding (Figure 7, Supplemental Figure 3).

**Figure 7.**
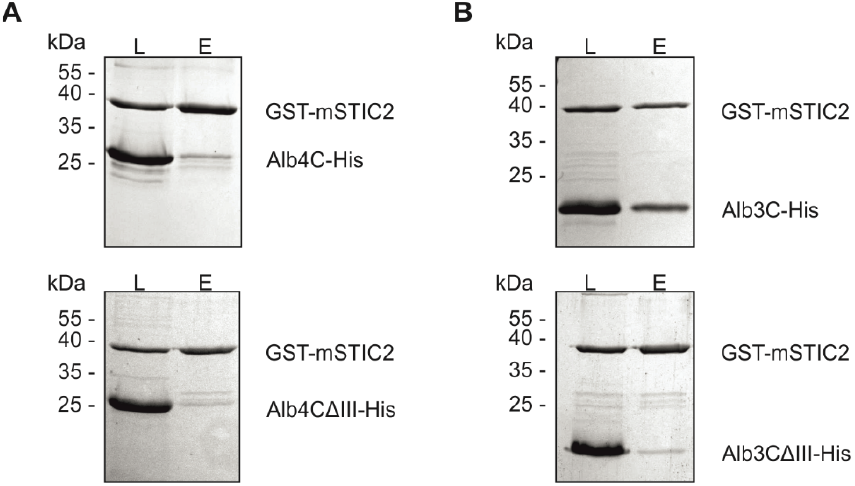
The conserved C-terminal motif III of Alb3 and Alb4 is crucial for the physical interaction with STIC2 In vitro pulldown experiments were performed with equal amounts of GST-tagged mSTIC2 and His-tagged Alb4 and Alb3 variants using glutathione resin. Loading (L) and eluate (E) samples were analyzed by SDS-PAGE and Coomassie blue staining. (**A**) Alb4C-His, C-terminal region of Alb4 comprising amino acids 334-499 (upper panel); Alb4CΔIII-His, Alb4C-His lacking motif III (Δ397-414) (lower panel). (**B**) Alb3C-His, C-terminal region of Alb3 comprising amino acids 350-462 (upper panel); Alb3CΔIII-His, Alb3C-His lacking motif III (Δ386-403) (lower panel).

To quantitatively analyze the interactions of STIC2 with Alb3C-His and Alb4C-His the binding reactions were assessed using isothermal titration calorimetry (ITC). Titrations of His-mSTIC2 into Alb3C-His resulted in an average dissociation constant of 135.8 µM (± 29.3 µM) (Figure 8A). An average dissociation constant of 16.6 µM (± 3.9 µM) was determined for His-mSTIC2 and Alb4C-His (Figure 8C). The constructs Alb3CΔIII-His and Alb4CΔIII-His showed no binding to STIC2 (Figure 8B and D).

**Figure 8.**
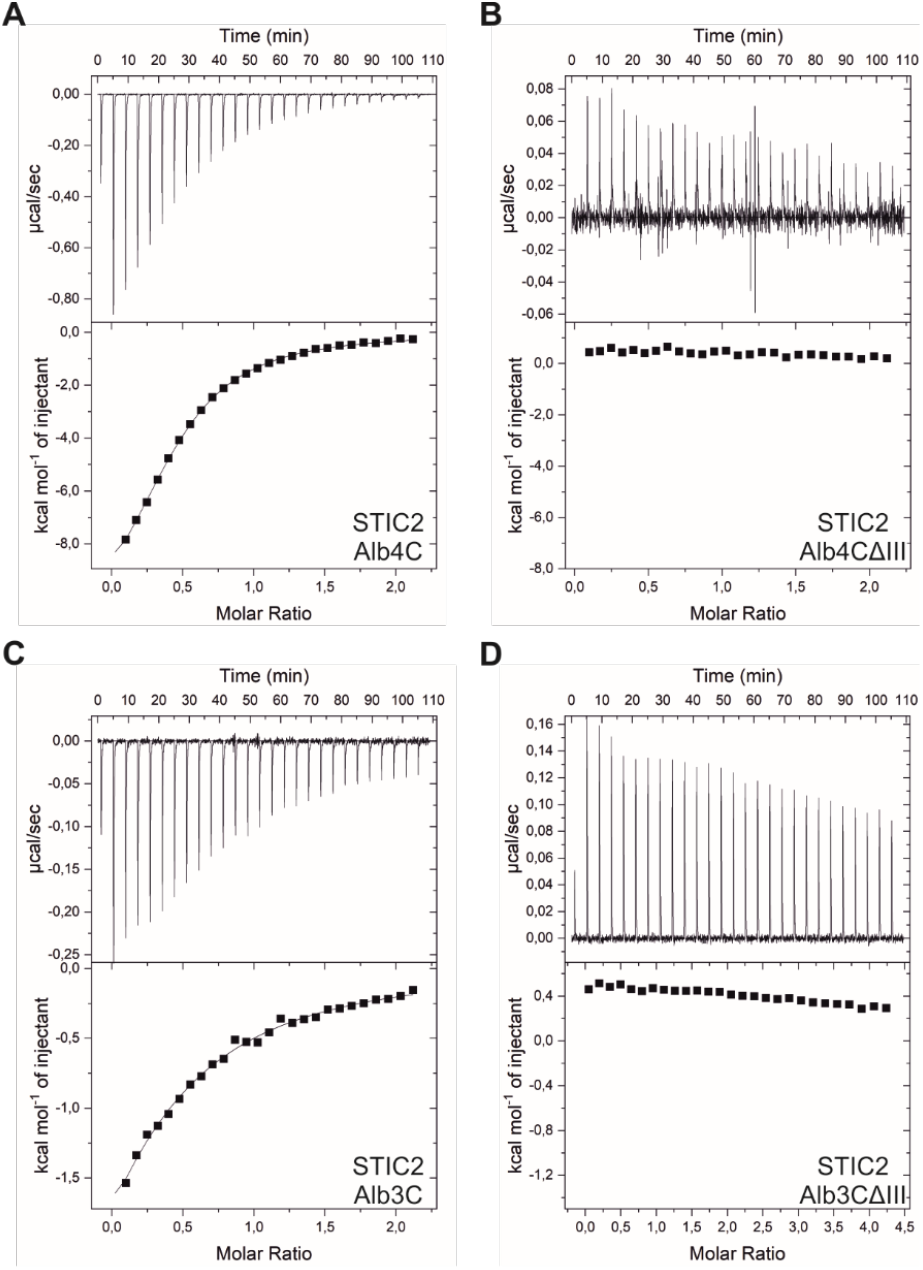
Conserved motif III is crucial for STIC2 binding in isothermal titration calorimetry Representative measurements of isothermal titration calorimetry (ITC) to determine binding affinities for the interaction of STIC2 with the C-terminal regions of Alb3 and Alb4. A 1.5 mM His-mSTIC2 solution was titrated into 0.15 mM of (**A**) Alb3C-His (350-462) or (**B**) Alb3CΔIII-His (Δ386-403). Analysis of titration isotherms from two biological replicates resulted in an average Kd of 135.8 ± 29.3 µM for the STIC2/Alb3C interaction. No interaction was observed for STIC2/Alb3CΔIII. The other titrations were performed with 1.0 mM His-mSTIC2 and 0.1 mM (**C**) Alb4C-His (334-499) or (**D**) Alb4CΔIII-His (Δ397-414) solutions. The titration isotherms of two biological replicates resulted in a Kd of 16.6 ± 3.9 µM for the STIC2/Alb4C interaction. No interaction was measured for STIC2/Alb4CΔIII.

## Discussion

In this work, we affinity-purified stalled *in vitro* translating ribosomes via twin-strep-tagged nascent peptides of the photosystem II reaction center protein D1 and ribosomal uS2c. This method allows for the first time the purification of a homogeneous chloroplast ribosome population and thus the analysis of the interaction network of ribosomes specifically translating a thylakoid membrane protein. Mass spectrometric analyses of the purified RNCs showed that a broad variety of proteins from metabolic, biogenesis, and protein processing pathways associate with D1-translating ribosomes in the chloroplast.

In accordance with rising evidence for a linkage between the synthesis of chlorophyll and proteins in chloroplasts (Westrich et al., 2021; Trösch et al., 2022), we observed the accumulation of chlorophyll biosynthesis enzymes such as the magnesium protoporphyrin IX methyltransferase (CHLM) and protochlorophyllide reductase (POR) in D1 RNCs. A well-coordinated expression of chlorophyll-binding apoproteins and chlorophyll synthesis is required in the chloroplast to avoid the formation of harmful reactive oxygen species and phototoxic stress due to the accumulation of free, highly photoreactive chlorophyll molecules (Apel and Hirt, 2004; Wang and Grimm, 2015). Furthermore, we identified several metabolic enzymes of e.g. the carbon or amino acid metabolism in the D1-RNC interactome. As it has been shown that ribosomes play an important role in the control of metabolic acclimation (Garcia-Molina et al., 2020) our data suggest that the coordination occurs, at least in part, directly at RNC complexes.

An interesting link between D1 synthesis and thylakoid membrane biogenesis arises from the association of the fuzzy onions-like (FZL) chloroplast membrane-remolding GTPase to D1 RNCs. Notably, this link was previously suggested, as during early steps of thylakoid biogenesis accumulation of FZL at the thylakoids is accompanied by increased thylakoid binding of RNCs and insertion of plastid-encoded PSII subunits (Liang et al., 2018). FZL was also identified in the ribosomal interaction network in *Chlamydomonas reinhardtii* chloroplasts (Westrich et al., 2021) and was described in the regulation and determination of thylakoid and chloroplast morphology in plants (Gao et al., 2006).

Previous studies concluded that the transcriptional and translational system might be linked in chloroplasts (Pfalz et al., 2006; Zoschke and Bock, 2018). In support of these data, we observed an association of fructokinase-like 1 (SCKL1/FLN1) in all D1 chain length samples. As part of the plastid-encoded RNA polymerase, FLN1 participates in the transcription of photosynthesis-related genes (Shiina et al., 2005; Börner et al., 2015).

In chloroplasts a diverse protein network is employed in protein processing and quality control (Trösch et al., 2015a; Breiman et al., 2016; Ries et al., 2020; Westrich et al., 2021). Consistent with this, subunits of the N-terminal methionine excision machinery, TIG1, chaperonin, heat shock protein, co-chaperones as well as components of the chloroplast caseinolytic protease (Clp) system and other proteases were enriched in D1 RNCs, and therefore likely contribute to cotranslational targeting of chloroplast proteins. However, the exact molecular function of many of these factors in cotranslational folding or quality control remains open and needs to be addressed in future studies.

In this work, we identified STIC2 as interactor of RNCs translating D1, while we did not observe any interaction of STIC2 with RNCs of the plastid-encoded uS2c. We therefore suggest that STIC2 is specifically recruited to stromal ribosomes translating D1. STIC2 homologues are widely distributed in bacteria (YbaB/EbfC proteins) (Cooley et al., 2009) but can also be found in streptophytes, chlorophytes, and cyanobacteria. Interestingly, bacterial YbaB was suggested to be involved in a pathway that facilitates the folding, assembly, and quality control of inner membrane proteins in *E. coli* (Skretas and Georgiou, 2010). In line with our data, STIC2 was recently described to participate in cpSRP-dependent thylakoid biogenesis based on the phenotypic impairments in *stic2/ffc* and *stic2/alb4 Arabidopsis* mutants (Bédard et al., 2017). Furthermore, a direct interaction of STIC2 with both thylakoid membrane proteins, Alb3 and Alb4, was reported (Bédard et al., 2017). We confirmed these interactions and provide data which demonstrate that the stroma-exposed conserved motif III within the C-terminal region (Falk et al., 2010) of Alb3 and Alb4 is crucial for the STIC2 interaction. The C-terminal region of the Alb3 insertase is also involved in the interaction with ribosomes (Ackermann et al., 2021) and contributes to binding of cpSRP54 in complex with its receptor, cpFtsY (Chandrasekar and Shan, 2017). We speculate that STIC2 generates a link between the D1 RNCs and the insertase machinery and might be involved in the spatiotemporal coordination of the translocon-bound RNC at the thylakoid membrane. However, further studies are required to investigate the molecular details of the STIC2 interaction with the D1 RNC complexes and the membrane insertase.

## Materials and methods

### Plant material and growth conditions

*Arabidopsis thaliana* (Col0) and *Pisum sativum* cv (Wonder of Kelvedon) were grown on soil under artificial light (8 h light, 120 µmol photons/m^2^ • s^1^, 22°C 65 % humidity, 16 h dark, 19,5°C, 65 % humidity).

### Overexpression and purification of recombinant proteins

For the expression of recombinant protein, a cDNA clone coding for *Arabidopsis thaliana* STIC2 (clone number U13069) was obtained from Arabidopsis Biological Resource Center (ABRC). Considering the cTP prediction of TargetP, the coding sequence for the mature STIC2 protein (aa 49-182) was cloned into BamHI/SalI linearized pETDuet1 (Novagen) and pGEX4T3 (Cytiva) using the In-Fusion^®^ HD EcoDry^™^ kit (Takara) and: for pETDuet1: CATCACCATCATCACCACAGCCAGGTGAATGGATTATTTGGAGGTGGAA rev pETDuet1: GCGGCCGCAAGCTTGTCGACTTACTTCATTCCTTCGCTGA for pGEX4T3: ATCTGGTTCCGCGTGGATCCGTGAATGGATTATTTGG rev pGEX4T3: GGCCGCTCGAGTCGACTTACTTCATTCCTTCGCTGA Constructs for expression of soluble C-terminal proteins of Alb3 (Alb3C: aa 350-462; Alb3CΔIII: lacking aa 386-403) and Alb4 (Alb4C: aa 334-499; Alb4CΔIII: lacking aa 397-414) were cloned as described before (Bals et al., 2010; Dünschede et al., 2011; Dünschede et al., 2015). Recombinant proteins with His-tag or GST-tag were expressed in *Escherichia coli* Rosetta2 (DE3) or BL21 (DE3) cells. The purification of proteins was carried out with HisTrap HP prepacked columns or glutathione sepharose (Cytiva) under native conditions. After purification, proteins for pulldown assays were stored in PBS buffer (300 mM sodium chloride, 2.7 mM potassium chloride, 10 mM disodium hydrogen phosphate, 1.8 mM potassium dihydrogen phosphate, pH 8.0). Otherwise, proteins were stored in HEPES buffer (20 mM HEPES, 300 mM sodium chloride, pH 8.0).

### Isolation of chloroplasts for translational active S30 extract

The isolation of intact chloroplasts from 8 day old pea leafs was performed according to Walter et al. (2015). The preparation of translationally active S30 extract was carried out as described in Yukawa et al. (2007).

### In vitro translation and RNC isolation

DNA templates of twin-strep-tag (TST) modified D1 and uS2c proteins were obtained from Invitrogen (GeneArt Gene Synthesis). Therefore, T7 promotor sequence and sequential full length CDS of either *psbA* from *Pisum sativum*, including the 5’ UTR (from -87 to -1), or *rps2* from *Arabidopsis thaliana* (5’ UTR: from -90 to -1) with internal TST were provided in pMA vector backbone. The TST CDS was inserted at position +76 (*psbA*) between E25 and N26 of the D1 protein or between G17 and V18 of uS2c at position +52 (*rps2*). The TST CDS was additionally adapted to pea chloroplast codon usage according to the Kazusa codon usage database (www.kazusa.or.jp/codon/): 5’-TCTGCTTGGTCCCATCCTCAATTTGAAAAAGGTGGTGGTTCTGGTGGTGGTAGTGGTGGATCTGCTTGGAGTCATCCACAATTCGAGAAA-3’ PCR with PRECISOR High-Fidelity DNA-polymerase (BioCat) was utilized to amplify truncated DNA templates of different length determined by reverse primer and lacking a stop codon: for. TST-D1: GTAATACGACTCACTATAGGGCGAGTAACAAGCCCTTAATTCTATAGTTA rev. TST-D1 (1-56): AGGGGCAGCAATGAAAGCG; rev. TST-D1 (1-69): TCCAGAAACAGGCTCACGAATACC; rev. TST-D1 (1-108): CGTTGTATAACCATTCATCAACGGATGC; rev. TST-D1 (1-136): GACGAAAACTAAGTTCCCACTCACG; rev. TST-D1 (1-195): GGTGCATAAGAATATTATGCTCAGCC; rev. TST-D1 (1-291): GCTGATACCTAACGCGGTAAACCAG; for TST-uS2c: TAATACGACTCACTATAGGG; rev TST-uS2c (1-158): CATATATTTAATCCCGCCTAGATATG; The truncated PCR products were used for *in vitro* transcription with the TranscriptAid T7 high yield transcription kit (Thermo Scientific) and transcripts were purified with MEGAclear(tm) transcription clean-up kit (Invitrogen).

According to the experimental procedure previously described by Yukawa et al. (2007) and modified by Walter et al. (2015), *in vitro* translation was run for 60 min at 28°C and illumination in 40 µl reactions. Chloramphenicol was added (final concentration 0.85 mg/ml) to stop the reaction and RNCs were subsequently enriched via sucrose cushion centrifugation or isolated in affinity-based purification using the streptactin-XT System.

Sucrose cushion centrifugation was utilized to enrich RNCs from the translation reactions. After translation, 160 µl total reaction were loaded on 2.9 ml sucrose cushion (1 M sucrose, 30 mM HEPES pH 7.7, 9 mM magnesium acetate, 70 mM potassium acetate, 2 µg/ml antipain, 2 µg/ml leupeptin, 5 mM DTT, 0.25 mg/ml chloramphenicol, 0.1 mM AEBSF) and centrifuged (270000 g, 90 min, 4°C, TLA 100.3, Beckman Coulter). The pellet was solubilized in sample buffer and subsequently applied to SDS-PAGE and Western blot analyses.

For mass spectrometric analyses RNCs were isolated with MagStrep XT magnetic beads (IBA Lifescience). Therefor, 100 µl bead suspension was equilibrated with wash buffer (30 mM HEPES pH 7.7, 200 mM sodium chloride, 70 mM potassium acetate, 9 mM magnesium acetate, 5 mM DTT) containing 0.005 % Tween20 in a magnetic separator. A total of 160 µl translation reaction was incubated with equilibrated beads for 30 min (4°C, rotation). The supernatant was removed in the separator and the beads were washed twice with wash buffer containing 0.005 % Tween20 and twice with wash buffer. The RNCs were eluted with BTX buffer (IBA Lifescience) (1xBTX, 70 mM potassium acetate, 10 mM magnesium acetate, 5 mM DTT) for 20 min (4°C, rotation) and collected from the beads in the magnetic separator. Isolation of RNCs for Western blot analyses was performed using 1 ml prepacked StrepTrap XT columns and the ÄKTApurifier (Cytiva). The columns were loaded with 320 µl-880 µl translation reaction (0.1 ml/min) and washed (0.5 ml/min) with 10 column volumes (CV) wash buffer containing 0.005 % Tween20. Elution was done with 10 CV BTX buffer and collected in 0.5 ml fractions. Protein-containing fractions were then pooled, concentrated (Amicon Ultra-0.5, Merck), and subsequently equal amounts of protein according to OD_280_ were applied to SDS-PAGE and Western blot analyses.

For SDS-PAGE and subsequent autoradiography analyses, the 40 µl *in vitro* translation reactions contained 20 µM 19 amino acid mix (Promega) and 0.45 µCi/µl of [35S]-methionine (Hartmann Analytics). The isolation of RNCs with MagStrep XT magnetic beads was essentially performed as described above with 30 µl of beads suspension.

### Mass spectrometry analyses

For tandem mass spectrometry (MS/MS), samples were prepared as described before (Kraus et al., 2020) using an in-gel tryptic digest. In brief, *in vitro* translation reactions were affinity-purified as described and the eluates were run on a SDS-PAGE until reaching the separation gel to remove impurities and cut out as a whole. The gel pieces of the different samples were washed and digested over night at 37 °C with MS-grade trypsin (Promega). MS/MS analysis was done using a previously described setup of a Waters nanoACQUITY gradient UPLC (www.waters.com) coupled to a Thermo Fisher Scientific Inc. Orbitrap ELITE mass spectrometer using a gradient over 60 minutes (Cormann et al., 2016; Kraus et al., 2020).

Data analysis was done with MaxQuant 1.6.10.43 (Cox and Mann, 2008) using a *Pisum sativum* database from 16.01.2019 of the “*Unité de Recherches en Génomique Info*” (URGI, UR1164, https://urgi.versailles.inra.fr/Species/Pisum) with additional annotations of candidates for potential homologues (Kreplak et al., 2019; https://www.pulsedb.org: Ps_Cameor_v1a-proteins_sprot_091019). Additionally, the database was extended by the contaminant list of MaxQuant. For qualitative analysis, the twin-strep-tagged sequences of TST-D1 and TST-uS2c were added to the database. These have to be removed for quantitative analysis using label-free quantification (LFQ, Supplemental Table 4) due to the twin-strep-tag sequence redundancy. The precursor mass tolerance was set to 20 p.p.m. As modifications, methionine oxidation and N-terminal acetylation were included. The FDR of protein identification was set to 1%. A maximum of two missed cleavage sites was allowed.

Two independent preparations for each of TST-D1 RNC samples with different nascent peptide lengths (56-291) and the TST-uS2c (158) RNC sample, respectively, were analyzed in technical duplicates by MS/MS. To identify enriched proteins, the label-free quantification intensities (LFQs) of each TST-D1 sample were compared with the LFQs of the TST-uS2c sample, and statistically significant enrichment was determined by two-sided unpaired *t*-test (*p*-value ≤ 0.05). On this basis, significantly enriched and D1-exclusively assigned proteins of TST-D1 (56) and TST-D1 (69) samples were grouped into a *short* RNC peptide category. Accordingly, the proteins of the TST-D1 (108) and TST-D1 (136) samples, and the TST-D1 (195) and TST-D1 (291) samples were categorized into *medium* and *long* groups, respectively. These categories are based on the exposure of transmembrane helices (TMHs) of the nascent D1 peptide, with the first TMH still fully buried inside the ribosome peptide tunnel for *short* D1 RNCs, an exposed first TMH in *medium* D1 RNCs, and at least two exposed TMHs in *long* D1 RNCs.

Proteins assigned to D1 and uS2c RNCs were depicted in Volcano plots. The ratios of averaged D1 over uS2c LFQs (log_2_) were plotted against *p*-values (–log_10_). A grey line represents the *p*-value threshold (≤ 0.05).

### Arabidopsis chloroplast isolation and fractionation

Chloroplast isolation and fractionation was previously described in Hristou et al. (2019). *Arabidopsis thaliana* plants, 3 to 4 week old, were homogenized in isolation buffer (300 mM sorbitol, 5 mM magnesium chloride, 5 mM EGTA, 5 mM EDTA, 10 mM sodium bicarbonate, 20 mM HEPES, pH 7.7) and filtered through Miracloth. The chloroplasts were pelleted (5 min, 1000 g, 4°C) and loaded on preformed Percoll gradients (50 % Percoll, 0.5 mM reduced L-glutathione in isolation buffer, 30 min, 40,000 g, 4°C, brake off). Intact chloroplasts were separated by centrifugation at 7800 g (4°C, 10 min, brake off) and could be collected from the lower green band. Afterwards, the chloroplasts were washed in 300 mM sorbitol, 3 mM magnesium chloride, and 50 mM HEPES pH 8.0 (1000 g, 4°C, 5 min). Lyses of the chloroplasts was carried out at 1 to 2 mg chlorophyll/ml in HM buffer (10 mM magnesium chloride, 50 mM HEPES pH 8.0), and thylakoids were then separated from stroma by centrifugation (14,000 rpm, 4°C, 10 min). Thylakoids were finally washed in HM buffer containing 150 mM sodium chloride and then adjusted to the initial chlorophyll concentration of chloroplasts with HM buffer. Gels for SDS-PAGE were loaded on a proportional chlorophyll basis (5 µg) with whole chloroplasts, stroma and thylakoids and subsequently applied to Coomassie^®^ *Brilliant Blue* R-250 staining or Western blot analyses.

### Isolation of Pisum sativum chloroplasts

Leafs from 9 day old pea plants (25 g of fresh weight) were grinded in 200 ml of 330 mM sorbitol, 50 mM HEPES, 3 mM magnesium chloride, 0.05 % BSA, and 5 mM ascorbic acid, pH 8.0 (grinding buffer). The homogenate was filtered through two layers of Miracloth and then centrifuged (4000 rpm, 4°C, 5 min). The supernatant was removed and the green pellet was resuspended in SHM buffer (330 mM sorbitol, 50 mM HEPES, 3 mM magnesium chloride, pH 8.0) and loaded onto Percoll gradients. For the gradients, 8 ml 80 % Percoll solution was layered under 12 ml 40 % Percoll solution (in 330 mM sorbitol, 50 mM HEPES, pH 8.0). By centrifugation at 7700 g (10 min, 4°C), intact chloroplasts were separated from disrupted chloroplasts and accumulated in a lower green band. The intact chloroplasts were collected and washed with SHM buffer (5000 rpm, 4 °C, 2 min). After that, the chloroplasts were adjusted to 2 mg/ml chlorophyll with either HM or LSB buffer (20 mM HEPES, 50 mM potassium acetate,6 mM magnesium acetate, 2 mM DTT, pH 7.5) for fractionation or isolation of ribosomes, respectively.

### Pea chloroplast fractionation and size exclusion chromatography

Isolated chloroplasts from pea plants were lysed at 2 mg chlorophyll/ml in HM buffer. Centrifugation (14,000 rpm, 4°C, 10 min) was then performed twice to separate stroma from thylakoids. Thylakoids were washed twice with HM buffer containing 150 mM NaCl. Size exclusion chromatography (SEC) with 500 µl pea stroma was conducted over a Superdex 200 column in HM buffer with 150 mM sodium chloride. SEC fractions were then subjected to Western blot analysis.

### Pepspot analyses

Peptide libraries were obtained from JPT Peptide Technologies GmbH, Berlin. For Alb3, peptide membranes with 11 and 43 spots (15mer peptides, overlapping by 11 amino acids) covering amino acids 155-208 and 282-462 were designed. Alb4 peptide membranes, with 11 and 56 spots (15mer, overlap of 12 amino acids), comprised amino acids 139-192 and 266-499.

According to manufactures instructions, the membranes were first incubated in methanol and then washed with TBST buffer (50 mM Tris-HCl, 150 mM sodium chloride, 0.3 % Tween 20, pH 7.5). Blocking of the membranes was performed in anti His HRP conjugate blocking buffer (Qiagen). After blocking at room temperature for 2 h, His-mSTIC2 (5 µg/ml) was added and incubated for another 3 h. After washing with TBST, bound His-mSTIC2 protein was detected by anti His HRP conjugate (Qiagen) and ECL reaction (Pierce).

### Pulldown analyses

For GST-pulldown experiments, 20 µg of His-tagged and GST-tagged proteins were incubated in PBS buffer with a final volume of 200 µl (10 min, rotation). A 10 µl load sample was taken and mixed with sample buffer. After that, 100 µl of equilibrated glutathione sepharose (Cytiva) was added to the pulldown reaction and incubated for another 30 min. Next, the pulldown reaction was loaded on a Wizard^®^ Minicolumn (Promega) and the supernatant was removed by short spinning. Unbound proteins were then washed out with 5 ml PBS buffer and the remaining supernatant was removed by centrifugation (14,000 rpm, 30 sec). For elution, 30 µl PBS buffer with 10 mM reduced L-glutathione was added to the column and incubated for 15 min. Eluted proteins were collected by centrifugation (14,000 rpm, 30 sec) and mixed with 15 µl sample buffer. Subsequently, 15 µl of load and eluate samples were separated on SDS-PAGE and stained with Coomassie^®^ *Brilliant Blue* R-250.

### Western blot analyses

Proteins separated by SDS-PAGE were blotted onto nitrocellulose membranes. The transferred proteins were detected by specific antibodies against Alb3 (Bals et al., 2010), cpSRP54M (Walter et al., 2015), FLN1 (PhytoAB), FZL (PhytoAB), POR (Agrisera), uL4 (Agrisera), uL24 (PhytoAB), rbcs (Agrisera), and twin-strep-tag (IBA Lifescience). Antibody against TIG1 was a gift from F. Willmund (Ries et al., 2017). A polyclonal STIC2-directed antibody was generated with His-mSTIC2 protein (aa 49-182) in rabbit (Davids Biotechnology).

### Isothermal titration calorimetry

ITC was performed to determine binding affinities for the interaction of mSTIC2 with Alb3C (aa 350-462), Alb4C (aa 334-499), or these proteins lacking the conserved motif III Alb3CΔIII (Δ386-403) and Alb4CΔIII (Δ397-414), respectively. The experiments were carried out in 20 mM HEPES buffer (150 mM sodium chloride, pH 8.0) at 25°C using an Auto-iTC200 (MicroCal). STIC2 protein with a tenfold concentration (1.0 mM or 1.5 mM) was titrated into solutions of 0.1 mM Alb4 or 0.15 mM Alb3 proteins, respectively. Data analysis and fitting was carried out with OriginPro 7 using a one set of sites model.

## Acknowledgements

We like to acknowledge Silke Funke for excellent technical assistance. We thank Felix Willmund for the kind gift of antibodies against TIG1. This work was supported by the Deutsche Forschungsgemeinschaft (SCHU 1163/6-2 and 836/3-2 within research unit FOR2092 to D.S. and M.M.N., respectively; BA 1902/2-2 to S.B.)

## References

Ackermann, B., Dünschede, B., Pietzenuk, B., Justesen, B.H., Krämer, U., Hofmann, E., Günther Pomorski, T., and Schünemann, D. (2021). Chloroplast Ribosomes Interact With the Insertase Alb3 in the Thylakoid Membrane. Front. Plant Sci. 12.

Akopian, D., Shen, K., Zhang, X., and Shan, S.-O. (2013). Signal recognition particle: an essential protein-targeting machine. Annual review of biochemistry 82: 693–721.

Apel, K., and Hirt, H. (2004). Reactive oxygen species: metabolism, oxidative stress, and signal transduction. Annu. Rev. Plant Biol. 55 (1): 373–399.

Bals, T., Dünschede, B., Funke, S., and Schunemann, D. (2010). Interplay between the cpSRP pathway components, the substrate LHCP and the translocase Alb3: an in vivo and in vitro study. FEBS letters 584 (19): 4138–4144.

Ban, N., Beckmann, R., Cate, J.H.D., Dinman, J.D., Dragon, F., Ellis, S.R., Lafontaine, D.L.J., Lindahl, L., Liljas, A., Lipton, J.M., McAlear, M.A., Moore, P.B., Noller, H.F., Ortega, J., Panse, V.G., Ramakrishnan, V., Spahn, C.M.T., Steitz, T.A., Tchorzewski, M., Tollervey, D., Warren, A.J., Williamson, J.R., Wilson, D., Yonath, A., and Yusupov, M. (2014). A new system for naming ribosomal proteins. Current opinion in structural biology 24: 165–169.

Bédard, J., Trösch, R., Wu, F., Ling, Q., Flores-Pérez, Ú., Töpel, M., Nawaz, F., and Jarvis, P. (2017). Suppressors of the Chloroplast Protein Import Mutant tic40 Reveal a Genetic Link between Protein Import and Thylakoid Biogenesis. The Plant cell 29 (7): 1726–1747.

Benz, M., Bals, T., Gügel, I.L., Piotrowski, M., Kuhn, A., Schünemann, D., Soll, J., and Ankele, E. (2009). Alb4 of Arabidopsis promotes assembly and stabilization of a non chlorophyll-binding photosynthetic complex, the CF1CF0-ATP synthase. Molecular plant 2 (6): 1410–1424.

Bieri, P., Leibundgut, M., Saurer, M., Boehringer, D., and Ban, N. (2017). The complete structure of the chloroplast 70S ribosome in complex with translation factor pY. The EMBO journal 36 (4): 475–486.

Bohne, A.-V., Schwarz, C., Schottkowski, M., Lidschreiber, M., Piotrowski, M., Zerges, W., and Nickelsen, J. (2013). Reciprocal regulation of protein synthesis and carbon metabolism for thylakoid membrane biogenesis. PLoS Biol 11 (2): e1001482.

Börner, T., Aleynikova, A.Y., Zubo, Y.O., and Kusnetsov, V.V. (2015). Chloroplast RNA polymerases: Role in chloroplast biogenesis. Biochimica et biophysica acta 1847 (9): 761– 769.

Breiman, A., Fieulaine, S., Meinnel, T., and Giglione, C. (2016). The intriguing realm of protein biogenesis: Facing the green co-translational protein maturation networks. Biochimica et biophysica acta 1864 (5): 531–550.

Chandrasekar, S., and Shan, S.-O. (2017). Anionic Phospholipids and the Albino3 Translocase Activate Signal Recognition Particle-Receptor Interaction during Light-harvesting Chlorophyll a/b-binding Protein Targeting. The Journal of biological chemistry 292 (1): 397–406.

Cooley, A.E., Riley, S.P., Kral, K., Miller, M.C., DeMoll, E., Fried, M.G., and Stevenson, B. (2009). DNA-binding by Haemophilus influenzae and Escherichia coli YbaB, members of a widely-distributed bacterial protein family. BMC microbiology 9: 137.

Cormann, K.U., Möller, M., and Nowaczyk, M.M. (2016). Critical Assessment of Protein Cross-Linking and Molecular Docking: An Updated Model for the Interaction Between Photosystem II and Psb27. Front. Plant Sci. 7: 157.

Cox, J., and Mann, M. (2008). MaxQuant enables high peptide identification rates, individualized p.p.b.-range mass accuracies and proteome-wide protein quantification. Nature biotechnology 26 (12): 1367–1372.

Dünschede, B., Bals, T., Funke, S., and Schünemann, D. (2011). Interaction studies between the chloroplast signal recognition particle subunit cpSRP43 and the full-length translocase Alb3 reveal a membrane-embedded binding region in Alb3 protein. The Journal of biological chemistry 286 (40): 35187–35195.

Dünschede, B., Trager, C., Schröder, C.V., Ziehe, D., Walter, B., Funke, S., Hofmann, E., and Schünemann, D. (2015). Chloroplast SRP54 Was Recruited for Posttranslational Protein Transport via Complex Formation with Chloroplast SRP43 during Land Plant Evolution. The Journal of biological chemistry 290 (21): 13104–13114.

Falk, S., Ravaud, S., Koch, J., and Sinning, I. (2010). The C terminus of the Alb3 membrane insertase recruits cpSRP43 to the thylakoid membrane. The Journal of biological chemistry 285 (8): 5954–5962.

Fernandez, D.E. (2018). Two paths diverged in the stroma: targeting to dual SEC translocase systems in chloroplasts. Photosynthesis research 138 (3): 277–287.

Gao, H., Sage, T.L., and Osteryoung, K.W. (2006). FZL, an FZO-like protein in plants, is a determinant of thylakoid and chloroplast morphology. Proceedings of the National Academy of Sciences 103 (17): 6759–6764.

Garcia-Molina, A., Kleine, T., Schneider, K., Mühlhaus, T., Lehmann, M., and Leister, D. (2020). Translational Components Contribute to Acclimation Responses to High Light, Heat, and Cold in Arabidopsis. iScience 23 (7): 101331.

Gilkerson, J., Perez-Ruiz, J.M., Chory, J., and Callis, J. (2012). The plastid-localized pfkB-type carbohydrate kinases FRUCTOKINASE-LIKE 1 and 2 are essential for growth and development of Arabidopsis thaliana. BMC plant biology 12 (1): 102.

Hennon, S.W., Soman, R., Zhu, L., and Dalbey, R.E. (2015). YidC/Alb3/Oxa1 Family of Insertases. J. Biol. Chem. 290 (24): 14866–14874.

Hristou, A., Gerlach, I., Stolle, D.S., Neumann, J., Bischoff, A., Dünschede, B., Nowaczyk, M.M., Zoschke, R., and Schunemann, D. (2019). Ribosome-associated chloroplast SRP54 enables efficient co-translational membrane insertion of key photosynthetic proteins. Plant Cell 31 (11): 2734–2750.

Jagendorf, A.T., and Michaels, A. (1990). Rough thylakoids: Translation on photosynthetic membranes. Plant Science 71 (2): 137–145.

Kannangara, C.G., Vothknecht, U.C., Hansson, M., and Wettstein, D. von (1997). Magnesium chelatase: association with ribosomes and mutant complementation studies identify barley subunit Xantha-G as a functional counterpart of Rhodobacter subunit BchD. Mol Gen Genet 254 (1): 85–92.

Kim, J., Klein, P.G., and Mullet, J.E. (1991). Ribosomes pause at specific sites during synthesis of membrane-bound chloroplast reaction center protein D1. J. Biol. Chem. 266 (23): 14931–14938.

Kramer, G., Shiber, A., and Bukau, B. (2019). Mechanisms of Cotranslational Maturation of Newly Synthesized Proteins. Annu. Rev. Biochem. 88 (1): 337–364.

Kraus, A., Weskamp, M., Zierles, J., Balzer, M., Busch, R., Eisfeld, J., Lambertz, J., Nowaczyk, M.M., and Narberhaus, F. (2020). Arginine-Rich Small Proteins with a Domain of Unknown Function, DUF1127, Play a Role in Phosphate and Carbon Metabolism of Agrobacterium tumefaciens. Journal of bacteriology 202 (22).

Kreplak, J., Madoui, M.-A., Cápal, P., Novák, P., Labadie, K., Aubert, G., Bayer, P.E., Gali, K.K., Syme, R.A., Main, D., Klein, A., Bérard, A., Vrbová, I., Fournier, C., d’Agata, L., Belser, C., Berrabah, W., Toegelová, H., Milec, Z., Vrána, J., Lee, H., Kougbeadjo, A., Térézol, M., Huneau, C., Turo, C.J., Mohellibi, N., Neumann, P., Falque, M., Gallardo, K., McGee, R., Tar’an, B., Bendahmane, A., Aury, J.-M., Batley, J., Le Paslier, M.-C., Ellis, N., Warkentin, T.D., Coyne, C.J., Salse, J., Edwards, D., Lichtenzveig, J., Macas, J., Doležel, J., Wincker, P., and Burstin, J. (2019). A reference genome for pea provides insight into legume genome evolution. Nat Genet 51 (9): 1411–1422.

Liang, Z., Zhu, N., Mai, K.K., Liu, Z., Tzeng, D., Osteryoung, K.W., Zhong, S., Staehelin, L.A., and Kang, B.-H. (2018). Thylakoid-Bound Polysomes and a Dynamin-Related Protein, FZL, Mediate Critical Stages of the Linear Chloroplast Biogenesis Program in Greening Arabidopsis Cotyledons. Plant Cell 30 (7): 1476–1495.

McDowell, M.A., Heimes, M., and Sinning, I. (2021). Structural and molecular mechanisms for membrane protein biogenesis by the Oxa1 superfamily. Nature structural & molecular biology 28 (3): 234–239.

Nilsson, R., Brunner, J., Hoffman, N.E., and van Wijk, K.J. (1999). Interactions of ribosome nascent chain complexes of the chloroplast-encoded D1 thylakoid membrane protein with cpSRP54. The EMBO journal 18 (3): 733–742.

Nilsson, R., and van Wijk, K.J. (2002). Transient interaction of cpSRP54 with elongating nascent chains of the chloroplast-encoded D1 protein; ‘cpSRP54 caught in the act’. FEBS letters 524 (1-3): 127–133.

Nishimura, K., Kato, Y., and Sakamoto, W. (2017). Essentials of Proteolytic Machineries in Chloroplasts. Molecular plant 10 (1): 4–19.

Olinares, P.D.B., Ponnala, L., and van Wijk, K.J. (2010). Megadalton complexes in the chloroplast stroma of Arabidopsis thaliana characterized by size exclusion chromatography, mass spectrometry, and hierarchical clustering. Molecular & cellular proteomics MCP 9 (7): 1594–1615.

Peltier, J.-B., Ytterberg, A.J., Sun, Q., and van Wijk, K.J. (2004). New functions of the thylakoid membrane proteome of Arabidopsis thaliana revealed by a simple, fast, and versatile fractionation strategy. J. Biol. Chem. 279 (47): 49367–49383.

Pfalz, J., Liere, K., Kandlbinder, A., Dietz, K.-J., and Oelmüller, R. (2006). pTAC2, -6, and -12 are components of the transcriptionally active plastid chromosome that are required for plastid gene expression. Plant Cell 18 (1): 176–197.

Piskozub, M., Króliczewska, B., and Króliczewski, J. (2015). Ribosome nascent chain complexes of the chloroplast-encoded cytochrome b6 thylakoid membrane protein interact with cpSRP54 but not with cpSecY. Journal of bioenergetics and biomembranes 47 (3): 265–278.

Ries, F., Carius, Y., Rohr, M., Gries, K., Keller, S., Lancaster, C.R.D., and Willmund, F. (2017). Structural and molecular comparison of bacterial and eukaryotic trigger factors. Scientific reports 7: 10680.

Ries, F., Herkt, C., and Willmund, F. (2020). Co-Translational Protein Folding and Sorting in Chloroplasts. Plants 9 (2).

Röhl, T., and van Wijk, K.J. (2001). In vitro reconstitution of insertion and processing of cytochrome f in a homologous chloroplast translation system. J. Biol. Chem. 276 (38): 35465–35472.

Rohr, M., Ries, F., Herkt, C., Gotsmann, V.L., Westrich, L.D., Gries, K., Trösch, R., Christmann, J., Chaux-Jukic, F., Jung, M., Zimmer, D., Mühlhaus, T., Sommer, F., Schroda, M., Keller, S., Möhlmann, T., and Willmund, F. (2019). The Role of Plastidic Trigger Factor Serving Protein Biogenesis in Green Algae and Land Plants. Plant physiology 179 (3): 1093–1110.

Shiina, T., Tsunoyama, Y., Nakahira, Y., and Khan, M.S. (2005). Plastid RNA Polymerases, Promoters, and Transcription Regulators in Higher Plants. International Review of Cytology 244: 1–68.

Skretas, G., and Georgiou, G. (2010). Simple genetic selection protocol for isolation of overexpressed genes that enhance accumulation of membrane-integrated human G protein-coupled receptors in Escherichia coli. Applied and environmental microbiology 76 (17): 5852–5859.

Sun, J.-L., Li, J.-Y., Wang, M.-J., Song, Z.-T., and Liu, J.-X. (2021). Protein Quality Control in Plant Organelles: Current Progress and Future Perspectives. Molecular plant 14 (1): 95– 114.

Sun, Y., and Zerges, W. (2015). Translational regulation in chloroplasts for development and homeostasis. Biochimica et biophysica acta 1847 (9): 809–820.

Trösch, R., Mühlhaus, T., Schroda, M., and Willmund, F. (2015a). ATP-dependent molecular chaperones in plastids--More complex than expected. Biochimica et biophysica acta 1847 (9): 872–888.

Trösch, R., Ries, F., Westrich, L.D., Gao, Y., Herkt, C., Hoppstädter, J., Heck-Roth, J., Mustas, M., Scheuring, D., Choquet, Y., Räschle, M., Zoschke, R., and Willmund, F. (2022). Fast and global reorganization of the chloroplast protein biogenesis network during heat acclimation. Plant Cell 34 (3): 1075–1099.

Trösch, R., Töpel, M., Flores-Pérez, Ú., and Jarvis, P. (2015b). Genetic and Physical Interaction Studies Reveal Functional Similarities between ALBINO3 and ALBINO4 in Arabidopsis. Plant physiology 169 (2): 1292–1306.

Trösch, R., and Willmund, F. (2019). The conserved theme of ribosome hibernation: from bacteria to chloroplasts of plants. Biological chemistry 400 (7): 879–893.

van Wijk, K.J. (2015). Protein maturation and proteolysis in plant plastids, mitochondria, and peroxisomes. Annu. Rev. Plant Biol. 66 (1): 75–111.

Voelker, R., and Barkan, A. (1995). Two nuclear mutations disrupt distinct pathways for targeting proteins to the chloroplast thylakoid. The EMBO journal 14 (16): 3905–3914.

Voelker, R., Mendel-Hartvig, J., and Barkan, A. (1997). Transposon-disruption of a maize nuclear gene, tha1, encoding a chloroplast SecA homologue: in vivo role of cp-SecA in thylakoid protein targeting. Genetics 145 (2): 467–478.

Walter, B., Hristou, A., Nowaczyk, M.M., and Schunemann, D. (2015). In vitro reconstitution of co-translational D1 insertion reveals a role of the cpSec-Alb3 translocase and Vipp1 in photosystem II biogenesis. The Biochemical journal 468 (2): 315–324.

Wang, P., and Dalbey, R.E. (2011). Inserting membrane proteins: the YidC/Oxa1/Alb3 machinery in bacteria, mitochondria, and chloroplasts. Biochimica et biophysica acta 1808 (3): 866–875.

Wang, P., and Grimm, B. (2015). Organization of chlorophyll biosynthesis and insertion of chlorophyll into the chlorophyll-binding proteins in chloroplasts. Photosynthesis research 126 (2-3): 189–202.

Wang, P., and Grimm, B. (2021). Connecting Chlorophyll Metabolism with Accumulation of the Photosynthetic Apparatus. Trends in plant science.

Westrich, L.D., Gotsmann, V.L., Herkt, C., Ries, F., Kazek, T., Trösch, R., Armbruster, L., Mühlenbeck, J.S., Ramundo, S., Nickelsen, J., Finkemeier, I., Wirtz, M., Storchová, Z., Räschle, M., and Willmund, F. (2021). The versatile interactome of chloroplast ribosomes revealed by affinity purification mass spectrometry. Nucleic acids research 49 (1): 400– 415.

Yukawa, M., Kuroda, H., and Sugiura, M. (2007). A new in vitro translation system for non-radioactive assay from tobacco chloroplasts: Effect of pre-mRNA processing on translation in vitro. The Plant journal for cell and molecular biology 49 (2): 367–376.

Zhang, L., Paakkarinen, V., van Wijk, K.J., and Aro, E.M. (1999). Co-translational assembly of the D1 protein into photosystem II. J. Biol. Chem. 274 (23): 16062–16067.

Zhao, Q., and Liu, C. (2017). Chloroplast Chaperonin: An Intricate Protein Folding Machine for Photosynthesis. Frontiers in molecular biosciences 4: 98.

Ziehe, D., Dünschede, B., and Schünemann, D. (2018). Molecular mechanism of SRP-dependent light-harvesting protein transport to the thylakoid membrane in plants. Photosynthesis research 138 (3): 303–313.

Zoschke, R., and Barkan, A. (2015). Genome-wide analysis of thylakoid-bound ribosomes in maize reveals principles of cotranslational targeting to the thylakoid membrane. Proc Natl Acad Sci U S A 112 (13): E1678–87.

Zoschke, R., and Bock, R. (2018). Chloroplast Translation: Structural and Functional Organization, Operational Control, and Regulation. Plant Cell 30 (4): 745–770.

