## Supplemental information for "Proteomic identification of the interactome of stalled ribosome nascent chain complexes translating the thylakoid membrane protein D1"

**Stolle et al.**

### Supplemental Figures

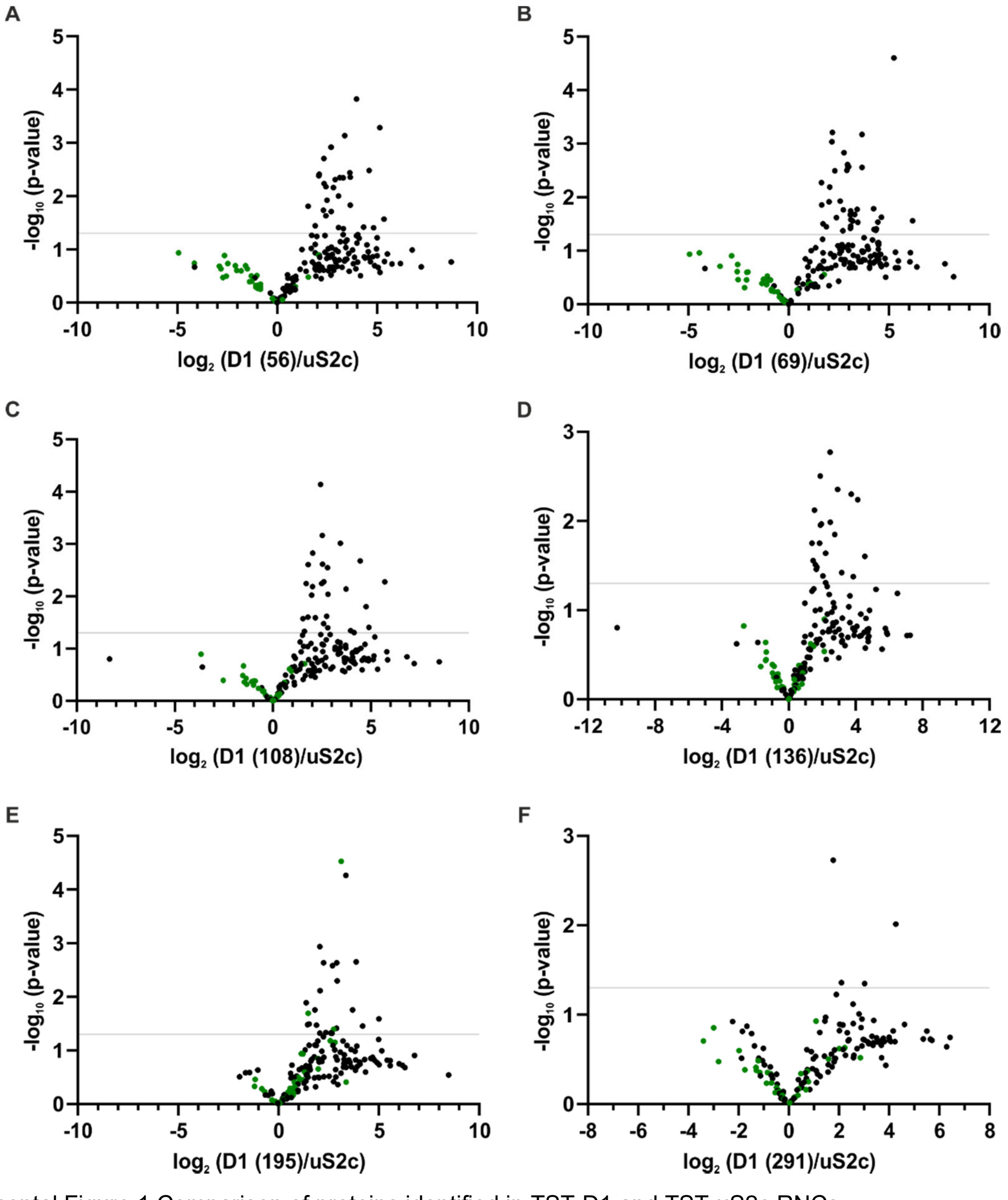


Supplemental Figure 1 Comparison of proteins identified in TST-D1 and TST-uS2c RNCs

Volcano plots representing proteins identified by tandem MS in RNCs with TST-uS2c (158) and TST-D1 peptides of different length (**A**-**F**). *P*-values (-$\log_{10}$) were plotted against the ratio of label-free quantification intensity means ($\log_2$). The *p*-values were determined by two-sided *t*-test ($\alpha = 0.05$). Ribosomal proteins are labeled in green.

**A**

Alb3 155-208

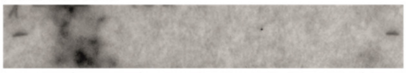

\- 1 2 3 4 5 6 7 8 9 10 11 -

| | | | |
|---|---|---|---|
| 1 | TYPLTKQQVESTLAM | 7 | IQQRYAGNQERIQLE |
| 2 | TKQQVESTLAMQNLQ | 8 | YAGNQERIQLETSRL |
| 3 | VESTLAMQNLQPKIK | 9 | QERIQLETSRLYKQA |
| 4 | LAMQNLQPKIKAIQQ | 10 | QLETSRLYKQAGVNP |
| 5 | NLQPKIKAIQQRYAG | 11 | TSRLYKQAGVNPLAG |
| 6 | KIKAIQQRYAGNQER | | |

Alb3 282-462

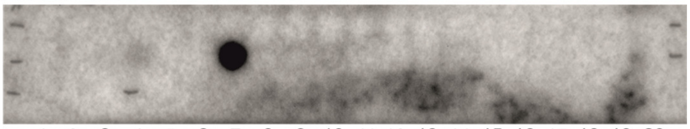

\- 1 2 3 4 5 6 7 8 9 10 11 12 13 14 15 16 17 18 19 20 -

\- 21 22 23 24 25 26 27 28 29 30 31 32 33 34 35 36 37 38 39 40 -

\- 41 42 43 -

| | | | | | |
|---|---|---|---|---|---|
| 1 | LVLPVLLIASQYVSM | 15 | WLTNNVLSTAQQVYL | 29 | EQEKRSKKNKAVAKD |
| 2 | VLLIASQYVSMEIMK | 16 | NVLSTAQQVYLRKLG | 30 | RSKKNKAVAKDTVEL |
| 3 | ASQYVSMEIMKPPQT | 17 | TAQQVYLRKLGGAKP | 31 | NKAVAKDTVELVEES |
| 4 | VSMEIMKPPQTDDPA | 18 | VYLRKLGGAKPNMDE | 32 | AKDTVELVEESQSES |
| 5 | IMKPPQTDDPAQKNT | 19 | KLGGAKPNMDENASK | 33 | VELVEESQSESEEGS |
| 6 | PQTDDPAQKNTLLVF | 20 | AKPNMDENASKIISA | 34 | EESQSESEEGSDDEE |
| 7 | DPAQKNTLLVFKFLP | 21 | MDENASKIISAGRAK | 35 | SESEEGSDDEEEEAR |
| 8 | KNTLLVFKFLPLMIG | 22 | ASKIISAGRAKRSIA | 36 | EGSDDEEEEAREGAL |
| 9 | LVFKFLPLMIGYFAL | 23 | ISAGRAKRSIAQPDD | 37 | DEEEEAREGALASST |
| 10 | FLPLMIGYFALSVPS | 24 | RAKRSIAQPDDAGER | 38 | EAREGALASSTTSKP |
| 11 | MIGYFALSVPSGLSI | 25 | SIAQPDDAGERFRQL | 39 | GALASSTTSKPLPEV |
| 12 | FALSVPSGLSIYWLT | 26 | PDDAGERFRQLKEQE | 40 | SSTTSKPLPEVGQRR |
| 13 | VPSGLSIYWLTNNVL | 27 | GERFRQLKEQEKRSK | 41 | SKPLPEVGQRRSKRS |
| 14 | LSIYWLTNNVLSTAQ | 28 | RQLKEQEKRSKKNKA | 42 | PEVGQRRSKRSKRKR |
| | | | | 43 | VGQRRSKRSKRKRTV |

**B**

Alb4 139-192

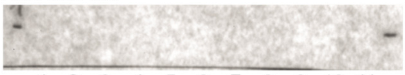

\- 1 2 3 4 5 6 7 8 9 10 11 -

| | | | |
|---|---|---|---|
| 1 | TFPLTKKQVESAMAM | 7 | IQERYAGDQEKIQLE |
| 2 | TKKQVESAMAMKSLT | 8 | YAGDQEKIQLETARL |
| 3 | VESAMAMKSLTPQIK | 9 | QEKIQLETARLYKLA |
| 4 | MAMKSLTPQIKAIQE | 10 | QLETARLYKLAGINP |
| 5 | SLTPQIKAIQERYAG | 11 | TARLYKLAGINPLAG |
| 6 | QIKAIQERYAGDQEK | | |

Alb4 266-499

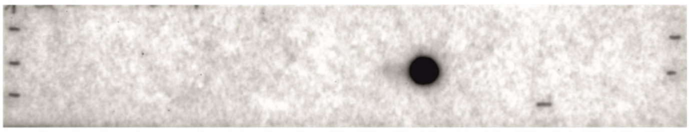

\- 1 2 3 4 5 6 7 8 9 10 11 12 13 14 15 16 17 18 19 20 -

\- 21 22 23 24 25 26 27 28 29 30 31 32 33 34 35 36 37 38 39 40 -

\- 41 42 43 44 45 46 47 48 49 50 51 52 53 54 55 56 -

| | | | | | |
|---|---|---|---|---|---|
| 1 | LVLPLLLVFSQYLSI | 20 | AKNPVEKFTNLVTKE | 39 | QKAEAALSNQNTDKA |
| 2 | LLLVFSQYLSIQIMQ | 21 | VEKFTNLVTKEDKTQ | 40 | AALSNQNTDKAHEQD |
| 3 | FSQYLSIQIMQSSQS | 22 | TNLVTKEDKTQQIEK | 41 | NQNTDKAHEQDEKSD |
| 4 | LSIQIMQSSQSNDPA | 23 | TKEDKTQQIEKSFSE | 42 | DKAHEQDEKSDTAIV |
| 5 | IMQSSQSNDPAMKSS | 24 | KTQQIEKSFSEPLVQ | 43 | EQDEKSDTAIVAEDD |
| 6 | SQSNDPAMKSSQAVT | 25 | IEKSFSEPLVQKSVS | 44 | KSDTAIVAEDDKKTE |
| 7 | DPAMKSSQAVTKLLP | 26 | FSEPLVQKSVSELKI | 45 | AIVAEDDKKTELSAV |
| 8 | KSSQAVTKLLPLMIG | 27 | LVQKSVSELKIPREK | 46 | EDDKKTELSAVDETS |
| 9 | AVTKLLPLMIGYFAL | 28 | SVSELKIPREKGGEK | 47 | KTELSAVDETSDGTV |
| 10 | LLPLMIGYFALSVPS | 29 | LKIPREKGGEKVTPE | 48 | SAVDETSDGTVAVNG |
| 11 | MIGYFALSVPSGLSL | 30 | REKGGEKVTPESPKP | 49 | ETSDGTVAVNGKPSI |
| 12 | FALSVPSGLSLYWLT | 31 | GEKVTPESPKPGERF | 50 | GTVAVNGKPSIQKDE |
| 13 | VPSGLSLYWLTNNIL | 32 | TPESPKPGERFRLLK | 51 | VNGKPSIQKDETTNG |
| 14 | LSLYWLTNNILSTAQ | 33 | PKPGERFRLLKEQEA | 52 | PSIQKDETTNGTFGI |
| 15 | WLTNNILSTAQQVWL | 34 | ERFRLLKEQEAKRRR | 53 | KDETTNGTFGIGHDT |
| 16 | NILSTAQQVWLQKYG | 35 | LLKEQEAKRRREKEE | 54 | TNGTFGIGHDTEQQH |
| 17 | TAQQVWLQKYGGAKN | 36 | QEAKRRREKEERQKA | 55 | FGIGHDTEQQHSHET |
| 18 | VWLQKYGGAKNPVEK | 37 | RRREKEERQKAEAAL | 56 | GHDTEQQHSHETEKR |
| 19 | KYGGAKNPVEKFTNL | 38 | KEERQKAEAALSNQN | | |

Supplemental Figure 2 Membrane bound peptide arrays for pepspot interaction analyses

Protein-protein interaction analyses were performed with membrane bound 15mer peptides and recombinant, His-tagged proteins. (**A**) Peptides covering Alb3 residues 155-208 and 282-462, shown under membranes, were incubated with His-mSTIC2 (aa 49-182) at a final concentration of 5 µg/ml. (**B**) Peptides covering Alb4 residues 139-192 and 266-499 were analyzed as in **A**. An Antibody-HRP conjugate against the His-tag was used to detect His-mSTIC2 protein on pepspot membranes.

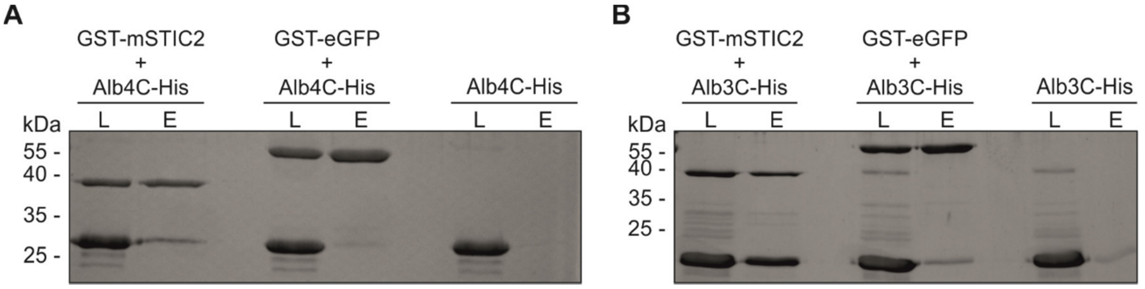


Supplemental Figure 3 STIC2 *in vitro* pulldown interaction analyses

*In vitro* pulldown experiments were performed with equal amounts of GST- and His-tagged proteins using glutathione resin. Loading (L) and eluate (E) samples were analyzed by SDS-PAGE and Coomassie blue staining. (**A**) Recombinant Alb4C-His (aa 334-499) was incubated with GST-mSTIC2 (aa 49-182). The Alb4C protein alone or incubated with GST-eGFP served as control samples. (**B**) Pulldown analyses with Alb3C-His (aa 350-462) were performed as described in **A**.

### Supplemental Tables

#### Supplemental Table 1 Proteins w/o significant quantitative differences in TST-D1 and TST-uS2c (158) RNCs;

Proteins indicated with asterisks were additionally identified in TST-D1 RNCs with another Razor accession (Razor Psat#) but not found in TST-uS2c (158) RNCs.

| Uniprot ID | Uniprot Name | Category | Function | Razor Psat# | Potential AT# | found in D1 RNC |
|---|---|---|---|---|---|---|
| O81004 | Y2287 | | | Psat0s2458g0040.1 | AT2G22870.1 | s, m, l |
| | | | | Psat7g113880.1 | | s, m, l |
| Q84WN0 | Y4920 | | | Psat1g169560.1 | AT4G37920.1 | s, m, l |
| O80934 | Y2766 | | | Psat2g068960.2 | AT2G37660.1 | s, m, l |
| A0A3Q0ETQ4 | A0A3Q0ETQ4 | | | Psat4g161440.1 | | s, m, l |
| A0A392R6B6 | A0A392R6B6 | | | Psat5g189080.1 | | s, m, l |
| Q8RWG3 | RER6 | | | Psat5g246640.1 | AT3G56140.1 | s, l |
| Q9C9F5 | C3H15 | | | Psat6g223920.1 | AT1G68200.1 | s, m, l |
| Q9LXC9 | IPYR6 | | Hydrolase | Psat1g212360.1 | AT5G09650.1 | s |
| P54150 | MSRA4 | | Peptide methionine sulfoxide reductase A4 | Psat2g010960.3 | AT4G25130.1 | m |
| Q8H0V1 | CK5P1 | | Potential kinase regulator, [4Fe-4S] binding | Psat0s3355g0160.1 | AT4G36390.1 | s, m, l |
| Q8S9M1 | PAP13 | | Probable plastid lipid associated protein | Psat0s4417g0040.1 | AT2G42130.1 | s, m, l |
| G7JYB8 | G7JYB8 | | Putative metallo-beta-lactamase | Psat2g091800.1 | | s, m, l |
| Q9M0Y6 | DPNPM | | Putative PAP-specific phosphatase, mitochondrial | Psat1g209080.1 | AT4G05090.1 | s, m, l |
| G7JR97 | G7JR97 | | Putative ribosomal protein L2 domain 2 | Psat7g241800.1 | | s, m, l |
| Q9M098 | EBFC1 | | STIC2-like protein | Psat4g009640.2 | AT4G30620.1 | s, m, l |
| Q03943 | IM30 | Biogenesis | Thylakoid biogenesis | Psat7g208480.2 | AT1G65260.1 | s, m, l |
| Q9LUJ3 | RDM1 | DNA processing | | Psat2g190480.2 | AT3G22680.1 | s |
| O78328 | DXS | Metabolism | | Psat4g029960.2 | AT4G15560.1 | s, m, l |
| O81360 | ABA2 | Metabolism | Abscisate biosynthesis | Psat0s1281g0080.1 | AT5G67030.1 | s, m, l |
| B9SL58 | PURA2 | Metabolism | Adenylosuccinate synthetase 2 | Psat3g071360.1 | AT3G57610.1 | m |
| P04717 | RBL | Metabolism | Calvin Cycle | Psat0s5127g0120.1 | AtCg00490 | s, m, l |
| | | | | Psat0s1330g0280.1 | | s, m, l |
| O65194 | RBS | Metabolism | Calvin Cycle | Psat1g080440.1 | AT1G67090 | s, m, l |
| P12858 | G3PA | Metabolism | Calvin Cycle | Psat3g091720.1 | AT1G12900 | s, m, l |
| P00869 | RBS2 | Metabolism | Calvin Cycle | Psat3g205240.1 | AT1G67090 | s, m, l |
| P12859 | G3PB | Metabolism | Calvin Cycle | Psat5g010520.1 | AT1G42970.1 | s, m, l |
| O20252 | S17P | Metabolism | Calvin Cycle | Psat5g127280.2 | AT3G55800 | s, m, l |
| F4IW47 | TKTC2 | Metabolism | Calvin Cycle | Psat6g238800.1 | AT2G45290.2 | s, m, l |
| P28552 | ATPG | Metabolism | CP ATP synthase gamma chain | Psat3g207840.1 | AT4G04640 | s, m, l |
| P08215 * | ATPA | Metabolism | CP ATP synthase subunit alpha | Psat3g186480.1 | AtCg00120 | s, m, l |
| P05037 | ATPB | Metabolism | CP ATP synthase subunit beta | Psat2g136800.1 | ATCG00480.1 | s, m, l |
| | | | | Psat4g098720.1 | | s, m, l |
| O24457 | ODPA3 | Metabolism | Carbon metabolism | Psat0s3525g0080.1 | AT1G01090.1 | s |

| | | | | | | |
|---|---|---|---|---|---|---|
| Q9SQI8 | ODP24 | Metabolism | Carbon metabolism | Psat6g228960.1 | AT3G25860.1 | s, m, l |
| Q01516 | ALFC1 | Metabolism | Carbon metabolism | Psat7g139080.1<br>Psat7g139120.1 | AT2G21330 | s, m, l<br>s, m, l |
| P52417 | GLGS2 | Metabolism | Starch metabolism | Psat5g110720.1 | AT5G48300.1 | s, m, l |
| Q8LB01 | DAPB2 | Metabolism - amino acids | 4-hydroxy-tetrahydrodipicolinate reductase 2 | Psat4g089800.1 | AT3G59890.1 | m |
| Q55512 | DHAS | Metabolism - amino acids | Aspartate-semialdehyde dehydrogenase | Psat4g004640.1 | AT1G14810.1 | s, m |
| O49485 | SERA1 | Metabolism - amino acids | D-3-phosphoglycerate dehydrogenase 1 | Psat6g219520.1 | AT4G34200.1 | s, m, |
| P08281 | GLNA2 | Metabolism - amino acids | Glutamine synthetase | Psat1g185960.3 | AT5G35630 | m, l |
| Q9C550 | LEU12 | Metabolism - amino acids | Isopropylmalate synthase 2 | Psat6g241040.1 | AT1G74040.1 | s, m, l |
| Q93ZN9 | DAPAT | Metabolism - amino acids | LL-diaminopimelate aminotransferase | Psat7g104520.1 | AT4G33680.1 | s, m, l |
| Q42777 | MCCA | Metabolism - amino acids | MC Methylcrotonoyl-CoA carboxylase subunit alpha | Psat6g059920.1 | AT1G03090.1 | s, m, l |
| Q94JQ3 | GLYP3 | Metabolism - amino acids | Serine hydroxymethyltransferase 3 | Psat1g192400.2 | AT4G32520.1 | s, m, l |
| Q1H537 | DCVR | Metabolism - chlorophyll | Divinyl chlorophyllide a 8-vinyl-reductase | Psat1g187280.1 | AT5G18660.1 | s, m, l |
| P45621 | GSA | Metabolism - chlorophyll | Glutamate-1-semialdehyde 2,1-aminomutase | Psat5g001920.1 | AT3G48730.1 | s, m, l |
| O22437 | CHLD | Metabolism - chlorophyll | Magnesium-chelatase subunit D | Psat0s3604g0040.1 | AT1G08520.1 | s, m, l |
| Q9FNB0 | CHLH | Metabolism - chlorophyll | Magnesium-chelatase subunit H | Psat6g120040.1 | AT5G13630.1 | s, m, l |
| Q06881 | BCCP | Metabolism - fatty acid | Biotin carboxyl carrier protein of acetyl-CoA carboxylase | Psat2g029880.1<br>Psat5g126160.1 | AT5G15530 | s, m, l<br>s, m, l |
| Q42783 | BCCP | Metabolism - fatty acid | Biotin carboxyl carrier protein of acetyl-CoA carboxylase | Psat3g102480.1 | AT5G16390 | s, m, l |
| Q9LLC1 | BCCP2 | Metabolism - fatty acid | Biotin carboxyl carrier protein of acetyl-CoA carboxylase 2 | Psat0s846g0040.1 | AT5G15530.1 | s, m, l |
| B9HBA8 | ACCC1 | Metabolism - fatty acid | Biotin carboxylase 1 | Psat0s916g0080.1 | AT5G35360 | s, m, l |
| Q42807 | STAD | Metabolism - fatty acid | Stearoyl-9-desaturase | Psat7g119680.1 | AT2G43710.1 | s, m |
| Q9LS25 | PP420 | Pentatricopeptide repeat | | Psat4g078640.1 | AT5G46580.1 | s |
| Q9M3A8 | PP273 | Pentatricopeptide repeat | | Psat5g077680.1 | AT3G49240.1 | s, m, l |
| Q94AQ8 | PNSB2 | Photosynthesis | Photosynthetic NDH subunit of subcomplex B 2 | Psat3g089080.1 | AT1G64770.1 | s, l |
| Q8SKU2 | TIC62 | Protein import | Redox-dependent import regulation | Psat7g185400.1 | AT3G18890.1 | s, m, l |
| Q8VZ74 | ERA | Protein processing | Ribosome biogenesis | Psat4g094880.1 | AT5G66470.1 | s, m, l |
| O98997 | RCA | Protein processing | Rubisco Activase | Psat5g123320.3 | AT2G39730 | s, m, l |
| Q7X9A0 | RCA1 | Protein processing | Rubisco Activase 1 | Psat2g039960.1 | AT2G39730.1 | s, m, l |
| Q02028 | HSP7S | Protein processing - chaperone | Chaperone | Psat1g222720.1<br>Psat1g222760.1 | AT5G49910.1 | s, m, l<br>s, m, l |
| Q9C667 | CPNB4 | Protein processing - chaperone | Chaperone | Psat2g080360.1 | AT1G26230.1 | m |
| O65282 * | CH20 | Protein processing - chaperone | Chaperone | Psat4g019040.1 | AT5G20720.1 | s, m, l |
| P35100 | CLPC | Protein processing - chaperone | Chaperone | Psat5g054560.2<br>Psat7g039080.1 | AT5G50920.1 | s, m, l<br>s, m, l |
| Q9SJZ7 | DNJA6 | Protein processing - chaperone | Chaperone | Psat5g090520.1 | AT2G22360.1 | s, m, l |
| Q05046 | CH62 | Protein processing - chaperone | Chaperone mitochondrial | Psat6g170600.1 | AT3G23990 | s, m, l |
| P36774 | LON2 | Protein processing - protease | Protease | Psat0s471g0440.1 | AT1G75460.1 | s, m, l |

| | | | | | | |
|---|---|---|---|---|---|---|
| Q94B60 | CLPP4 | Protein processing - protease | Protease | Psat2g142240.1 | AT5G45390.1 | s, m, l |
| Q8LB10 | CLPR4 | Protein processing - protease | Protease | Psat4g075600.1 | AT4G17040.1 | s, m, l |
| Q9XJ35 | CLPR1 | Protein processing - protease | Protease | Psat6g215320.4 | AT1G49970.1 | s, m, l |
| Q40983 | SPP | Protein processing - protease | Stromal processing peptidase | Psat2g189160.1 | AT5G42390 | s, m, l |
| Q9SR19 | RAF2 | Protein processing - PTM | Rubisco marturation | Psat6g143080.1 | AT3G04550.1 | s, m, l |
| Q43088 | RBCMT | Protein processing - PTM | Rubisco methyltransferase | Psat7g156240.1 | AT1G14030.1 | s, m, l |
| P48384 | TRXM | Redox | | Psat5g081600.1 | AT4G03520 | s, m, l |
| Q9MB35 | PERQ | Redox | Peroxidase activity | Psat4g018640.1 | AT3G26060.1 | s, m, l |
| B5LMR0 | RK16 | Ribosome - chloroplast | CP Ribosome | Psat0s1446g0200.1 | AtCg00790.1 | m, l |
| Q8VYM4 * | PSRP2 | Ribosome - chloroplast | CP Ribosome | Psat0s2674g0040.2 | AT3G52150.1 | s, m, l |
| P82195 | RK18 | Ribosome - chloroplast | CP Ribosome | Psat1g016560.1 | AT1G48350.1 | m |
| P23408 | RK22 | Ribosome - chloroplast | CP Ribosome | Psat1g052200.1 | ATCG00810.1 | s, m, l |
| P36210 | RK121 | Ribosome - chloroplast | CP Ribosome | Psat1g056240.1 | AT3G27830.1 | s, m, l |
| O23049 | RK6 | Ribosome - chloroplast | CP Ribosome | Psat1g066560.1 | AT1G05190.1 | s, m, l |
| B5LMQ9 | RK14 | Ribosome - chloroplast | CP Ribosome | Psat1g099160.1<br>Psat5g284200.1 | AtCg00780.1 | s, m, l<br>m |
| P29344 | RR1 | Ribosome - chloroplast | CP Ribosome | Psat1g162760.1 | At5g30510.1 | s, m, l |
| P82413 | RK19 | Ribosome - chloroplast | CP Ribosome | Psat2g176160.2 | AT4G17560.1 | s, m, l |
| O04603 | RK5 | Ribosome - chloroplast | CP Ribosome | Psat3g107160.1 | AT4G01310.1 | s, m, l |
| P82024 | RR21 | Ribosome - chloroplast | CP Ribosome | Psat3g169680.2 | AT3G27160.1 | s, m, l |
| O80362 | RK10 | Ribosome - chloroplast | CP Ribosome | Psat4g007600.1 | AT5G13510.1 | s, m, l |
| P82130 | RR20 | Ribosome - chloroplast | CP Ribosome | Psat4g136440.1 | At3g15190.1 | s, m, l |
| Q9SYL9 | RK13 | Ribosome - chloroplast | CP Ribosome | Psat4g140640.1 | AT1G78630.1 | s, m, l |
| O80361 | RK4 | Ribosome - chloroplast | CP Ribosome | Psat4g224360.1 | AT1G07320.1 | s, m, l |
| Q9MAP3 | RK11 | Ribosome - chloroplast | CP Ribosome | Psat5g110320.1 | AT1G32990.1 | s, m, l |
| Q6KGX3 | RR7 | Ribosome - chloroplast | CP Ribosome | Psat5g119080.1 | AtCg00900.1<br>AtCg01240.1 | s, m, l |
| P42732 | RR13 | Ribosome - chloroplast | CP Ribosome | Psat5g134240.2 | AT5G14320.1 | s, m, l |
| P15820 | RK32 | Ribosome - chloroplast | CP Ribosome | Psat5g142800.1 | AtCg01020.1 | l |
| P31163 | RK2 | Ribosome - chloroplast | CP Ribosome | Psat5g168160.1 | AtCg00830.1<br>AtCg01310.1 | s, m, l |
| Q9M4Y3 | RR10 | Ribosome - chloroplast | CP Ribosome | Psat5g170120.1 | AT3G13120.1 | s, m, l |
| P82190 | RK27 | Ribosome - chloroplast | CP Ribosome | Psat5g215240.1 | AT5G40950.1 | s, m, l |
| P11893 | RK24 | Ribosome - chloroplast | CP Ribosome | Psat5g249200.1 | At5g54600.1 | s, m, l |
| Q9FWS4 | RK31 | Ribosome - chloroplast | CP Ribosome | Psat6g009240.1 | AT1G75350.1 | s, m, l |
| P82278 | RR9 | Ribosome - chloroplast | CP Ribosome | Psat6g010760.1 | At1g74970.1 | s, m, l |
| P51412 | RK21 | Ribosome - chloroplast | CP Ribosome | Psat6g187560.1 | AT1G35680.1 | s, m, l |
| P08241 | RR2 | Ribosome - chloroplast | CP Ribosome | Psat6g219440.1 | ATCG00160 | s |

| | | | | | | |
|---|---|---|---|---|---|---|
| P31165 | RK15 | Ribosome - chloroplast | CP Ribosome | Psat6g225000.1 | At3g25920.1 | s, m, l |
| Q8VZ55 | RK35 | Ribosome - chloroplast | CP Ribosome | Psat7g006560.1 | AT2G24090.1 | s, m, l |
| Q8VY91 | RR6 | Ribosome - chloroplast | CP Ribosome | Psat7g023960.1 | AT1G64510.1 | s, m, l |
| P82248 | RK29 | Ribosome - chloroplast | CP Ribosome | Psat7g055240.1 | At5g65220.1 | s, m, l |
| Q9ST69 | RR5 | Ribosome - chloroplast | CP Ribosome | Psat7g115360.1 | At2g33800.1 | s, m, l |
| Q9LY66 | RK1 | Ribosome - chloroplast | CP Ribosome | Psat7g200640.1 | AT3G63490.1 | s, m, l |
| P11894 | RK9 | Ribosome - chloroplast | CP Ribosome | Psat7g223760.1 | At3g44890.1 | s, m, l |
| Q8L7S8 | RH3 | Ribosome biogenesis | rRNA processing | Psat0s2843g0040.1<br>Psat3g015480.1 | AT5G26742.1 | s, m, l<br>s, m, l |
| Q9SA52 | CP41B | Ribosome biogenesis | rRNA processing | Psat3g014320.1 | AT1G09340.1 | l |
| Q9FFQ1 | RH31 | RNA - helicase | DEAD-box RNA helicase | Psat4g214280.1 | AT5G63630.1 | s, m, l |
| Q9FGS0 | CP31B | RNA processing | mRNA processing | Psat0s1670g0040.1 | AT5G50250.1 | s, m, l |
| Q04836 | CP31A | RNA processing | mRNA processing | Psat5g118040.2 | AT4G24770.1 | s, m, l |
| Q9ZUU4 | CP29B | RNA processing | mRNA processing | Psat6g071720.1 | AT2G37220.1 | s, m, l |
| O34331 | YLBH | RNA processing | Putative rRNA methyltransferase YlbH | Psat1g076480.1 | | s, m, l |
| Q42586 | UMPS | RNA processing - synthase | Uridine-monophosphate synthase | Psat0s1718g0040.2 | AT3G54470.1 | m, l |
| A3PCG2 | EFTS | Translation regulation | Translation | Psat1g044600.1 | AT4G29060 | s, m, l |
| I1K0K6 | EFGC2 | Translation regulation | Translation | Psat4g069360.2 | AT1G62750.1 | s, m, l |
| F4K410 | SVR3 | Translation regulation | Translation | Psat6g127960.2 | AT5G13650.1 | s, m, l |
| O81983 | SYS | tRNA processing | tRNA ligase | Psat6g149440.1 | At1g11870.1 | s, m, l |

**Supplemental Table 2 Proteins enriched or specifically found in TST-D1 RNCs**

Proteins indicated with asterisks were identified exclusively in TST-D1 RNCs shown.

| Uniprot ID | Uniprot Name | Category | Function | Razor Psat# | Potential AT# | enriched in D1 RNC |
|---|---|---|---|---|---|---|
| Q6DYE4 * | Y1609 | | | Psat0s629g0200.5 | AT1G26090.1 | l |
| A0A1S2XK40 * | A0A1S2XK40 | | | Psat1g069960.2 | | s |
| Q9SKN5 * | ARFJ | | | Psat2g033160.1 | AT2G28350.1 | m |
| Q9ASX5 * | Y5520 | | | Psat2g061840.1 | AT5G05200.1 | s |
| O80934 | Y2766 | | | Psat3g138600.1 | AT2G37660.1 | s, m |
| A0A1S2Y854 * | A0A1S2Y854 | | | Psat5g011440.1 | At1G63610 | s, m, l |
| G7I7C8 * | G7I7C8 | | | Psat6g095240.1 | AT5G04440 | s |
| Q9LVP0 * | Y5639 | | | Psat6g141840.1 | AT5G63930.1 | l |
| A0A2K3P005 * | COBWD | | | Psat6g237480.1 | AT1G80480.1 | s, l |
| A0A2K3NKT1 | A0A2K3NKT1 | | | Psat7g260440.1 | | m |
| Q9ZP40 | PG1 | | Plastoglobulin-1 | Psat2g138880.1 | AT2G35490 | s, l |
| A0A1S3E5A8 | A0A1S3E5A8 | | Putative armadillo repeat superfamily protein | Psat6g044440.4 | AT1G23180 | s |
| P43394 | K502 | | Putative FAD-NADP Oxidoreductase-like | Psat6g222520.1 | AT1G15140.1 | m, l |
| A0A2Z6NLS5 | A0A2Z6NLS5 | | Putative heavy metal transport/detoxification superfamily protein | Psat1g080040.1 | | l |
| Q9LQK7 * | AB7I | | SufD homolog, potential [4Fe-4S] repair | Psat2g173560.2 | AT1G32500.1 | s, m, l |
| Q9M394 * | SCKL1/FLN1 | Biogenesis | Chloroplast biogenesis | Psat7g243520.1 | AT3G54090.1 | s, m, l |
| P37107 | cpSRP54 | Biogenesis | Protein targeting | Psat3g032320.1 | AT5G03940.1 | s, m, l |
| O82230 * | STIC2 | Biogenesis | Thylakoid biogenesis | Psat1g022480.1 | AT2G24020.1 | s, m, l |
| Q1KPV0 * | FZL | Biogenesis | Thylakoid biogenesis | Psat6g060000.3 | AT1G03160.1 | s, m |
| Q42545 * | FTSZ1 | Cell cycle | Cell division protein FtsZ homolog 1 | Psat2g010320.1 | AT5G55280.1 | s, l |
| O82533 | FTZ21 | Cell cycle | Cell division protein FtsZ homolog 2-1 | Psat6g081080.2 | AT2G36250.1 | s |
| Q6STH5 | HF101 | Fe-S assembly | Assembly of Fe-S clusters on selected proteins,<br>ATP dependent /4Fe-4S insertion | Psat1g039560.1 | AT3G24430.1 | m |
| Q9LIB2 * | PHS1 | Metabolism | | Psat1g066720.1 | AT3G29320.1 | m |
| Q9SH69 * | 6PGD1 | Metabolism | | Psat7g028480.1 | AT1G64190.1 | m |
| Q42961 | PGKH | Metabolism | Calvin Cycle | Psat1g014040.1 | AT1G56190 | s, m |
| P26302 * | KPPR | Metabolism | Calvin Cycle | Psat2g140680.1 | AT1G32060.1 | s |
| P46275 * | F16P1 | Metabolism | Calvin Cycle | Psat3g134680.1 | AT3G54050.1 | s |
| Q43157 | RPE | Metabolism | Calvin Cycle | Psat4g049960.2 | AT5G61410.1 | m, l |
| Q9C8P0 | ODP25 | Metabolism | Carbon metabolism | Psat0s2229g0080.1 | AT1G34430.1 | s, l |
| Q9SAJ6 * | G3PP1 | Metabolism | Carbon metabolism | Psat1g005000.1 | AT1G79530.1 | s |
| Q9FLW9 * | PKP2 | Metabolism | Carbon metabolism<br>Carbon metabolism | Psat1g096960.1<br>Psat6g194760.1 * | AT5G52920.1 | s, m, l<br>s, m, l |
| A8MS68 | PLPD1 | Metabolism | Carbon metabolism | Psat4g023520.1 | AT3G16950.1 | s, m |
| Q43117 | KPYA | Metabolism | Carbon metabolism | Psat6g113960.1 | AT3G22960.1 | s, m, l |
| Q9ZU52 * | ALFP3 | Metabolism | Carbon metabolism | Psat6g223200.1 | AT2G01140.1 | s, m, l |
| P17067 | CAHC | Metabolism | Carbonic anhydrase | Psat1g058960.5 | AT3G01500 | s |
| P08215 | ATPA | Metabolism | CP ATP synthase subunit alpha | Psat2g111080.1<br>Psat6g219360.1 | AtCg00120 | m<br>m |
| Q94ID7 * | GGPPS | Metabolism | Geranylgeranyl pyrophosphate synthase | Psat4g091240.1 | AT4G36810.1 | s |

| | | | | | | |
|---|---|---|---|---|---|---|
| F4K0E8 * | ISPG | Metabolism | Isoprene biosynthesis | Psat5g278240.1 | AT5G60600.1 | s, m |
| Q9SN86 * | MDHP | Metabolism | Malate dehydrogenase | Psat4g051000.1 | AT3G47520.1 | s, m, l |
| Q00218 | AROG | Metabolism | Phospho-2-dehydro-3-deoxyheptonate aldolase 2, chorismate biosynthesis | Psat1g216320.1 | AT4G33510.1 | s, m |
| P52418 * | PUR1 | Metabolism | Purine metabolism | Psat7g181880.1 | AT2G16570.1 | s, m, l |
| Q94A41 | AMY3 | Metabolism | Starch metabolism | Psat4g132240.1 | AT1G69830.1 | s, m |
| Q6ZY51 * | PWD | Metabolism | Starch metabolism | Psat6g244000.1 | AT5G26570.1 | s, l |
| Q84KI6 * | SQD1 | Metabolism | Sulfolipids biosynthesis | Psat3g076360.2 | AT4G33030.1 | s |
| Q8W250 * | DXR | Metabolism | Terpenoid metabolism | Psat4g058000.1 | AT5G62790.1 | s, m, l |
| F6H7K5 | THI42 | Metabolism | Thiamine thiazole synthase 2 | Psat6g209120.1 | AT5G54770.1 | s, m |
| A0A072UP88 * | TAL | Metabolism | Transaldolase | Psat4g070360.2 | AT5G13420.1 | s |
| Q94B35 | ISPH | Metabolism | | Psat7g185320.1 | AT4G34350.1 | s, m |
| Q94AR8 | LEUC | Metabolism - amino acids | 3-isopropylmalate dehydratase large subunit | Psat6g107560.1 | AT4G13430.1 | s |
| Q93YZ7 | ILVH2 | Metabolism - amino acids | Acetolactate synthase small subunit 2 | Psat7g063320.1 | AT2G31810.1 | s, m |
| Q9SCL7 * | NAGK | Metabolism - amino acids | Acetylglutamate kinase | Psat5g026120.1 | AT3G57560.1 | s |
| Q9LEU8 * | ARLY | Metabolism - amino acids | Argininosuccinate lyase | Psat5g048920.1 | AT5G10920.1 | s, m |
| Q9SZX3 * | ASSY | Metabolism - amino acids | Argininosuccinate synthase | Psat0s3176g0120.1 *<br>Psat4g065240.1 | AT4G24830.1 | s, m, l<br>m |
| Q9SGD6 * | AROD6 | Metabolism - amino acids | Arogenate dehydratase/prephenate dehydratase 6 | Psat0s1945g0080.1 | AT1G08250.1 | m |
| Q8GSJ1 * | HIS1B | Metabolism - amino acids | ATP phosphoribosyltransferase 2 | Psat1g223400.1 | AT1G09795.1 | s |
| P37142 | AKH | Metabolism - amino acids | Bifunctional aspartokinase/homoserine dehydrogenase | Psat7g075960.1 | AT4G19710.1 | s, m, l |
| O81852 | AKH2 | Metabolism - amino acids | Bifunctional aspartokinase/homoserine dehydrogenase 2 | Psat5g099120.1 | AT4G19710.1 | s |
| Q42601 | CARB | Metabolism - amino acids | Carbamoyl-phosphate synthase large chain | Psat5g277200.2 | AT1G29900.1 | s |
| P55217 * | CGS1 | Metabolism - amino acids | Cystathionine gamma-synthase 1 | Psat3g199280.1 | AT3G01120.1 | s |
| Q8L7R2 | KHSE | Metabolism - amino acids | Homoserine kinase | Psat2g000800.1 | AT2G17265.1 | s, m |
| O82043 | ILV5 | Metabolism - amino acids | Ketol-acid reductoisomerase | Psat7g165680.3 | AT3G58610.1 | m |
| I3SWS0 * | I3SWS0 | Metabolism - amino acids | Probable bifunctional methylthioribulose-1-phosphate dehydratase/enolase-phosphatase E1 | Psat6g155960.2 | At5g53850 | l |
| Q41651 * | CYPB | Metabolism - amino acids | Peptidyl-prolyl cis-trans isomerase | Psat6g157800.1 | AT3G62030.1 | m, l |
| P25269 * | TRBP2 | Metabolism - amino acids | Tryptophan synthase beta chain 2 | Psat5g194440.1 | AT4G27070.1 | s |
| Q43155 * | GLTB | Metabolism - amino acids, nitrogen | Ferredoxin-dependent glutamate synthase | Psat3g078160.3 | AT5G04140.1 | s, l |
| P30124 * | HEM2 | Metabolism - chlorophyll | Delta-aminolevulinic acid dehydratase | Psat5g087720.1 | AT1G69740.1 | s, l |
| Q9ZS34 | CHLP | Metabolism - chlorophyll | Geranylgeranyl diphosphate reductase | Psat4g050200.1 | AT1G74470.1 | l |
| Q9SW18 | CHLM | Metabolism - chlorophyll | Magnesium protoporphyrin IX methyltransferase | Psat7g039320.1 | AT4G25080.1 | s, m, l |
| P93162 | CHLI | Metabolism - chlorophyll | Magnesium-chelatase subunit I | Psat1g200160.2 | At4g18480.1 | m |
| P35055 * | HEM6 | Metabolism - chlorophyll | Oxygen-dependent coproporphyrinogen-III oxidase | Psat2g001760.1 | AT1G03475.1 | s, m |
| Q43082 | HEM3 | Metabolism - chlorophyll | Porphobilinogen deaminase | Psat7g172760.1 | AT5G08280.1 | s |
| Q01289 | POR | Metabolism - chlorophyll | Protochlorophyllide reductase | Psat5g245240.1 | AT5G54190 | s, m, l |
| Q93X62 | FABG1 | Metabolism - fatty acid | 3-oxoacyl-reductase 1 | Psat5g094480.1 | AT1G24360.1 | s, m |
| Q41008 | ACCA | Metabolism - fatty acid | Acetyl-CoA carboxylase carboxyl transferase subunit alpha | Psat0s1978g0280.1 | AT2G38040.1 | m |
| P18823 | ACCD | Metabolism - fatty acid | Acetyl-CoA carboxylase carboxyl transferase subunit beta | Psat0s6989g0080.1 | ATCG00500.1 | s |
| Q42533 | BCCP1 | Metabolism - fatty acid | Biotin carboxyl carrier protein of acetyl-CoA carboxylase 1 | Psat1g067360.1 | AT5G16390.1 | s |
| Q9SLA8 * | FABI | Metabolism - fatty acid | NAD-dependent enoyl-reductase | Psat3g150560.1 | AT2G05990.1 | m |

| | | | | | | |
|---|---|---|---|---|---|---|
| Q42561 * | FATA1 | Metabolism - fatty acid | Oleoyl-acyl carrier protein thioesterase 1 | Psat3g149200.1 | AT3G25110.1 | m |
| B9FSC8 * | OPR11 | Metabolism - fatty acid | Oxylipin biosynthesis | Psat2g190280.2 | AT1G76690.1 | s, l |
| P06585 * | PSBA | Photosynthesis | Photosynthesis | Psat0ss8367g0080.1 | ATCG00020.1 | s, m, l |
| P10933 | FENR1 | Photosynthesis | Photosynthesis | Psat2g154080.2 | AT5G66190.1 | s, m |
| Q8LDL0 | HHL1 | Photosynthesis - repair | PSII repair cycle | Psat5g093920.2 | AT1G67700.1 | s, m |
| A0A072V684 * | A0A072V684 | Protein processing | Acyl-CoA N-acyltransferase (NAT) superfamily protein | Psat1g134240.1 | | s |
| P08927 | RUBB | Protein processing - chaperone | Chaperone | Psat1g001680.1<br>Psat6g016920.1 | AT1G55490 | s, m, l<br>s, m, l |
| P08926 | RUBA | Protein processing - chaperone | Chaperone | Psat7g144320.1 | AT2G28000.1 | s, m, l |
| Q9LF37 * | CLPB3 | Protein processing - chaperone | Chaperone | Psat1g069480.1 | AT5G15450.1 | s, m |
| O65282 | CH20 | Protein processing - chaperone | Chaperone | Psat1g135800.1 | AT5G20720.1 | m |
| Q8S9L5 | TIG | Protein processing - chaperone | Chaperone | Psat2g009800.1 | AT5G55220.1 | s |
| Q9SIF2 | HS905 | Protein processing - chaperone | Chaperone | Psat2g102720.1 | AT2G04030.1 | s, l |
| G7L2M8 | GRPE | Protein processing - chaperone | Chaperone | Psat3g085320.3 | AT5G17710.1 | s |
| Q9LHA8 | MD37C | Protein processing - chaperone | Chaperone | Psat3g182640.1 | AT3G12580.1 | s |
| O22265 | cpSRP43 | Protein processing - chaperone | Chaperone | Psat7g178840.1 | AT2G47450.1 | s, m |
| Q9FUZ2 * | DEF1B | Protein processing - deformylase | Peptide deformylase 1B,<br>substrate specificity towards PSII D1 polypeptide | Psat3g166600.1 | AT5G14660.1 | s |
| Q9FV52 | MAP1B | Protein processing - protease | N-terminal methionine excision | Psat2g085800.1 | AT1G13270.1 | s, l |
| P30184 * | AMPL1 | Protein processing - protease | Protease | Psat1g025200.2 | AT2G24200.1 | s, m |
| Q94AM1 * | OOPDA | Protein processing - protease | Protease | Psat2g133960.1 | AT5G65620.1 | s, l |
| Q8VZF3 * | CGEP | Protein processing - protease | Protease | Psat3g019800.3 | AT2G47390.1 | s |
| Q8VY06 * | PREP2 | Protein processing - protease | Protease | Psat4g213560.1 | AT1G49630.1 | s, m |
| Q9S834 | CLPP5 | Protein processing - protease | Protease | Psat6g161200.1 | AT1G02560.1 | s, m, l |
| O82261 * | DEGP2 | Protein processing - protease | Protease, primary cleavage of photodamaged D1 protein | Psat6g167960.1 | AT2G47940.1 | s, m |
| O64730 * | P2C26 | Protein processing - PTM | PSII phosphatase | Psat5g153360.1 | AT2G30170.1 | s, m, l |
| P29450 * | TRXF | Redox | | Psat3g102640.1 | AT5G16400 | s, l |
| Q9SEU6 * | TRXM4 | Redox | | Psat5g242680.1<br>Psat5g229000.1 * | AT3G15360.1 | l<br>s, l |
| Q9ZUC1 * | AOR | Redox | NADPH-dependent alkenal/one oxidoreductase | Psat4g123800.1 | AT1G23740.1 | l |
| Q8VYM4 | PSRP2 | Ribosome - chloroplast | CP Ribosome | Psat0s362g0040.1 | AT3G52150.1 | l |
| Q9BBP8 | RR3 | Ribosome - chloroplast | CP Ribosome | Psat4g046880.1 | ATCG00800.1 | l |
| P82191 | RK3 | Ribosome - chloroplast | CP Ribosome | Psat5g001240.1 | At2g43030.1 | l |
| A4GGC9 | RR12 | Ribosome - chloroplast | CP Ribosome | Psat6g178680.1 | ATCG00065.1 | l |
| Q10PV9 | RH47B | RNA processing | | Psat1g016000.1 | AT1G12770 | s, m, l |
| P19684 * | ROC5 | RNA processing | | Psat3g028520.1 | AT3G52380.1 | s, l |
| P28644 | ROC1 | RNA processing | | Psat5g050360.1 | AT5G50250 | s, m |
| Q84W56 * | RNJ | RNA processing | | Psat5g139840.1 | AT5G63420.1 | l |
| Q9C7Y2 * | MORF5 | RNA processing | mRNA processing | Psat6g105080.3 | AT1G32580.1 | m, l |
| Q69LE7 * | PNP1 | RNA processing | r/mRNA processing | Psat0s2078g0040.1 | AT3G03710.1 | s, m |
| Q9LU63 | ATP1 | Signaling | | Psat6g049600.1 | AT5G51110.1 | l |
| Q657X6 * | EXEC2 | Signaling | Together with EX1, executor protein of programmed cell death | Psat5g114480.1 | AT1G27510.1 | s, m, l |
| Q0ZJ28 | RPOB | Transcription | DNA-directed RNA polymerase subunit beta | Psat7g176880.1 | ATCG00190 | s |

| | | | | | | |
|---|---|---|---|---|---|---|
| A0A1S2Z662 | ICT1 | Translation | peptidyl-tRNA hydrolase ICT1, mitochondrial | Psat4g068640.1 | At5g19830.1 | s, m, l |
| Q9M1X0 | RRFC | Translation | Ribosome-recycling factor, chloroplastic | Psat6g060640.1 | AT3G63190.1 | m |
| Q8RX79 * | APG3 | Translation regulation | Peptide chain release factor | Psat3g023960.1 | AT3G62910.1 | m |
| O24310 | EFTU | Translation regulation | Translation | Psat5g116480.1 | At4g20360 | s |
| O82234 | IF32 | Translation regulation | Translation | Psat7g006440.1 | AT2G24060.1 | s, m |
| B9HQZ6 * | SYAP | tRNA processing - synthase | tRNA ligase | Psat5g000840.1<br>Psat5g000880.3 * | AT5G22800.1 | s, m, l<br>s, m, l |
| F4IFC5 * | SYTM2 | tRNA processing - synthase | tRNA ligase | Psat5g104480.1 | AT2G04842.1 | s, m, l |

**Supplemental Table 3 Ribosomal proteins of purified RNCs identified in MS/MS**

Identified in TST-D1 RNCs with short(s), medium (m), and long (l) peptide chains and TST-uS2c RNCs (S2), respectively. According to *Pisum sativum* accessions (Razor Psat), Uniport (Uniprot ID) and *Arabidopsis thaliana* (AT) accessions were assigned. Asterisks mark significantly enriched proteins.

**30S ribosomal proteins**

| Uniprot ID | Uniprot Name | Ban nomenclatur | RNC chains | Razor Psat | Potential AT |
|---|---|---|---|---|---|
| P29344 | RR1 | bS1c | s, m, l, S2 | Psat1g162760.1 | AT5G30510 |
| P08241 | RR2 | uS2c | s, S2 | Psat6g219440.1 | ATCG00160 |
| Q9BBP8 | RR3 | uS3c | s, m, l*, S2 | Psat4g046880.1 | ATCG00800 |
| Q9ST69 | RR5 | uS5c | s, m, l, S2 | Psat7g115360.1 | AT2G33800 |
| Q8VY91 | RR6 | bS6c | s, m, l, S2 | Psat7g023960.1 | AT1G64510 |
| Q6KGX3 | RR7 | uS7c | s, m, l, S2 | Psat5g119080.1 | AtCg00900 |
| P82278 | RR9 | uS9c | s, m, l, S2 | Psat6g010760.1 | AT1G74970 |
| Q9M4Y3 | RR10 | uS10c | s, m, l, S2 | Psat5g170120.1 | AT3G13120 |
| A4GGC9 | RR12 | uS12c | s, m, l*, S2 | Psat6g178680.1 | ATCG00065 |
| P42732 | RR13 | uS13c | s, m, l, S2 | Psat5g134240.2 | AT5G14320 |
| P82130 | RR20 | bS20c | s, m, l, S2 | Psat4g136440.1 | AT3G15190 |
| P82024 | RR21 | bS21c | s, m, l, S2 | Psat3g169680.2 | AT3G27160 |
| Q8VYM4 | PSRP2 | cS22 | s, m, l, S2<br>l | Psat0s362g0040.1<br>Psat0s2674g0040.2 | AT3G52150 |

**50S ribosomal proteins**

| Uniprot ID | Uniprot Name | Ban nomenclatur | RNC chains | Razor Psat | Potential AT |
|---|---|---|---|---|---|
| Q9LY66 | RK1 | uL1c | s, m, l, S2 | Psat7g200640.1 | AT3G63490 |
| P31163 | RK2 | uL2c | s, m, l, S2 | Psat5g168160.1 | ATCG00830<br>ATCG01310 |
| P82191 | RK3 | uL3c | s, m, l*, S2 | Psat5g001240.1 | AT2G43030 |
| O80361 | RK4 | uL4c | s, m, l, S2 | Psat4g224360.1 | AT1G07320 |
| O04603 | RK5 | uL5c | s, m, l, S2 | Psat3g107160.1 | AT4G01310 |
| O23049 | RK6 | uL6c | s, m, l, S2 | Psat1g066560.1 | AT1G05190 |
| P11894 | RK9 | bL9c | s, m, l, S2 | Psat7g223760.1 | AT3G44890 |
| O80362 | RK10 | uL10c | s, m, l, S2 | Psat4g007600.1 | AT5G13510 |
| Q9MAP3 | RK11 | uL11c | s, m, l, S2 | Psat5g110320.1 | AT1G32990 |
| P36210 | RK121 | bL12c | s, m, l, S2 | Psat1g056240.1 | AT3G27830 |
| Q9SYL9 | RK13 | uL13c | s, m, l, S2 | Psat4g140640.1 | AT1G78630 |
| B5LMQ9 | RK14 | uL14c | s, m, l, S2<br>m, S2 | Psat1g099160.1<br>Psat5g284200.1 | ATCG00780 |
| P31165 | RK15 | uL15c | s, m, l, S2 | Psat6g225000.1 | AT3G25920 |
| B5LMR0 | RK16 | uL16c | m, l, S2 | Psat0s1446g0200.1 | ATCG00790 |
| P82195 | RK18 | uL18c | m, S2 | Psat1g016560.1 | AT1G48350 |
| P82413 | RK19 | bL19c | s, m, l, S2 | Psat2g176160.2 | AT4G17560 |
| P51412 | RK21 | bL21c | s, m, l, S2 | Psat6g187560.1 | AT1G35680 |
| P23408 | RK22 | uL22c | s, m, l, S2 | Psat1g052200.1 | ATCG00810 |
| P11893 | RK24 | uL24c | s, m, l, S2 | Psat5g249200.1 | AT5G54600 |
| P82190 | RK27 | bL27c | s, m, l, S2 | Psat5g215240.1 | AT5G40950 |
| P82248 | RK29 | uL29c | s, m, l, S2 | Psat7g055240.1 | AT5G65220 |
| Q9FWS4 | RK31 | bL31c | s, m, l, S2 | Psat6g009240.1 | AT1G75350 |
| P15820 | RK32 | bL32c | l, S2 | Psat5g142800.1 | ATCG01020 |
| Q8VZ55 | RK35 | bL35c | s, m, l, S2 | Psat7g006560.1 | AT2G24090 |

**Supplemental Table 4 MaxQuant analyses MS/MS label-free quantification intensities**

| Uniprot ID | Uniprot Name | Razor Psat# | Potential At# | LFQ intensity TST-D1 (56)_1_A | LFQ intensity TST-D1 (56)_1_B | LFQ intensity TST-D1 (56)_2_A | LFQ intensity TST-D1 (56)_2_B | LFQ intensity TST-D1 (69)_1_A | LFQ intensity TST-D1 (69)_1_B | LFQ intensity TST-D1 (69)_2_A | LFQ intensity TST-D1 (69)_2_B | LFQ intensity TST-D1 (108)_1_A | LFQ intensity TST-D1 (108)_1_B | LFQ intensity TST-D1 (108)_2_A | LFQ intensity TST-D1 (108)_2_B | LFQ intensity TST-D1 (136)_1_A | LFQ intensity TST-D1 (136)_1_B | LFQ intensity TST-D1 (136)_2_A | LFQ intensity TST-D1 (136)_2_B | LFQ intensity TST-D1 (195)_1_A | LFQ intensity TST-D1 (195)_1_B | LFQ intensity TST-D1 (195)_2_A | LFQ intensity TST-D1 (195)_2_B | LFQ intensity TST-D1 (291)_1_A | LFQ intensity TST-D1 (291)_1_B | LFQ intensity TST-D1 (291)_2_A | LFQ intensity TST-D1 (291)_2_B | LFQ intensity TST-uS2c (158)_1_A | LFQ intensity TST-uS2c (158)_1_B | LFQ intensity TST-uS2c (158)_2_A | LFQ intensity TST-uS2c (158)_2_B |
|---|---|---|---|---|---|---|---|---|---|---|---|---|---|---|---|---|---|---|---|---|---|---|---|---|---|---|---|---|---|---|---|
| O81360 | ABA2 | Psat0s1281g0080.1 | AT5G67030.1 | 13515 | 9571,8 | 96655 | 169680 | 17405 | 25964 | 113830 | 263900 | 10309 | 8890,6 | 96050 | 105950 | 5715,5 | 0 | 0 | 18663 | 0 | 0 | 57992 | 5405,2 | 7581,5 | 0 | 3206,5 | 3854,1 | 4899,2 | 4741,7 | 0 | 7937,3 |
| P04717 | RBL | Psat0s1330g0280.1 | AtCg00490 | 38386000 | 34984000 | 3515000 | 3852100 | 49048000 | 48826000 | 4155700 | 4455500 | 78545000 | 77929000 | 3884800 | 2771900 | 75926000 | 59798000 | 3958200 | 4073100 | 31973000 | 31873000 | 3716500 | 4037200 | 42684000 | 28193000 | 2789200 | 2485100 | 3114100 | 4107200 | 0 | 2179700 |
| B5LMR0 | RK16 | Psat0s1446g0200.1 | AtCg00790.1 | 0 | 0 | 0 | 0 | 0 | 0 | 0 | 0 | 0 | 0 | 10855 | 13972 | 0 | 0 | 0 | 7441,3 | 0 | 0 | 8625,3 | 31036 | 0 | 0 | 7015,4 | 0 | 5612,7 | 0 | 0 | 18808 |
| Q9FGS0 | CP31B | Psat0s1670g0040.1 | AT5G50250.1 | 558240 | 466570 | 1034,8 | 2656,7 | 987670 | 616240 | 2927,6 | 1193,6 | 564020 | 216490 | 0 | 0 | 1046800 | 1521500 | 1217,1 | 10272 | 973890 | 535960 | 4583,1 | 3923,6 | 1032600 | 470200 | 0 | 0 | 3804,3 | 13307 | 0 | 1951 |
| Q42586 | UMPS | Psat0s1718g0040.2 | AT3G54470.1 | 0 | 0 | 15424 | 14220 | 0 | 0 | 17977 | 29499 | 0 | 0 | 16467 | 22625 | 6693,9 | 0 | 4892 | 0 | 0 | 8795,1 | 9997,2 | 10338 | 0 | 0 | 0 | 0 | 0 | 0 | 0 | 6214,1 |
| Q9SGD6 | AROD6 | Psat0s1945g0080.1 | AT1G08250.1 | 37453 | 42839 | 0 | 0 | 45074 | 31155 | 0 | 0 | 52180 | 66145 | 0 | 10710 | 33441 | 19569 | 0 | 0 | 33942 | 38512 | 0 | 0 | 32086 | 0 | 0 | 0 | 0 | 0 | 0 | 0 |
| Q41008 | ACCA | Psat0s1978g0280.1 | AT2G38040.1 | 0 | 0 | 22764 | 28562 | 0 | 0 | 46253 | 24945 | 17585 | 22862 | 21611 | 37480 | 0 | 26912 | 27885 | 30222 | 0 | 0 | 29796 | 39107 | 0 | 0 | 22250 | 41372 | 14514 | 0 | 0 | 13982 |
| Q69LE7 | PNP1 | Psat0s2078g0040.1 | AT3G03710.1 | 12135 | 19933 | 0 | 0 | 23800 | 20135 | 7130,2 | 5574 | 26072 | 21541 | 7319,6 | 0 | 0 | 30781 | 0 | 0 | 24064 | 18694 | 0 | 0 | 15264 | 0 | 0 | 0 | 0 | 0 | 0 | 0 |
| Q9C8P0 | ODP25 | Psat0s2229g0080.1 | AT1G34430.1 | 31904 | 32885 | 0 | 45327 | 29434 | 51331 | 49759 | 52158 | 41126 | 51181 | 13798 | 46919 | 25117 | 25304 | 52938 | 49552 | 0 | 0 | 13162 | 47947 | 98009 | 71650 | 25838 | 53163 | 9047,1 | 15597 | 0 | 33255 |
| O81004 | Y2287 | Psat0s2458g0040.1 | AT2G22870.1 | 0 | 0 | 83182 | 39528 | 18459 | 19318 | 82078 | 73621 | 0 | 0 | 53830 | 17161 | 10101 | 0 | 72072 | 45814 | 0 | 0 | 68959 | 18583 | 0 | 0 | 154360 | 63625 | 11328 | 0 | 0 | 54274 |
| Q8VYM4 | PSRP2 | Psat0s2674g0040.2 | AT3G52150.1 | 242120 | 329560 | 11822 | 16167 | 283120 | 369590 | 11799 | 31436 | 291680 | 217280 | 41947 | 79475 | 309330 | 130300 | 63618 | 65858 | 566810 | 531920 | 76562 | 263070 | 357790 | 410080 | 41629 | 13517 | 5944,3 | 19693 | 0 | 177780 |
| P18823 | ACCD | Psat0s6989g0080.1 | ATCG00500.1 | 41150 | 123460 | 97814 | 144630 | 24595 | 32692 | 268470 | 149420 | 0 | 41754 | 98154 | 109270 | 17985 | 7125,9 | 57070 | 40590 | 0 | 0 | 57657 | 47271 | 18356 | 0 | 14268 | 9245 | 23554 | 33240 | 0 | 51463 |
| P08215 | ATPA | Psat6g219360.1 | AtCg00120 | 168420 | 207590 | 580230 | 580570 | 242800 | 158810 | 812940 | 801580 | 243660 | 219700 | 427320 | 555160 | 112650 | 93174 | 130770 | 184220 | 122830 | 142360 | 260030 | 191090 | 74895 | 147840 | 142540 | 40110 | 169710 | 148080 | 0 | 153160 |
| Q8L7S8 | RH3 | Psat0s2843g0040.1 | AT5G26742.1 | 218560 | 284390 | 27006 | 25717 | 240270 | 381320 | 39542 | 35775 | 467760 | 341770 | 19817 | 24059 | 177580 | 186350 | 11392 | 23615 | 226140 | 190260 | 18029 | 52103 | 320040 | 176110 | 17860 | 0 | 23817 | 4055,2 | 0 | 21871 |
| Q9SZX3 | ASSY | Psat0s3176g0120.1 | AT4G24830.1 | 0 | 0 | 17176 | 0 | 0 | 5315,7 | 28171 | 22575 | 0 | 7712,9 | 0 | 14587 | 4511,8 | 0 | 0 | 11830 | 5929,6 | 0 | 0 | 13829 | 0 | 6465,2 | 0 | 0 | 0 | 0 | 0 | 0 |
| Q8H0V1 | CK5P1 | Psat0s3355g0160.1 | AT4G36390.1 | 16664 | 0 | 9288,5 | 10891 | 11646 | 12233 | 14044 | 16419 | 16646 | 16151 | 9864,1 | 17873 | 8770,2 | 0 | 8811,5 | 9822,4 | 0 | 0 | 11432 | 17095 | 0 | 16887 | 0 | 0 | 0 | 14369 | 0 | 6668,7 |
| O24457 | ODPA3 | Psat0s3525g0080.1 | AT1G01090.1 | 12123 | 10656 | 0 | 0 | 14762 | 21534 | 21485 | 0 | 81512 | 29162 | 0 | 0 | 20213 | 0 | 0 | 0 | 31182 | 22012 | 0 | 0 | 27037 | 33714 | 0 | 0 | 2288,6 | 0 | 0 | 0 |
| O22437 | CHLD | Psat0s3604g0040.1 | AT1G08520.1 | 18120 | 15757 | 213800 | 98873 | 16074 | 26175 | 439880 | 830230 | 12012 | 0 | 134980 | 152940 | 21416 | 17749 | 83404 | 77633 | 13949 | 0 | 54610 | 468200 | 13953 | 0 | 10626 | 33451 | 28485 | 26339 | 0 | 24710 |
| Q8VYM4 | PSRP2 | Psat0s362g0040.1 | AT3G52150.1 | 18049 | 13734 | 0 | 0 | 16651 | 14245 | 0 | 0 | 0 | 14018 | 0 | 0 | 46095 | 0 | 0 | 0 | 22235 | 71582 | 0 | 10832 | 0 | 0 | 0 | 0 | 0 | 0 | 0 | 0 |
| P31163 | RK2 | Psat5g168160.1 | AtCg00830.1<br>AtCg01310.1 | 3942 | 4602,5 | 3704,9 | 3721,2 | 4771,9 | 3624,3 | 3215,3 | 4340,9 | 4697,5 | 8501,6 | 9060,9 | 16492 | 42960 | 4802,4 | 23383 | 5197,1 | 4149,7 | 0 | 161820 | 52734 | 1867,1 | 4101,5 | 40871 | 14703 | 72112 | 173660 | 0 | 244420 |
| Q8S9M1 | PAP13 | Psat0s4417g0040.1 | AT2G42130.1 | 26868 | 11300 | 13094 | 7253,6 | 23146 | 11668 | 10961 | 6567,3 | 9205,1 | 0 | 11411 | 5385 | 9015,1 | 4642,8 | 0 | 0 | 10495 | 8829,6 | 9509,4 | 0 | 11137 | 14392 | 0 | 0 | 18417 | 7835,7 | 0 | 0 |
| P36774 | LON2 | Psat0s471g0440.1 | AT1G75460.1 | 9174,7 | 8282,1 | 78416 | 44799 | 10897 | 11683 | 36441 | 54209 | 16026 | 17199 | 53597 | 67459 | 8086,5 | 10119 | 15893 | 0 | 0 | 0 | 15590 | 23781 | 0 | 0 | 62206 | 83043 | 53493 | 58630 | 0 | 11137 |
| P06585 | PSBA | Psat0ss8367g0080.1 | ATCG00020.1 | 24193 | 14301 | 29491 | 34991 | 25235 | 29384 | 64441 | 99546 | 3128,4 | 30762 | 72809 | 138360 | 31754 | 12283 | 65023 | 57255 | 39962 | 56697 | 58946 | 220330 | 41845 | 22432 | 0 | 9393,8 | 0 | 0 | 0 | 0 |
| P04717 | RBL | Psat0s5127g0120.1 | AtCg00490 | 3045800 | 2077400 | 448530 | 287730 | 378840 | 281550 | 662060 | 424670 | 2799400 | 201940 | 327730 | 268290 | 1635100 | 2324500 | 679920 | 493010 | 731980 | 218750 | 347930 | 201580 | 1635200 | 2154700 | 145780 | 218000 | 407190 | 282770 | 0 | 111730 |
| Q6DYE4 | Y1609 | Psat0s629g0200.5 | AT1G26090.1 | 1843,3 | 0 | 0 | 0 | 0 | 0 | 0 | 0 | 2301,7 | 1934,6 | 0 | 0 | 2515,2 | 0 | 0 | 0 | 0 | 2475,6 | 0 | 1172,3 | 1910,5 | 0 | 0 | 0 | 0 | 0 | 0 | 0 |
| Q0ZJ28 | RPOB | Psat7g176880.1 | ATCG00190 | 6609,1 | 7142,4 | 2875,5 | 4030,3 | 6351 | 6204,5 | 3565,2 | 4437,1 | 9815,2 | 7371,3 | 4211,3 | 0 | 0 | 0 | 0 | 0 | 0 | 0 | 0 | 0 | 5373,1 | 6378,1 | 0 | 0 | 0 | 0 | 0 | 1328,4 |
| Q9LLC1 | BCCP2 | Psat0s846g0040.1 | AT5G15530.1 | 689050 | 794420 | 4862400 | 3482800 | 780710 | 875840 | 7710300 | 7284500 | 936780 | 1214700 | 3882800 | 3944200 | 802080 | 824640 | 8264000 | 6326500 | 2147500 | 1589000 | 4835500 | 4351100 | 1086200 | 1177000 | 5410900 | 6272800 | 2309600 | 3010400 | 0 | 3024800 |
| B9HBA8 | ACCC1 | Psat0s916g0080.1 | AT5G35360 | 62750000 | 65643000 | 351250000 | 263200000 | 75934000 | 75017000 | 352070000 | 302550000 | 125620000 | 122310000 | 347120000 | 280910000 | 50205000 | 53862000 | 478430000 | 472540000 | 52318000 | 47805000 | 340060000 | 259900000 | 62251000 | 57213000 | 654960000 | 729840000 | 188250000 | 173860000 | 2089600 | 194290000 |
| P08927 | RUBB | Psat1g001680.1 | AT1G55490 | 11165000 | 9261900 | 16745000 | 17765000 | 13346000 | 11869000 | 19216000 | 13635000 | 15460000 | 14703000 | 17433000 | 17894000 | 11817000 | 12399000 | 12990000 | 12077000 | 12300000 | 10445000 | 15828000 | 10157000 | 8333400 | 8689700 | 3206000 | 4226200 | 6718800 | 4716800 | 0 | 7267300 |
| Q9SAJ6 | G3PP1 | Psat1g005000.1 | AT1G79530.1 | 15925 | 14128 | 0 | 0 | 27399 | 64654 | 2186,3 | 2246,4 | 30717 | 25037 | 0 | 0 | 50604 | 46129 | 0 | 0 | 93296 | 92232 | 0 | 0 | 51322 | 26079 | 0 | 0 | 0 | 0 | 0 | 0 |
| Q42961 | PGKH | Psat1g014040.1 | AT1G56190 | 134540 | 194770 | 281340 | 144100 | 142580 | 126090 | 335290 | 162970 | 175710 | 252390 | 146690 | 53805 | 1093200 | 1115900 | 24065 | 77676 | 330020 | 315130 | 66369 | 121890 | 478480 | 215570 | 0 | 252110 | 4768,7 | 63933 | 0 | 21175 |
| Q10PV9 | RH47B | Psat1g016000.1 | AT1G12770 | 22685 | 20690 | 21011 | 13864 | 65366 | 43761 | 22723 | 35888 | 32277 | 34675 | 30516 | 8725,5 | 7989,9 | 12676 | 0 | 16931 | 19523 | 12733 | 9672,8 | 23903 | 0 | 0 | 10132 | 0 | 0 | 0 | 0 | 15432 |
| P82195 | RK18 | Psat1g016560.1 | AT1G48350.1 | 0 | 0 | 0 | 0 | 0 | 7080,5 | 0 | 0 | 0 | 0 | 0 | 0 | 3162,9 | 4754,3 | 0 | 23792 | 0 | 0 | 24225 | 0 | 0 | 0 | 9863,3 | 0 | 0 | 0 | 0 | 21176 |
| O82230 | STIC2 | Psat1g022480.1 | AT2G24020.1 | 12271 | 25196 | 32274 | 46889 | 28216 | 0 | 15073 | 17053 | 13538 | 0 | 20309 | 22201 | 38094 | 59789 | 25267 | 34639 | 19747 | 35105 | 14444 | 22733 | 36896 | 38495 | 0 | 0 | 0 | 0 | 0 | 0 |
| P30184 | AMPL1 | Psat1g025200.2 | AT2G24200.1 | 25047 | 28766 | 0 | 0 | 26518 | 27604 | 0 | 3801,8 | 21564 | 28137 | 5350,9 | 0 | 52053 | 45974 | 0 | 8579,6 | 47424 | 18434 | 0 | 0 | 48420 | 40102 | 0 | 0 | 0 | 0 | 0 | 0 |
| Q6STH5 | HF101 | Psat1g039560.1 | AT3G24430.1 | 6393,1 | 48495 | 272000 | 734100 | 23037 | 36261 | 589270 | 931430 | 35179 | 49170 | 265970 | 472470 | 8062,6 | 16168 | 27403 | 12409 | 23709 | 17651 | 10509 | 54750 | 3746 | 13113 | 0 | 0 | 3880,4 | 0 | 0 | 3034,8 |
| A3PCG2 | EFTS | Psat1g044600.1 | AT4G29060 | 151650 | 210270 | 56852 | 76164 | 184630 | 173490 | 28264 | 54741 | 328680 | 281840 | 37684 | 36713 | 337820 | 378240 | 50524 | 61860 | 538720 | 591280 | 93888 | 186360 | 464170 | 659380 | 28562 | 21739 | 27304 | 8170,9 | 0 | 96299 |
| P23408 | RK22 | Psat1g052200.1 | ATCG00810.1 | 0 | 0 | 27290 | 0 | 0 | 0 | 0 | 0 | 0 | 0 | 12127 | 29299 | 33399 | 30496 | 0 | 9334,5 | 34572 | 32077 | 17042 | 130300 | 0 | 74532 | 6814,3 | 5522,7 | 8640,6 | 10527 | 0 | 34686 |
| P36210 | RK121 | Psat1g056240.1 | AT3G27830.1 | 357850 | 285330 | 465980 | 536410 | 146110 | 501220 | 527870 | 472510 | 214360 | 182680 | 1458400 | 900830 | 1743300 | 1009900 | 1141800 | 865000 | 527600 | 214110 | 2769700 | 1113100 | 238280 | 218410 | 473310 | 405000 | 441800 | 438250 | 0 | 2081800 |
| P17067 | CAHC | Psat1g058960.5 | AT3G01500 | 489290 | 377130 | 153290 | 94885 | 303690 | 265220 | 213530 | 95678 | 658060 | 861220 | 161330 | 59628 | 867210 | 755090 | 68070 | 66653 | 368320 | 277710 | 73426 | 38483 | 156840 | 119070 | 63931 | 48547 | 31540 | 31402 | 0 | 41405 |
| O23049 | RK6 | Psat1g066560.1 | AT1G05190.1 | 6919,7 | 9906,8 | 32058 | 8390,7 | 12942 | 0 | 17915 | 23568 | 269640 | 53282 | 84983 | 22916 | 33093 | 17798 | 37288 | 36965 | 17216 | 16397 | 125530 | 212140 | 52767 | 0 | 51867 | 32500 | 43385 | 98668 | 0 | 175920 |
| Q9LIB2 | PHS1 | Psat1g066720.1 | AT3G29320.1 | 12628 | 10160 | 0 | 0 | 10685 | 11530 | 0 | 0 | 15025 | 19650 | 29187 | 0 | 19514 | 10388 | 0 | 0 | 6347,1 | 0 | 0 | 0 | 0 | 0 | 0 | 0 | 0 | 0 | 0 | 0 |
| Q42533 | BCCP1 | Psat1g067360.1 | AT5G16390.1 | 1210500 | 660080 | 1020600 | 1286200 | 2124800 | 1185400 | 3416500 | 2738600 | 1467700 | 1353900 | 2551700 | 4506100 | 819810 | 857910 | 7315900 | 5833900 | 2544000 | 646030 | 3160500 | 1972000 | 3083000 | 2043600 | 11553000 | 5138700 | 711270 | 816130 | 0 | 1347800 |
| Q9LF37 | CLPB3 | Psat1g069480.1 | AT5G15450.1 | 32752 | 28002 | 0 | 0 | 37004 | 46454 | 3250,8 | 4479,8 | 44287 | 69837 | 2949,8 | 0 | 53008 | 30506 | 0 | 0 | 41426 | 63057 | 0 | 0 | 21833 | 16768 | 0 | 0 | 0 | 0 | 0 | 0 |
| A0A1S2XK40 | A0A1S2XK40 | Psat1g069960.2 |  | 1274,7 | 1193,4 | 0 | 0 | 0 | 1184,7 | 1433,6 | 0 | 1404,1 | 0 | 0 | 0 | 0 | 2208,5 | 0 | 0 | 3254,3 | 5128,8 | 0 | 0 | 2485 | 4181,5 | 0 | 0 | 0 | 0 | 0 | 0 |
| O34331 | YLBH | Psat1g076480.1 |  | 3492,7 | 3740,6 | 3421 | 0 | 3625,1 | 3416 | 3324,1 | 4018,3 | 3604,8 | 3098 | 0 | 0 | 4315 | 3904,4 | 2566,6 | 3121,7 | 4862,3 | 0 | 3257,2 | 0 | 3615,7 | 4750,8 | 0 | 0 | 2754 | 0 | 0 | 4451,7 |
| A0A2Z6NLS5 | A0A2Z6NLS5 | Psat1g080040.1 |  | 36614 | 30528 | 10401 | 39410 | 31293 | 33035 | 4477,8 | 10044 | 35698 | 38983 | 35655 | 24893 | 129730 | 88485 | 15632 | 23671 | 177440 | 209020 | 61777 | 88934 | 252440 | 92636 | 10728 | 11430 | 18674 | 15349 | 0 | 70395 |
| O65194 | RBS | Psat1g080440.1 | AT1G67090 | 2681800 | 2225300 | 255710 | 256220 | 2062900 | 2071700 | 127440 | 118740 | 2200100 | 2216400 | 110130 | 2399,3 | 3314600 | 3120900 | 104530 | 109860 | 1200800 | 1256300 | 178280 | 29327 | 2037800 | 1615200 | 30218 | 29442 | 67897 | 98810 | 0 | 96067 |
| Q9FLW9 | PKP2 | Psat1g096960.1 | AT5G52920.1 | 1207900 | 948980 | 265100 | 300760 | 658750 | 894420 | 497460 | 418410 | 999060 | 727660 | 409250 | 380350 | 554980 | 265350 | 446200 | 335280 | 389360 | 308850 | 361120 | 137840 | 359150 | 279800 | 402360 | 379850 | 169480 | 126660 | 0 | 119100 |
| B5LMQ9 | RK14 | Psat1g099160.1 | AtCg00780.1 | 0 | 14498 | 19968 | 20054 | 0 | 14934 | 12995 | 21176 | 22309 | 0 | 38043 | 36609 | 16311 | 21819 | 17567 | 16689 | 25354 | 25526 | 124780 | 44574 | 29923 | 23101 | 19328 | 11129 | 17650 | 18059 | 0 | 69627 |
| Q9LP77 | Y1848 | Psat1g118000.1 | AT1G48480.1 | 0 | 0 | 0 | 0 | 0 | 0 | 0 | 0 | 0 | 0 | 0 | 0 | 0 | 0 | 0 | 0 | 0 | 0 | 0 | 0 | 0 | 0 | 0 | 0 | 14899 | 13573 | 0 | 23345 |
| A0A072V684 | A0A072V684 | Psat1g134240.1 |  | 5415,6 | 4736,3 | 0 | 7650,7 | 0 | 0 | 0 | 0 | 0 | 0 | 0 | 0 | 4663,6 | 5616,6 | 0 | 0 | 0 | 0 | 0 | 0 | 0 | 0 | 0 | 0 | 0 | 0 | 0 | 0 |
| O65282 | CH20 | Psat1g135800.1 | AT5G20720.1 | 600680 | 534600 | 176160 | 125530 | 562560 | 382150 | 140210 | 174090 | 368130 | 335380 | 323430 | 227530 | 1004400 | 1110200 | 141760 | 153120 | 582110 | 279020 | 246770 | 71713 | 309390 | 265740 | 52587 | 0 | 180880 | 65492 | 0 | 137600 |
| P29344 | RR1 | Psat1g162760.1 | At5g30510.1 | 98476 | 146380 | 219450 | 223800 | 94735 | 111750 | 383370 | 438800 | 147130 | 93606 | 497740 | 510580 | 244910 | 195040 | 431430 | 384200 | 101770 | 157330 | 841390 | 1048100 | 169810 | 237510 | 219070 | 299160 | 204870 | 201530 | 0 | 827450 |
| Q84WN0 | Y4920 | Psat1g169560.1 | AT4G37920.1 | 0 | 7371,6 | 39038 | 34594 | 0 | 9177,2 | 42207 | 47468 | 10912 | 0 | 22491 | 79748 | 6894 | 0 | 0 | 0 | 0 | 8024,8 | 10674 | 36009 | 0 | 0 | 0 | 0 | 7286,6 | 0 | 0 | 0 |
| P08281 | GLNA2 | Psat1g185960.3 | AT5G35630 | 432150 | 491400 | 0 | 0 | 520780 | 475650 | 0 | 0 | 699170 | 731850 | 0 | 0 | 850010 | 989470 | 5484 | 4812,3 | 975640 | 676160 | 0 | 0 | 734940 | 826010 | 2608,3 | 0 | 0 | 30814 | 0 | 0 |
| Q1H537 | DCVR | Psat1g187280.1 | AT5G18660.1 | 0 | 11715 | 136670 | 97223 | 15171 | 0 | 52027 | 48590 | 19759 | 10129 | 32119 | 71539 | 0 | 0 | 13644 | 15423 | 0 | 0 | 27941 | 0 | 0 | 11271 | 0 | 10004 | 0 | 0 | 0 | 5622,1 |
| Q94JQ3 | GLYP3 | Psat1g192400.2 | AT4G32520.1 | 14302 | 13389 | 3204,2 | 3423,8 | 13056 | 13777 | 3141,5 | 2844,6 | 20735 | 29378 | 2659,6 | 1693,8 | 24190 | 28174 | 4413,2 | 4601,4 | 10796 | 11540 | 0 | 0 | 0 | 8492,2 | 3215,2 | 4136,3 | 0 | 0 | 0 | 1857,9 |
| P93162 | CHLI | Psat1g200160.2 | At4g18480.1 | 490320 | 302300 | 1500000 | 1020600 | 291000 | 253820 | 2041800 | 1462800 | 500090 | 149640 | 1462700 | 1263100 | 199850 | 238550 | 345940 | 292160 | 99516 | 98546 | 362710 | 200890 | 74476 | 33709 | 20977 | 56042 | 113060 | 123910 | 0 | 134640 |
| Q9M0Y6 | DPNPM | Psat1g209080.1 | AT4G05090.1 | 0 | 0 | 39436 | 33449 | 29457 | 0 | 43317 | 39377 | 26166 | 0 | 0 | 46802 | 30747 | 39418 | 0 | 0 | 0 | 32716 | 0 | 33036 | 0 | 42176 | 0 | 0 | 0 | 18708 | 0 | 0 |
| Q9LXC9 | IPYR6 | Psat1g212360.1 | AT5G09650.1 | 20886 | 8876,9 | 2579,6 | 0 | 7572,7 | 0 | 0 | 0 | 0 | 0 | 0 | 0 | 124660 | 67740 | 0 | 0 | 47327 | 30201 | 0 | 0 | 14984 | 11489 | 0 | 0 | 0 | 0 | 0 | 3100 |
| Q00218 | AROG | Psat1g216320.1 | AT4G33510.1 | 72696 | 87210 | 51962 | 43476 | 61632 | 94268 | 60073 | 55869 | 275960 | 199920 | 103730 | 64764 | 91068 | 101390 | 23719 | 23222 | 44865 | 68008 | 45958 | 12021 | 43388 | 65855 | 8789,1 | 0 | 20162 | 7637 | 0 | 8421,2 |
| Q02028 | HSP7S | Psat1g222720.1 | AT5G49910.1 | 1188400 | 1387600 | 1335300 | 1355300 | 1395200 | 1556400 | 1090600 | 1125300 | 1343400 | 1231000 | 832020 | 1193600 | 988160 | 1577500 | 1096900 | 1114300 | 714780 | 620070 | 578420 | 1069000 | 466090 | 1175200 | 338190 | 233050 | 1455800 | 1198900 | 0 | 1017200 |
| Q02028 | HSP7S | Psat1g222760.1 | AT5G49910.1 | 3397500 | 3045900 | 3921700 | 3848400 | 4070800 | 3176100 | 4967900 | 3341700 | 5936100 | 4611200 | 2674100 | 2157200 | 3496800 | 3672800 | 3327700 | 3695100 | 3023400 | 2425800 | 2264700 | 1282000 | 1839500 | 1742900 | 2405900 | 2280600 | 3722700 | 3651500 | 0 | 4162000 |
| Q8GSJ1 | HIS1B | Psat1g223400.1 | AT1G09795.1 | 10218 | 16618 | 0 | 4190,9 | 13532 | 12068 | 7222,9 | 0 | 21605 | 23272 | 0 | 0 | 14665 | 16942 | 0 | 0 | 11061 | 0 | 0 | 0 | 0 | 0 | 0 | 0 | 0 | 0 | 0 | 0 |
| Q8L7R2 | KHSE | Psat2g000800.1 | AT2G17265.1 | 48043 | 40466 | 26613 | 17107 | 50967 | 75495 | 21759 | 37742 | 53511 | 48529 | 7265,7 | 25872 | 54058 | 99055 | 54691 | 63163 | 51740 | 130000 | 15547 | 52089 | 141320 | 64714 | 11247 | 13964 | 0 | 0 | 0 | 20048 |
| P35055 | HEM6 | Psat2g001760.1 | AT1G03475.1 | 20093 | 29689 | 0 | 4643,8 | 23060 | 22573 | 0 | 5786,9 | 35991 | 48922 | 3556 | 0 | 212910 | 94400 | 0 | 0 | 42435 | 21700 | 0 | 0 | 27449 | 9290,3 | 0 | 0 | 0 | 0 | 0 | 0 |
| Q8S9L5 | TIG | Psat2g009800.1 | AT5G55220.1 | 38821 | 54152 | 34879 | 35664 | 66296 | 60610 | 20817 | 9016 | 107280 | 110080 | 35246 | 6010 | 171720 | 88666 | 0 | 0 | 104480 | 126830 | 0 | 6442,6 | 107770 | 119300 | 0 | 3491,8 | 9461,1 | 0 | 0 | 15619 |
| Q42545 | FTSZ1 | Psat2g010320.1 | AT5G55280.1 | 6292,6 | 7398,8 | 13988 | 6866 | 20837 | 16972 | 2060,1 | 10231 | 6350,9 | 7423 | 0 | 0 | 19187 | 18192 | 0 | 0 | 20928 | 16334 | 0 | 7536,9 | 18075 | 19607 | 0 | 0 | 0 | 0 | 0 | 0 |
| P54150 | MSRA4 | Psat2g010960.3 | AT4G25130.1 | 11801 | 11250 | 0 | 0 | 10342 | 9149 | 0 | 0 | 14144 | 14462 | 4394,8 | 0 | 13320 | 10268 | 0 | 0 | 9272,2 | 11146 | 0 | 0 | 7704,6 | 0 | 0 | 0 | 0 | 0 | 0 | 2977,5 |
| Q06881 | BCCP | Psat2g029880.1 | AT5G15530 | 415400 | 450420 | 4961600 | 3528500 | 425930 | 496830 | 5806400 | 4111700 | 589460 | 879040 | 3451500 | 4641500 | 411550 | 329560 | 4483000 | 6307200 | 1008300 | 767790 | 3494900 | 3751900 | 612280 | 759610 | 7449400 | 6062600 | 2268600 | 2748200 | 0 | 2720000 |
| Q9SKN5 | ARFJ | Psat2g033160.1 | AT2G28350.1 | 0 | 0 | 0 | 5527,6 | 9380,6 | 0 | 0 | 0 | 5290,6 | 0 | 0 | 7132,2 | 0 | 0 | 0 | 10924 | 0 | 0 | 0 | 0 | 0 | 0 | 9440,1 | 0 | 0 | 0 | 0 | 0 |
| Q7X9A0 | RCA1 | Psat2g039960.1 | AT2G39730.1 | 951090 | 815990 | 219720 | 171220 | 799860 | 966250 | 134090 | 210280 | 1645300 | 1478300 | 136370 | 144590 | 920670 | 1152500 | 204490 | 72140 | 1164700 | 943250 | 125590 | 80727 | 720090 | 754330 | 37003 | 19001 | 16908 | 24727 | 0 | 53664 |
| P08241 | RR2 | Psat6g219440.1 | ATCG00160 | 0 | 0 | 2314,4 | 0 | 2708,7 | 0 | 3005,4 | 0 | 0 | 0 | 0 | 0 | 0 | 0 | 0 | 0 | 0 | 0 | 5140,9 | 0 | 0 | 0 | 0 | 0 | 0 | 0 | 0 | 6508,7 |
| Q9ASX5 | Y5520 | Psat2g061840.1 | AT5G05200.1 | 0 | 0 | 4804 | 7799,9 | 0 | 3621,1 | 5256,9 | 5139,7 | 0 | 0 | 0 | 0 | 0 | 0 | 0 | 0 | 0 | 0 | 4764,2 | 0 | 0 | 0 | 0 | 0 | 0 | 0 | 0 | 0 |
| O80934 | Y2766 | Psat2g068960.2 | AT2G37660.1 | 180860 | 212140 | 553290 | 573100 | 224160 | 227610 | 902790 | 1180200 | 233570 | 268340 | 828330 | 791160 | 170320 | 185190 | 1064300 | 1078800 | 253970 | 196620 | 607580 | 782910 | 425800 | 207910 | 496720 | 651630 | 365610 | 249330 | 0 | 466540 |
| Q9C667 | CPNB4 | Psat2g080360.1 | AT1G26230.1 | 0 | 0 | 3445,8 | 4473,5 | 0 | 0 | 5368,8 | 2621,5 | 4263,5 | 0 | 0 | 4792,7 | 0 | 0 | 0 | 0 | 0 | 0 | 3245,8 | 0 | 0 | 0 | 0 | 0 | 0 | 0 | 0 | 2628,9 |
| Q9FV52 | MAP1B | Psat2g085800.1 | AT1G13270.1 | 47658 | 120440 | 23896 | 29029 | 29091 | 38931 | 40702 | 12456 | 31771 | 36498 | 21185 | 7879,1 | 18914 | 33306 | 26438 | 13647 | 28250 | 28690 | 25850 | 32376 | 105130 | 43546 | 8511 | 0 | 16196 | 0 | 0 | 16332 |
| G7JYB8 | G7JYB8 | Psat2g091800.1 |  | 6016,7 | 6389,2 | 12292 | 9339,9 | 9074 | 6522,3 | 11867 | 8662,8 | 0 | 7855,1 | 9854,2 | 0 | 12141 | 0 | 8503,7 | 8752,1 | 0 | 0 | 19209 | 4332,9 | 0 | 0 | 0 | 0 | 0 | 5861,1 | 0 | 15927 |
| Q9SIF2 | HS905 | Psat2g102720.1 | AT2G04030.1 | 656270 | 402260 | 692090 | 521350 | 392010 | 371590 | 877410 | 385640 | 975450 | 1026200 | 207250 | 217680 | 791720 | 804700 | 109160 | 135810 | 702380 | 501180 | 271460 | 226560 | 609290 | 432840 | 0 | 29396 | 49845 | 19240 | 0 | 23852 |
| P08215 | ATPA | Psat2g111080.1 | AtCg00120 | 43255 | 17574 | 89156 | 114940 | 15288 | 21317 | 76253 | 160260 | 45109 | 17507 | 99866 | 146020 | 20576 | 12429 | 36090 | 39663 | 8102,5 | 0 | 61802 | 21824 | 9078,1 | 4624,5 | 0 | 0 | 0 | 0 | 0 | 23169 |
| Q94AM1 | OOPDA | Psat2g133960.1 | AT5G65620.1 | 0 | 2848,4 | 0 | 0 | 3090,4 | 1960,7 | 0 | 1580,9 | 0 | 0 | 0 | 0 | 1424,4 | 3851,8 | 0 | 0 | 3836,2 | 4072,2 | 1577,2 | 0 | 0 | 0 | 0 | 0 | 0 | 0 | 0 | 0 |
| P05037 | ATPB | Psat2g136800.1 | ATCG00480.1 | 8687,7 | 16957 | 687770 | 254120 | 22303 | 23083 | 1125700 | 1159600 | 20353 | 32633 | 690270 | 198820 | 18407 | 14131 | 111400 | 142340 | 0 | 0 | 97892 | 107240 | 0 | 0 | 30166 | 41084 | 19232 | 31557 | 0 | 28632 |
| Q9ZP40 | PG1 | Psat2g138880.1 | AT2G35490 | 18869 | 18149 | 19474 | 14111 | 18159 | 24504 | 57010 | 38191 | 0 | 20304 | 31611 | 0 | 11255 | 9817,7 | 18735 | 0 | 25570 | 29310 | 17291 | 22259 | 50453 | 31550 | 0 | 0 | 0 | 0 | 0 | 12277 |
| P26302 | KPPR | Psat2g140680.1 | AT1G32060.1 | 33396 | 38130 | 0 | 0 | 42089 | 44667 | 4479,1 | 0 | 44289 | 54185 | 0 | 0 | 69949 | 78460 | 0 | 0 | 45467 | 38731 | 0 | 0 | 40612 | 28367 | 0 | 0 | 0 | 0 | 0 | 0 |
| Q94B60 | CLPP4 | Psat2g142240.1 | AT5G45390.1 | 288220 | 387660 | 10505 | 8649,5 | 66488 | 420510 | 3771,9 | 4111,9 | 272040 | 314550 | 4155,5 | 1448,4 | 63476 | 260730 | 0 | 0 | 477860 | 106800 | 0 | 3757,3 | 71211 | 69732 | 1246,6 | 0 | 0 | 1347,7 | 0 | 0 |
| P10933 | FENR1 | Psat2g154080.2 | AT5G66190.1 | 171670 | 118460 | 69569 | 47152 | 146260 | 105760 | 185740 | 73984 | 81534 | 62444 | 44909 | 36005 | 410290 | 214840 | 31195 | 30642 | 48690 | 83796 | 29928 | 32013 | 219920 | 84865 | 14424 | 28496 | 23043 | 23700 | 0 | 30357 |
| Q9LQK7 | AB7I | Psat2g173560.2 | AT1G32500.1 | 8782,8 | 8548,8 | 2320,2 | 0 | 7838,7 | 12057 | 3492,4 | 3925,2 | 16149 | 13686 | 4369,9 | 3221,2 | 13276 | 8902,4 | 2337,3 | 0 | 15030 | 13932 | 1866,2 | 16840 | 84794 | 31339 | 0 | 0 | 0 | 0 | 0 | 0 |
| P82413 | RK19 | Psat2g176160.2 | AT4G17560.1 | 3777,1 | 4485,3 | 10800 | 24837 | 5755,9 | 7687,3 | 29490 | 29702 | 16069 | 14904 | 60948 | 177610 | 4113,6 | 9343,1 | 146120 | 143720 | 11985 | 10204 | 173250 | 290270 | 5146,9 | 11282 | 28926 | 28420 | 145540 | 134860 | 0 | 497730 |
| Q40983 | SPP | Psat2g189160.1 | AT5G42390 | 0 | 0 | 3083,9 | 0 | 0 | 0 | 0 | 0 | 0 | 11566 | 0 | 4054 | 0 | 4155,2 | 0 | 0 | 3610,5 | 0 | 0 | 4345,6 | 2909,4 | 0 | 0 | 0 | 1981,5 | 2451,5 | 0 | 2828,2 |
| B9FSC8 | OPR11 | Psat2g190280.2 | AT1G76690.1 | 0 | 2813,4 | 0 | 0 | 2766,5 | 2522 | 0 | 2984,5 | 4467,9 | 0 | 0 | 0 | 0 | 5911,2 | 0 | 0 | 7331,3 | 7828,5 | 0 | 1900,7 | 0 | 0 | 0 | 0 | 0 | 0 | 0 | 0 |
| Q9LUJ3 | RDM1 | Psat2g190480.2 | AT3G22680.1 | 5785,4 | 0 | 32567 | 20611 | 0 | 7614,1 | 26273 | 0 | 0 | 0 | 20060 | 16787 | 0 | 0 | 17235 | 16016 | 0 | 0 | 17495 | 28802 | 0 | 0 | 15512 | 9364,1 | 0 | 0 | 0 | 12196 |
| Q9SA52 | CP41B | Psat3g014320.1 | AT1G09340.1 | 47597 | 56275 | 0 | 0 | 62599 | 49823 | 0 | 0 | 99122 | 94626 | 0 | 0 | 101640 | 190230 | 0 | 0 | 81570 | 112490 | 0 | 14424 | 55913 | 42143 | 0 | 0 | 12333 | 0 | 0 | 0 |

| | | | | | | | | | | | | | | | | | | | | | | | | | | | | | | | |
|---|---|---|---|---|---|---|---|---|---|---|---|---|---|---|---|---|---|---|---|---|---|---|---|---|---|---|---|---|---|---|---|
| Q8L7S8 | RH3 | Psat3g015480.1 | AT5G26742.1 | 82713 | 61747 | 90681 | 71710 | 115640 | 154070 | 64268 | 0 | 242750 | 109680 | 77509 | 54514 | 94024 | 81864 | 88675 | 72756 | 45501 | 63516 | 39165 | 14746 | 52970 | 44986 | 6422,6 | 0 | 100700 | 111130 | 0 | 120750 |
| Q8VZF3 | CGEP | Psat3g019800.3 | AT2G47390.1 | 8290,2 | 8591,7 | 0 | 0 | 14103 | 7865,4 | 0 | 2979,7 | 7755,7 | 19544 | 0 | 0 | 9805,6 | 9776 | 0 | 0 | 24000 | 0 | 0 | 0 | 0 | 10427 | 0 | 0 | 0 | 0 | 0 | 0 |
| Q8RX79 | APG3 | Psat3g023960.1 | AT3G62910.1 | 0 | 5514,1 | 0 | 0 | 6186,4 | 0 | 0 | 0 | 0 | 7601,5 | 0 | 3785,4 | 5450,7 | 5956,4 | 0 | 0 | 0 | 0 | 0 | 0 | 0 | 0 | 0 | 0 | 0 | 0 | 0 | 0 |
| P19684 | ROC5 | Psat3g028520.1 | AT3G52380.1 | 15956 | 13614 | 0 | 13420 | 16875 | 17806 | 8232,6 | 7273,7 | 18731 | 17069 | 0 | 0 | 16590 | 80340 | 0 | 0 | 0 | 23437 | 0 | 0 | 14126 | 15869 | 0 | 53510 | 0 | 0 | 0 | 0 |
| P37107 | cpSRP54 | Psat3g032320.1 | AT5G03940.1 | 141230 | 140810 | 133980 | 125790 | 97802 | 112010 | 219540 | 222860 | 201170 | 153580 | 248530 | 153240 | 95920 | 151860 | 143740 | 211100 | 198680 | 133730 | 408910 | 367930 | 102490 | 243740 | 148050 | 171150 | 0 | 0 | 0 | 34149 |
| B9SL58 | PURA2 | Psat3g071360.1 | AT3G57610.1 | 5467,8 | 0 | 0 | 0 | 0 | 3952,3 | 0 | 0 | 2597,3 | 2293,8 | 0 | 0 | 6453,1 | 26949 | 0 | 1751,9 | 4722,4 | 4774,4 | 0 | 0 | 0 | 5289,8 | 0 | 0 | 1174,9 | 0 | 0 | 0 |
| Q84KI6 | SQD1 | Psat3g076360.2 | AT4G33030.1 | 21832 | 43808 | 0 | 0 | 24421 | 28127 | 0 | 12106 | 19160 | 11900 | 0 | 0 | 79781 | 68631 | 0 | 0 | 45992 | 67242 | 0 | 0 | 76884 | 45369 | 0 | 0 | 0 | 0 | 0 | 0 |
| Q43155 | GLTB | Psat3g078160.3 | AT5G04140.1 | 83042 | 82050 | 20010 | 0 | 199850 | 104140 | 11550 | 0 | 139530 | 70321 | 0 | 0 | 127160 | 96252 | 0 | 0 | 125480 | 196740 | 10289 | 0 | 111230 | 103000 | 0 | 0 | 0 | 0 | 0 | 0 |
| G7L2M8 | GRPE | Psat3g085320.3 | AT5G17710.1 | 68029 | 47675 | 29009 | 22521 | 94198 | 76831 | 59089 | 88266 | 209220 | 88187 | 11390 | 15776 | 222360 | 93495 | 32790 | 38088 | 119160 | 51823 | 62173 | 30004 | 50079 | 43928 | 11564 | 7123,8 | 19693 | 26235 | 0 | 25043 |
| Q94AQ8 | PNSB2 | Psat3g089080.1 | AT1G64770.1 | 4803,4 | 9961,2 | 0 | 0 | 6835 | 6016,8 | 3225,6 | 0 | 0 | 6436,3 | 0 | 0 | 4223,3 | 4535 | 0 | 0 | 7495,9 | 0 | 3567,6 | 0 | 0 | 0 | 0 | 0 | 0 | 0 | 0 | 2453,4 |
| P12858 | G3PA | Psat3g091720.1 | AT1G12900 | 2049600 | 1988500 | 285870 | 330660 | 1951900 | 2293600 | 415100 | 355290 | 2725700 | 1768400 | 135570 | 293670 | 5977700 | 3746700 | 227530 | 338950 | 2480300 | 3671700 | 146700 | 185410 | 2454500 | 2486400 | 39001 | 59621 | 148210 | 181890 | 0 | 57673 |
| Q42783 | BCCP | Psat3g102480.1 | AT5G16390 | 16776000 | 13711000 | 40979000 | 32817000 | 22650000 | 14545000 | 34630000 | 29024000 | 15394000 | 19704000 | 47859000 | 36146000 | 11012000 | 11918000 | 48793000 | 61344000 | 16031000 | 12587000 | 48015000 | 37918000 | 40150000 | 23492000 | 94773000 | 96499000 | 31032000 | 32373000 | 522400 | 29937000 |
| P29450 | TRXF | Psat3g102640.1 | AT5G16400 | 123440 | 79815 | 0 | 18081 | 215140 | 84369 | 0 | 0 | 63769 | 53089 | 0 | 0 | 276290 | 213930 | 0 | 0 | 71162 | 85184 | 0 | 0 | 74748 | 88532 | 168600 | 0 | 0 | 0 | 0 | 0 |
| O04603 | RK5 | Psat3g107160.1 | AT4G01310.1 | 6346,1 | 0 | 22845 | 50621 | 0 | 16179 | 61797 | 62503 | 0 | 7506,6 | 110590 | 164300 | 35393 | 76936 | 96294 | 171540 | 129690 | 105560 | 401160 | 83066 | 116670 | 15546 | 180350 | 112970 | 106260 | 111900 | 0 | 377480 |
| P46275 | F16P1 | Psat3g134680.1 | AT3G54050.1 | 25969 | 11151 | 0 | 0 | 17400 | 20678 | 3679,1 | 0 | 21309 | 7791,5 | 0 | 0 | 66040 | 46451 | 0 | 0 | 24574 | 58180 | 0 | 0 | 21948 | 15775 | 0 | 0 | 0 | 0 | 0 | 0 |
| O80934 | Y2766 | Psat3g138600.1 | AT2G37660.1 | 20701 | 36923 | 0 | 2452,6 | 19400 | 14209 | 0 | 0 | 40362 | 19574 | 0 | 2543,2 | 103840 | 87553 | 0 | 0 | 196810 | 147280 | 0 | 0 | 54515 | 22117 | 0 | 0 | 0 | 0 | 0 | 0 |
| Q42561 | FATA1 | Psat3g149200.1 | AT3G25110.1 | 11321 | 7515,2 | 0 | 0 | 7719,9 | 17766 | 0 | 0 | 20880 | 19746 | 0 | 0 | 12378 | 16270 | 0 | 4675,2 | 17742 | 12561 | 0 | 0 | 5113,1 | 0 | 0 | 0 | 0 | 0 | 0 | 0 |
| Q9SLA8 | FABI | Psat3g150560.1 | AT2G05990.1 | 8747,4 | 8683,4 | 0 | 0 | 9951,2 | 0 | 0 | 0 | 49816 | 0 | 0 | 0 | 5917,9 | 8835,1 | 0 | 3545,1 | 10123 | 10560 | 0 | 0 | 8571,5 | 0 | 0 | 0 | 0 | 0 | 0 | 0 |
| Q9FUZ2 | DEF1B | Psat3g166600.1 | AT5G14660.1 | 8125,2 | 8287,9 | 0 | 5649,8 | 0 | 14737 | 0 | 5650,2 | 12833 | 0 | 0 | 0 | 3725,8 | 8779,5 | 0 | 0 | 13403 | 7376,1 | 0 | 0 | 18804 | 0 | 0 | 0 | 0 | 0 | 0 | 0 |
| P82024 | RR21 | Psat3g169680.2 | AT3G27160.1 | 7121,6 | 9672,5 | 2478,7 | 4310 | 10450 | 14510 | 2105,3 | 7498,7 | 9565 | 7754,9 | 11115 | 14067 | 7167 | 5421,3 | 12881 | 16360 | 9735,5 | 10710 | 23391 | 104950 | 9826,1 | 14956 | 10811 | 10019 | 16596 | 16040 | 0 | 49522 |
| Q9LHA8 | MD37C | Psat3g182640.1 | AT3G12580.1 | 3174,2 | 4732,6 | 2806 | 5421,7 | 7216,8 | 7303,4 | 2640,5 | 3956,3 | 10961 | 9094,5 | 3468,7 | 0 | 7748,2 | 4784,9 | 0 | 0 | 0 | 0 | 2634,2 | 0 | 0 | 0 | 2741,4 | 2790,5 | 0 | 0 | 0 | 1854,3 |
| P08215 | ATPA | Psat3g186480.1 | AtCg00120 | 36134 | 22624 | 296010 | 427700 | 43296 | 40741 | 0 | 742090 | 76573 | 61093 | 542950 | 198950 | 30034 | 0 | 114430 | 81331 | 43466 | 23945 | 75855 | 67135 | 21069 | 13916 | 0 | 38533 | 58831 | 65091 | 0 | 80722 |
| Q9BBP8 | RR3 | Psat4g046880.1 | ATCG00800.1 | 81142 | 120490 | 26378 | 24643 | 0 | 0 | 25350 | 27171 | 0 | 39005 | 44395 | 32885 | 129710 | 75541 | 23770 | 33007 | 167300 | 56079 | 66463 | 115090 | 57816 | 179820 | 23705 | 21219 | 17719 | 21416 | 0 | 20934 |
| P55217 | CGS1 | Psat3g199280.1 | AT3G01120.1 | 4581,7 | 0 | 0 | 0 | 0 | 5766,3 | 0 | 2177,7 | 5757,4 | 6523,5 | 0 | 0 | 14262 | 5084,8 | 0 | 0 | 5319,1 | 6786,4 | 0 | 0 | 6622,7 | 0 | 0 | 0 | 0 | 0 | 0 | 0 |
| P00869 | RBS2 | Psat3g205240.1 | AT1G67090 | 8957100 | 8144100 | 804550 | 995690 | 12644000 | 11414000 | 744930 | 786340 | 7421500 | 8615100 | 758840 | 608050 | 17530000 | 16333000 | 892960 | 858920 | 6447800 | 5084000 | 562620 | 269750 | 12820000 | 8439200 | 304770 | 253160 | 359490 | 449320 | 0 | 484240 |
| P28552 | ATPG | Psat3g207840.1 | AT4G04640 | 15045 | 20277 | 76636 | 100140 | 16229 | 12378 | 145990 | 146040 | 18726 | 17674 | 100060 | 86262 | 13964 | 10990 | 33567 | 22390 | 12667 | 12761 | 68557 | 64656 | 0 | 0 | 0 | 13005 | 19760 | 15990 | 0 | 26818 |
| Q55512 | DHAS | Psat4g004640.1 | AT1G14810.1 | 52074 | 0 | 0 | 17728 | 0 | 41851 | 0 | 0 | 60689 | 49250 | 0 | 27206 | 17324 | 87927 | 0 | 0 | 0 | 0 | 120340 | 0 | 35693 | 0 | 0 | 0 | 14927 | 0 | 0 | 0 |
| O80362 | RK10 | Psat4g007600.1 | AT5G13510.1 | 8737,9 | 4424,1 | 17895 | 13809 | 7792,2 | 8860,2 | 14733 | 20017 | 14171 | 20778 | 27442 | 54702 | 13861 | 98217 | 31808 | 30011 | 16774 | 15129 | 190100 | 12569 | 3376,5 | 27116 | 9363,7 | 2688,7 | 19626 | 38827 | 0 | 236150 |
| Q9M098 | EBFC1 | Psat4g009640.2 | AT4G30620.1 | 99269 | 83887 | 151510 | 118840 | 43366 | 57573 | 150290 | 146450 | 102530 | 112700 | 556720 | 270040 | 37118 | 327390 | 200230 | 234770 | 42497 | 47615 | 200630 | 215040 | 25926 | 281240 | 30942 | 40232 | 286320 | 199370 | 0 | 35431 |
| Q9MB35 | PERQ | Psat4g018640.1 | AT3G26060.1 | 403570 | 474480 | 37411 | 42006 | 673360 | 467650 | 60088 | 65884 | 371640 | 329420 | 32337 | 31988 | 820650 | 406340 | 213110 | 256940 | 930350 | 1112700 | 69190 | 50840 | 533830 | 281620 | 27503 | 11180 | 13051 | 0 | 0 | 32086 |
| O65282 | CH20 | Psat4g019040.1 | AT5G20720.1 | 1135400 | 892020 | 219780 | 251800 | 971330 | 950360 | 206870 | 192220 | 786380 | 544780 | 159890 | 344940 | 1744600 | 1132700 | 195990 | 328700 | 613250 | 999620 | 216420 | 118370 | 616360 | 348530 | 30463 | 23225 | 136910 | 245410 | 0 | 93534 |
| A8MS68 | PLPD1 | Psat4g023520.1 | AT3G16950.1 | 318390 | 262030 | 226260 | 305330 | 246950 | 266520 | 385680 | 260170 | 173640 | 290020 | 187460 | 312180 | 93276 | 264390 | 249070 | 179260 | 219910 | 165650 | 42233 | 236430 | 206610 | 335500 | 17835 | 45230 | 15414 | 20845 | 0 | 134470 |
| O78328 | DXS | Psat4g029960.2 | AT4G15560.1 | 77018 | 67592 | 22970 | 13927 | 89010 | 289850 | 24345 | 76790 | 62383 | 0 | 20524 | 11727 | 262850 | 34772 | 10106 | 25018 | 71491 | 74000 | 7801,6 | 27514 | 92364 | 132650 | 0 | 58920 | 24585 | 30029 | 0 | 15305 |
| Q43157 | RPE | Psat4g049960.2 | AT5G61410.1 | 0 | 0 | 429240 | 287760 | 45171 | 41459 | 292370 | 456960 | 0 | 15236 | 326260 | 185730 | 33407 | 89967 | 90619 | 89736 | 84570 | 112990 | 109970 | 114350 | 73538 | 37605 | 0 | 0 | 50736 | 28444 | 0 | 21673 |
| Q9ZS34 | CHLP | Psat4g050200.1 | AT1G74470.1 | 19516 | 39807 | 78339 | 66645 | 32917 | 36052 | 228230 | 183560 | 42524 | 59526 | 127400 | 52894 | 16455 | 27420 | 25459 | 30866 | 31397 | 35044 | 52304 | 22031 | 0 | 8281,5 | 16498 | 0 | 22446 | 17115 | 0 | 11830 |
| Q9SN86 | MDHP | Psat4g051000.1 | AT3G47520.1 | 28957 | 20998 | 5174,3 | 0 | 22856 | 25646 | 8699,5 | 4533 | 27837 | 37100 | 4572,5 | 0 | 31416 | 44706 | 5212,6 | 0 | 112030 | 127080 | 0 | 10222 | 24597 | 106710 | 0 | 0 | 0 | 0 | 0 | 0 |
| Q8W250 | DXR | Psat4g058000.1 | AT5G62790.1 | 9665 | 6817,6 | 11692 | 0 | 5039,8 | 6060,7 | 2692,5 | 3132,9 | 9223,8 | 12517 | 0 | 2650,2 | 23226 | 17150 | 0 | 0 | 13048 | 8895,6 | 0 | 5240,1 | 11961 | 16774 | 0 | 0 | 0 | 0 | 0 | 0 |
| Q9SZX3 | ASSY | Psat4g065240.1 | AT4G24830.1 | 48863 | 33945 | 57995 | 155560 | 34616 | 53241 | 95348 | 171260 | 39690 | 56224 | 59337 | 72258 | 31239 | 164520 | 32880 | 40193 | 12099 | 46747 | 31009 | 97382 | 21741 | 18231 | 0 | 36854 | 14076 | 12148 | 0 | 12677 |
| A0A1S2Z662 | ICT1 | Psat4g068640.1 | At5g19830.1 | 10062 | 41506 | 8055,9 | 12808 | 54492 | 33721 | 16231 | 22892 | 21153 | 23584 | 21300 | 13723 | 13207 | 21801 | 21857 | 15927 | 39194 | 40625 | 10394 | 55012 | 49099 | 66168 | 24940 | 19250 | 0 | 8023,6 | 0 | 11319 |
| I1K0K6 | EFGC2 | Psat4g069360.2 | AT1G62750.1 | 52312 | 36091 | 13505 | 8054,8 | 59185 | 43589 | 21514 | 12311 | 43068 | 47901 | 7533,5 | 11732 | 173600 | 86976 | 0 | 9622 | 60277 | 41972 | 12848 | 0 | 45040 | 27876 | 6525,6 | 0 | 6203,9 | 0 | 0 | 3704,6 |
| A0A072UP88 | TAL | Psat4g070360.2 | AT5G13420.1 | 2359,4 | 2841,2 | 0 | 0 | 2693,8 | 1541,9 | 0 | 2689 | 1748,1 | 2075,9 | 0 | 0 | 2745,2 | 2532,7 | 0 | 0 | 2665,9 | 3343,8 | 0 | 0 | 0 | 6926,3 | 0 | 0 | 0 | 0 | 0 | 0 |
| Q8LB10 | CLPR4 | Psat4g075600.1 | AT4G17040.1 | 107480 | 150990 | 13144 | 11607 | 125540 | 116140 | 22566 | 29618 | 72411 | 57987 | 20160 | 15026 | 118020 | 106830 | 5459,9 | 6474,5 | 142910 | 225300 | 7205,6 | 7572,7 | 156560 | 63787 | 0 | 0 | 24808 | 8043,8 | 0 | 3910 |
| Q9LS25 | PP420 | Psat4g078640.1 | AT5G46580.1 | 11267 | 0 | 21679 | 14181 | 0 | 0 | 13242 | 11787 | 0 | 0 | 0 | 8316,2 | 0 | 0 | 13210 | 8384,3 | 0 | 0 | 0 | 6786,6 | 0 | 0 | 0 | 0 | 10477 | 0 | 0 | 0 |
| Q8LB01 | DAPB2 | Psat4g089800.1 | AT3G59890.1 | 0 | 0 | 0 | 0 | 0 | 0 | 0 | 0 | 0 | 0 | 11495 | 1669,9 | 0 | 0 | 3208,8 | 0 | 0 | 0 | 0 | 0 | 0 | 0 | 0 | 0 | 1824400 | 2270900 | 0 | 195580 |
| Q94ID7 | GGPPS | Psat4g091240.1 | AT4G36810.1 | 0 | 6260,4 | 0 | 0 | 3429 | 4529,7 | 2707,7 | 2809,3 | 6842 | 0 | 0 | 0 | 6674,1 | 6287,2 | 0 | 0 | 4304,1 | 4525,2 | 0 | 0 | 8088,6 | 0 | 0 | 0 | 0 | 0 | 0 | 0 |
| Q8VZ74 | ERA | Psat4g094880.1 | AT5G66470.1 | 5051,3 | 4919,1 | 44873 | 57852 | 5869,5 | 7696,8 | 185120 | 296400 | 6889,2 | 6895,5 | 30905 | 37579 | 14109 | 3853,9 | 55855 | 0 | 0 | 0 | 10059 | 48331 | 3407,1 | 11907 | 0 | 0 | 10646 | 9245,4 | 0 | 10209 |
| P05037 | ATPB | Psat4g098720.1 | ATCG00480.1 | 0 | 0 | 118620 | 167550 | 0 | 0 | 243300 | 146860 | 4582,1 | 5047,2 | 135210 | 142300 | 0 | 0 | 42328 | 47102 | 0 | 0 | 27022 | 57333 | 0 | 0 | 0 | 0 | 5740,7 | 0 | 0 | 3783,8 |
| Q9ZUC1 | AOR | Psat4g123800.1 | AT1G23740.1 | 0 | 0 | 0 | 0 | 7955,1 | 8417,8 | 0 | 0 | 18297 | 13737 | 0 | 0 | 17013 | 7314,5 | 0 | 0 | 24028 | 14904 | 0 | 14285 | 15360 | 11851 | 0 | 0 | 0 | 0 | 0 | 0 |
| Q94A41 | AMY3 | Psat4g132240.1 | AT1G69830.1 | 8538,4 | 8632 | 10528 | 7207,2 | 11098 | 9554,8 | 5147,8 | 7056,3 | 16994 | 8095,6 | 0 | 3220,7 | 10031 | 4914,5 | 5221,6 | 5972,5 | 12583 | 0 | 7341,9 | 4230,7 | 0 | 0 | 0 | 0 | 3528,9 | 4362,5 | 0 | 0 |
| P82130 | RR20 | Psat4g136440.1 | At3g15190.1 | 20679 | 25301 | 15688 | 15180 | 24776 | 37147 | 27785 | 31292 | 26064 | 41292 | 92640 | 57199 | 33151 | 46802 | 34183 | 35779 | 52300 | 57538 | 92501 | 75367 | 52058 | 0 | 36388 | 61219 | 25431 | 31633 | 0 | 143460 |
| Q9SYL9 | RK13 | Psat4g140640.1 | AT1G78630.1 | 0 | 16664 | 16892 | 14555 | 0 | 25064 | 15990 | 0 | 25380 | 21330 | 15548 | 22890 | 38707 | 0 | 0 | 30768 | 0 | 28078 | 49544 | 62170 | 0 | 38746 | 0 | 0 | 0 | 21355 | 0 | 35209 |
| A0A3Q0ETQ4 | A0A3Q0ETQ4 | Psat4g161440.1 | | 0 | 4281,5 | 5509,2 | 4133,6 | 0 | 0 | 3500,8 | 4801,9 | 0 | 7667,6 | 6783,2 | 6169,3 | 0 | 0 | 6219,4 | 8511,1 | 0 | 0 | 6768,7 | 10030 | 0 | 0 | 11083 | 8999,2 | 3780 | 4081,7 | 0 | 6140,7 |
| Q8VY06 | PREP2 | Psat4g213560.1 | AT1G49630.1 | 17267 | 26918 | 75372 | 44587 | 27447 | 24864 | 22876 | 0 | 48927 | 37942 | 24853 | 0 | 48999 | 41442 | 0 | 0 | 45120 | 28760 | 0 | 0 | 0 | 0 | 0 | 0 | 0 | 0 | 0 | 0 |
| Q9FFQ1 | RH31 | Psat4g214280.1 | AT5G63630.1 | 57821 | 10437 | 0 | 1717,5 | 59539 | 108530 | 847,73 | 1279,4 | 11263 | 11972 | 0 | 985,02 | 2055,7 | 25712 | 1632,2 | 1524,1 | 0 | 36887 | 1075,7 | 1998,5 | 4963 | 51661 | 0 | 0 | 954,06 | 1633,5 | 0 | 1188,9 |
| O80361 | RK4 | Psat4g224360.1 | AT1G07320.1 | 107800 | 116840 | 5463 | 108480 | 136810 | 91176 | 20940 | 39018 | 108730 | 132910 | 74057 | 104560 | 102910 | 363000 | 65818 | 119850 | 235990 | 151220 | 136110 | 204840 | 168340 | 165200 | 32039 | 68729 | 52823 | 0 | 0 | 311820 |
| B9HQZ6 | SYAP | Psat5g000840.1 | AT5G22800.1 | 7281,9 | 0 | 251940 | 125890 | 4199,8 | 0 | 78154 | 57607 | 0 | 6990,4 | 201160 | 74037 | 5283 | 0 | 34526 | 26335 | 0 | 11113 | 0 | 2066 | 0 | 0 | 13495 | 0 | 0 | 0 | 0 | 0 |
| B9HQZ6 | SYAP | Psat5g000880.3 | AT5G22800.1 | 0 | 9940,4 | 13729 | 10343 | 21904 | 19594 | 14479 | 11837 | 11047 | 9666,8 | 8193,5 | 9944,6 | 11833 | 18577 | 0 | 0 | 12249 | 15457 | 3237,8 | 18422 | 15263 | 17207 | 0 | 0 | 0 | 0 | 0 | 0 |
| P82191 | RK3 | Psat5g001240.1 | At2g43030.1 | 22200 | 24775 | 15474 | 22206 | 21162 | 22723 | 17786 | 20253 | 23901 | 45526 | 41492 | 46694 | 64509 | 56421 | 43826 | 49214 | 133300 | 68807 | 139280 | 139030 | 63721 | 84123 | 63632 | 28036 | 43564 | 40958 | 0 | 86693 |
| P45621 | GSA | Psat5g001920.1 | AT3G48730.1 | 153790 | 126850 | 32599 | 0 | 193440 | 142540 | 21821 | 34218 | 262120 | 283330 | 20748 | 0 | 398520 | 513590 | 0 | 0 | 320230 | 303470 | 9289,5 | 0 | 171340 | 198440 | 0 | 0 | 0 | 9451,8 | 0 | 0 |
| P52418 | PUR1 | Psat7g181880.1 | AT2G16570.1 | 21017 | 11465 | 0 | 5613,9 | 11513 | 11450 | 11264 | 11593 | 16421 | 0 | 8611,5 | 0 | 15712 | 12157 | 12404 | 9733,9 | 11407 | 0 | 9992,3 | 0 | 0 | 0 | 0 | 0 | 0 | 0 | 0 | 0 |
| P12859 | G3PB | Psat5g010520.1 | AT1G42970.1 | 867420 | 291690 | 29698 | 55262 | 310300 | 157040 | 86749 | 14267 | 190900 | 777790 | 22240 | 36983 | 681110 | 833830 | 0 | 15797 | 485770 | 512580 | 0 | 11282 | 419950 | 362950 | 0 | 0 | 77434 | 3199,1 | 0 | 0 |
| A0A1S2Y854 | A0A1S2Y854 | Psat5g011440.1 | At1G63610 | 11622 | 11045 | 3947,8 | 3897,5 | 6531,1 | 0 | 5272,9 | 4360 | 12159 | 0 | 3146,7 | 5426 | 10893 | 6437,2 | 5328,1 | 0 | 10810 | 11733 | 0 | 3161 | 18312 | 11139 | 0 | 0 | 0 | 0 | 0 | 0 |
| Q9SCL7 | NAGK | Psat5g026120.1 | AT3G57560.1 | 6493,8 | 6400,9 | 10607 | 0 | 6532,5 | 6730,2 | 0 | 0 | 9227,7 | 15160 | 0 | 0 | 0 | 6189,9 | 0 | 0 | 0 | 9670,9 | 0 | 0 | 0 | 0 | 0 | 0 | 0 | 0 | 0 | 0 |
| Q9LEU8 | ARLY | Psat5g048920.1 | AT5G10920.1 | 52826 | 64053 | 0 | 0 | 49485 | 59943 | 0 | 3749,6 | 114880 | 62853 | 0 | 0 | 114350 | 104440 | 18914 | 0 | 77777 | 115880 | 0 | 0 | 87958 | 61060 | 0 | 0 | 0 | 0 | 0 | 0 |
| P28644 | ROC1 | Psat5g050360.1 | AT5G50250 | 15993 | 10341 | 13859 | 17465 | 10020 | 9617,3 | 38368 | 12183 | 9142,6 | 11981 | 7010 | 10378 | 10835 | 10743 | 10896 | 9674,6 | 22611 | 0 | 11505 | 12615 | 9513,4 | 0 | 6814,4 | 0 | 5237,6 | 0 | 0 | 0 |
| P35100 | CLPC | Psat5g054560.2 | AT5G50920.1 | 353060 | 295330 | 36285 | 27971 | 294160 | 248240 | 40048 | 36125 | 378240 | 330400 | 24795 | 18243 | 446510 | 236330 | 13226 | 17309 | 119580 | 161230 | 15812 | 6900,9 | 121450 | 118490 | 9306,4 | 6148,4 | 0 | 6977,2 | 0 | 18448 |
| Q9M3A8 | PP273 | Psat5g077680.1 | AT3G49240.1 | 11284 | 18267 | 3898,8 | 4577,4 | 7664,2 | 0 | 4375,5 | 3808,7 | 0 | 0 | 6582,9 | 5528,4 | 4039,1 | 0 | 0 | 0 | 0 | 0 | 0 | 6079 | 0 | 10042 | 0 | 0 | 0 | 3642,7 | 0 | 1925,7 |
| P48384 | TRXM | Psat5g081600.1 | AT4G03520 | 34182000 | 45137000 | 4635300 | 6004500 | 42164000 | 41749000 | 6132300 | 11132000 | 38349000 | 36651000 | 3027600 | 6394500 | 28162000 | 31847000 | 8695900 | 9557000 | 47582000 | 32721000 | 4628300 | 10261000 | 41418000 | 36556000 | 6457400 | 10366000 | 2386300 | 4869900 | 5266600 | 8688900 |
| P30124 | HEM2 | Psat5g087720.1 | AT1G69740.1 | 102450 | 50532 | 0 | 0 | 74579 | 67970 | 6753,4 | 0 | 135960 | 117420 | 0 | 0 | 146750 | 128090 | 0 | 0 | 128940 | 101440 | 10112 | 0 | 75131 | 112700 | 0 | 0 | 0 | 0 | 0 | 0 |
| Q9SJZ7 | DNJA6 | Psat5g090520.1 | AT2G22360.1 | 147530 | 222260 | 736670 | 764090 | 296310 | 432800 | 487050 | 720510 | 193160 | 629810 | 902870 | 564520 | 156200 | 113330 | 1197600 | 868690 | 476440 | 482930 | 490470 | 274450 | 393610 | 303200 | 982800 | 344300 | 358170 | 369450 | 0 | 355570 |
| Q8LDL0 | HHL1 | Psat5g093920.2 | AT1G67700.1 | 27880 | 12272 | 53648 | 29737 | 25103 | 16062 | 23967 | 29503 | 19081 | 23150 | 0 | 0 | 10510 | 19557 | 18838 | 20780 | 6181,6 | 0 | 13220 | 12930 | 18909 | 19253 | 0 | 0 | 12076 | 0 | 0 | 0 |
| Q93X62 | FABG1 | Psat5g094480.1 | AT1G24360.1 | 166510 | 127950 | 6847,5 | 44238 | 80215 | 81763 | 45552 | 23504 | 46996 | 55295 | 35120 | 29831 | 143390 | 64455 | 49102 | 30305 | 98942 | 39955 | 2773,9 | 9215,6 | 131940 | 123090 | 0 | 0 | 0 | 0 | 0 | 2877 |
| O81852 | AKH2 | Psat5g099120.1 | AT4G19710.1 | 17783 | 40659 | 71055 | 72652 | 36494 | 26467 | 167540 | 144810 | 42845 | 19526 | 114540 | 62607 | 14875 | 0 | 8340,6 | 15388 | 10049 | 0 | 51599 | 0 | 0 | 0 | 6219,5 | 0 | 12756 | 0 | 0 | 11003 |
| F4IFC5 | SYTM2 | Psat5g104480.1 | AT2G04842.1 | 9204,1 | 13504 | 2842,4 | 4183 | 10580 | 11847 | 11520 | 6095,4 | 6893,1 | 0 | 0 | 0 | 12464 | 15849 | 8918,8 | 4632,2 | 13330 | 11652 | 8364,8 | 7688,3 | 12386 | 19667 | 0 | 0 | 0 | 0 | 0 | 0 |
| Q9MAP3 | RK11 | Psat5g110320.1 | AT1G32990.1 | 51143 | 22084 | 11736 | 13783 | 19886 | 48562 | 17212 | 49538 | 0 | 0 | 90851 | 120920 | 41317 | 165430 | 33864 | 70298 | 152600 | 0 | 184390 | 266770 | 246150 | 129650 | 64222 | 52097 | 106730 | 33783 | 0 | 436600 |
| P52417 | GLGS2 | Psat5g110720.1 | AT5G48300.1 | 13541 | 10009 | 2856,4 | 2720,4 | 14950 | 13027 | 1427,4 | 7963,8 | 13926 | 20159 | 2022,5 | 2748,9 | 57102 | 27198 | 0 | 0 | 12367 | 20855 | 0 | 14020 | 30283 | 63813 | 0 | 0 | 3920,6 | 2667,4 | 0 | 0 |
| Q657X6 | EXEC2 | Psat5g114480.1 | AT1G27510.1 | 0 | 2612,4 | 97200 | 82170 | 0 | 0 | 44184 | 76937 | 15662 | 42206 | 30210 | 26510 | 0 | 7456,4 | 19744 | 14126 | 28533 | 40843 | 28045 | 84164 | 31672 | 0 | 0 | 0 | 0 | 0 | 0 | 0 |
| O24310 | EFTU | Psat5g116480.1 | At4g20360 | 482090 | 389580 | 182790 | 118110 | 169610 | 341590 | 364080 | 144980 | 563410 | 444930 | 146250 | 121790 | 1088100 | 585740 | 23585 | 88218 | 1124500 | 546350 | 9581,7 | 15024 | 485520 | 224560 | 0 | 6289,2 | 26264 | 53631 | 0 | 38146 |
| Q04836 | CP31A | Psat5g118040.2 | AT4G24770.1 | 818620 | 220190 | 6075,5 | 14595 | 135560 | 717890 | 4398,9 | 0 | 95731 | 147370 | 0 | 5547,1 | 185660 | 154690 | 6597,1 | 0 | 226460 | 0 | 0 | 5855,3 | 81824 | 306410 | 0 | 0 | 12026 | 11294 | 0 | 6456,1 |
| Q6KGX3 | RR7 | Psat5g119080.1 | AtCg00900.1<br>AtCg01240.1 | 0 | 129190 | 5297,2 | 4741,6 | 111870 | 103080 | 6190,4 | 8558,6 | 56940 | 56138 | 11832 | 10082 | 183350 | 0 | 10998 | 7791,6 | 259110 | 271380 | 63639 | 105750 | 0 | 0 | 53452 | 18891 | 22996 | 25596 | 0 | 67938 |
| O98997 | RCA | Psat5g123320.3 | AT2G39730 | 3398900 | 3185000 | 22038000 | 18447000 | 2937600 | 3496000 | 22683000 | 17332000 | 3567800 | 4206100 | 19481000 | 16865000 | 2214000 | 1623600 | 5508100 | 5605300 | 2152100 | 2045100 | 11039000 | 6628800 | 1590900 | 1214500 | 2114000 | 1573200 | 2080200 | 2473000 | 0 | 2993800 |
| Q06881 | BCCP | Psat5g126160.1 | AT5G15530 | 21321000 | 18962000 | 82616000 | 86284000 | 11762000 | 8664500 | 68506000 | 45913000 | 14151000 | 16829000 | 84078000 | 86357000 | 7558600 | 4484800 | 111620000 | 113810000 | 10675000 | 9707800 | 63910000 | 73042000 | 12171000 | 11444000 | 75795000 | 56564000 | 40684000 | 47647000 | 335050 | 34972000 |
| O20252 | S17P | Psat5g127280.2 | AT3G55800 | 243110 | 274600 | 12800 | 36507 | 243130 | 193220 | 17128 | 43174 | 400610 | 396400 | 11448 | 26242 | 162840 | 447010 | 31184 | 2816,8 | 497490 | 464860 | 4965,3 | 55513 | 468170 | 555580 | 15921 | 48751 | 1809,1 | 13002 | 0 | 8975,2 |
| P42732 | RR13 | Psat5g134240.2 | AT5G14320.1 | 0 | 30953 | 6010,4 | 0 | 26138 | 35869 | 6687,3 | 5135,6 | 8955,7 | 10462 | 12924 | 14810 | 77896 | 82032 | 16614 | 17986 | 42395 | 61090 | 40361 | 161580 | 77896 | 140030 | 8281,3 | 8736 | 11114 | 11368 | 0 | 55527 |
| Q84W56 | RNJ | Psat5g139840.1 | AT5G63420.1 | 5351,3 | 5844,7 | 0 | 0 | 0 | 0 | 0 | 0 | 11063 | 16703 | 0 | 0 | 5423,1 | 7870,6 | 0 | 0 | 6360,3 | 3385,1 | 4035,9 | 0 | 7430,8 | 7658,9 | 0 | 0 | 0 | 0 | 0 | 0 |
| P15820 | RK32 | Psat5g142800.1 | AtCg01020.1 | 0 | 0 | 0 | 0 | 0 | 0 | 0 | 0 | 0 | 0 | 0 | 0 | 0 | 0 | 0 | 0 | 0 | 0 | 0 | 21630 | 0 | 0 | 0 | 27014 | 22318 | 0 | 0 | 27800 |
| O64730 | P2C26 | Psat5g153360.1 | AT2G30170.1 | 24703 | 22724 | 3749,5 | 0 | 17581 | 15908 | 3461 | 3823,9 | 14427 | 25871 | 2052,1 | 3952,8 | 16763 | 9280,8 | 0 | 0 | 13769 | 11627 | 0 | 0 | 0 | 6683,8 | 0 | 3111,5 | 0 | 0 | 0 | 0 |
| Q9M4Y3 | RR10 | Psat5g170120.1 | AT3G13120.1 | 7192,1 | 0 | 0 | 3587,3 | 0 | 0 | 4624,6 | 4373,6 | 0 | 0 | 6357,8 | 5138 | 0 | 0 | 5821,7 | 5842,1 | 3831,2 | 6428,8 | 15919 | 14102 | 0 | 0 | 4024,6 | 5246,9 | 0 | 8365,6 | 0 | 14598 |
| A0A392R6B6 | A0A392R6B6 | Psat5g189080.1 | | 0 | 0 | 77913 | 68304 | 0 | 0 | 122430 | 75305 | 0 | 26023 | 72855 | 41095 | 0 | 0 | 36850 | 37301 | 0 | 0 | 0 | 33038 | 0 | 0 | 0 | 0 | 34055 | 33664 | 0 | 22733 |
| P25269 | TRBP2 | Psat5g194440.1 | AT4G27070.1 | 3155,1 | 0 | 0 | 2439,6 | 0 | 0 | 0 | 0 | 5182,1 | 0 | 0 | 0 | 6216,8 | 4382,4 | 0 | 0 | 0 | 0 | 0 | 0 | 0 | 3757,9 | 0 | 0 | 0 | 0 | 0 | 0 |
| P82190 | RK27 | Psat5g215240.1 | AT5G40950.1 | 23460 | 20246 | 18151 | 40108 | 27548 | 31077 | 41706 | 21892 | 30108 | 32876 | 110580 | 106520 | 36147 | 37731 | 89928 | 113790 | 37004 | 40978 | 194780 | 131800 | 27983 | 0 | 142130 | 122260 | 115650 | 125430 | 0 | 478120 |
| Q9SEU6 | TRXM4 | Psat5g229000.1 | AT3G15360.1 | 0 | 5786,7 | 0 | 0 | 10679 | 14421 | 859,59 | 0 | 4631,1 | 4971,5 | 0 | 0 | 9058,6 | 9039,6 | 0 | 0 | 14796 | 15507 | 0 | 3345 | 15425 | 12334 | 0 | 0 | 0 | 0 | 0 | 0 |
| Q9SEU6 | TRXM4 | Psat5g242680.1 | AT3G15360.1 | 1542700 | 1662100 | 279780 | 501270 | 2150300 | 2125300 | 419360 | 711220 | 2028000 | 1889300 | 165570 | 382170 | 1964600 | 1287400 | 418030 | 414370 | 903000 | 580180 | 742630 | 304430 | 413680 | 1969900 | 516550 | 542370 | 75004 | 122690 | 0 | 484890 |
| Q01289 | POR | Psat5g245240.1 | AT5G54190 | 133310 | 65603 | 47505 | 50438 | 72956 | 91042 | 149330 | 52512 | 132130 | 119350 | 105600 | 42988 | 18393 | 110350 | 28150 | 29513 | 36496 | 64598 | 53008 | 62612 | 38266 | 45624 | 9371,9 | 9972 | 4696,8 | 0 | 0 | 9880 |
| Q8RWG3 | RER6 | Psat5g246640.1 | AT3G56140.1 | 0 | 1898 | 1417,3 | 0 | 0 | 9408,7 | 1826 | 0 | 0 | 0 | 0 | 1533,3 | 1604,5 | 3389,8 | 0 | 0 | 2970,4 | 0 | 1557 | 3546,9 | 3551,9 | 5826,5 | 0 | 0 | 0 | 0 | 0 | 1735,1 |
| P11893 | RK24 | Psat5g249200.1 | At5g54600.1 | 20962 | 0 | 22431 | 20126 | 48709 | 44593 | 31734 | 28498 | 74543 | 58822 | 24480 | 46774 | 32222 | 72357 | 33761 | 35223 | 0 | 19807 | 92026 | 9955 | 43709 | 63324 | 89229 | 46513 | 19181 | 28083 | 0 | 66440 |
| Q42601 | CARB | Psat5g277200.2 | AT1G29900.1 | 244140 | 291560 | 206480 | 241630 | 295030 | 263260 | 256810 | 246310 | 405120 | 401330 | 141590 | 81658 | 782440 | 679520 | 21184 | 48447 | 184860 | 143410 | 45480 | 18842 | 145440 | 139130 | 3792 | 0 | 3753,5 | 19141 | 0 | 4908,6 |
| F4K0E8 | ISPG | Psat5g278240.1 | AT5G60600.1 | 8203,5 | 6907,2 | 13251 | 5653,9 | 0 | 8017 | 11928 | 1323,8 | 9099,3 | 8769,7 | 10970 | 4362,2 | 10291 | 8054,2 | 0 | 0 | 0 | 9697,6 | 0 | 0 | 0 | 8494,5 | 0 | 0 | 0 | 0 | 0 | 0 |
| B5LMQ9 | RK14 | Psat5g284200.1 | AtCg00780.1 | 0 | 0 | 0 | 0 | 0 | 0 | 0 | 0 | 0 | 0 | 0 | 21746 | 0 | 0 | 0 | 0 | 0 | 0 | 0 | 0 | 0 | 0 | 0 | 0 | 15050 | 0 | 0 | 112020 |
| Q9FWS4 | RK31 | Psat6g009240.1 | AT1G75350.1 | 0 | 10724 | 2172,1 | 5088,4 | 0 | 10517 | 6781,5 | 11514 | 16113 | 0 | 12615 | 24898 | 10000 | 0 | 23659 | 10003 | 22691 | 25392 | 22784 | 54944 | 0 | 0 | 34275 | 58795 | 15683 | 11795 | 0 | 26600 |
| P82278 | RR9 | Psat6g010760.1 | At1g74970.1 | 18061 | 0 | 0 | 6824,1 | 19607 | 0 | 24083 | 8068,2 | 19078 | 0 | 81280 | 37620 | 19431 | 31600 | 41282 | 14760 | 0 | 22619 | 155910 | 0 | 22613 | 23483 | 80761 | 15009 | 42161 | 11482 | 0 | 53379 |

| | | | | | | | | | | | | | | | | | | | | | | | | | | | | | | | |
|---|---|---|---|---|---|---|---|---|---|---|---|---|---|---|---|---|---|---|---|---|---|---|---|---|---|---|---|---|---|---|---|
| P08927 | RUBB | Psat6g016920.1 | AT1G55490 | 830200 | 1022900 | 1441500 | 813940 | 1583100 | 1089600 | 1008900 | 527660 | 1118100 | 1089800 | 1204100 | 951890 | 1428600 | 1464400 | 554220 | 511340 | 1368400 | 1605100 | 867770 | 1359900 | 857830 | 461910 | 325300 | 45481 | 279110 | 294020 | 0 | 232580 |
| A0A1S3E5A8 | A0A1S3E5A8 | Psat6g044440.4 | AT1G23180 | 3277,6 | 0 | 7987,8 | 8386 | 7056,9 | 4662,3 | 8984 | 6146,8 | 6942,7 | 6413 | 0 | 5816,8 | 24064 | 4141,1 | 0 | 0 | 6731,3 | 6819,1 | 0 | 8631,3 | 4354,6 | 16504 | 0 | 0 | 3077,9 | 0 | 0 | 0 |
| Q9LU63 | ATP1 | Psat6g049600.1 | AT5G51110.1 | 622330 | 977920 | 136130 | 232110 | 269700 | 738750 | 251260 | 240770 | 452620 | 646040 | 136170 | 187130 | 598310 | 231920 | 96301 | 258190 | 350870 | 637020 | 217030 | 239600 | 721180 | 423720 | 145930 | 122390 | 134650 | 40906 | 0 | 61816 |
| Q42777 | MCCA | Psat6g059920.1 | AT1G03090.1 | 10689 | 0 | 19000 | 11863 | 0 | 0 | 14119 | 13683 | 0 | 0 | 18973 | 16597 | 0 | 5672 | 17070 | 27204 | 0 | 0 | 20150 | 40894 | 0 | 0 | 17930 | 14487 | 14297 | 16729 | 0 | 21696 |
| Q1KPV0 | FZL | Psat6g060000.3 | AT1G03160.1 | 4700,5 | 0 | 7915,9 | 4465,1 | 14613 | 0 | 8954,9 | 5243,3 | 9181,8 | 14789 | 6793,1 | 7237,5 | 0 | 0 | 0 | 0 | 0 | 0 | 6382,3 | 4537,2 | 0 | 0 | 0 | 0 | 0 | 0 | 0 | 0 |
| Q9M1X0 | RRFC | Psat6g060640.1 | AT3G63190.1 | 119770 | 313140 | 91775 | 128490 | 267090 | 455620 | 118880 | 87683 | 80789 | 78862 | 46770 | 46942 | 281200 | 324450 | 77303 | 299080 | 88353 | 363250 | 107160 | 40384 | 330650 | 360820 | 43673 | 11510 | 3900,8 | 25998 | 0 | 11673 |
| Q9ZUU4 | CP29B | Psat6g071720.1 | AT2G37220.1 | 167060 | 171480 | 29456 | 13729 | 220550 | 172860 | 58129 | 4090,6 | 142300 | 78553 | 39065 | 39370 | 304450 | 354860 | 51467 | 36974 | 168210 | 174150 | 84378 | 11711 | 111650 | 40772 | 16736 | 29428 | 47464 | 69580 | 0 | 22340 |
| O82533 | FTZ21 | Psat6g081080.2 | AT2G36250.1 | 28907 | 8649,8 | 4882,6 | 5448 | 40073 | 33108 | 13265 | 14233 | 25713 | 26142 | 1771,9 | 4432,3 | 11781 | 29918 | 2700,3 | 4354,5 | 55676 | 60981 | 2964,9 | 6356,3 | 49420 | 59371 | 1792,7 | 0 | 0 | 0 | 0 | 11548 |
| G7I7C8 | G7I7C8 | Psat6g095240.1 | AT5G04440 | 3503,5 | 0 | 2842,7 | 0 | 0 | 0 | 0 | 2244,3 | 0 | 0 | 2885,5 | 0 | 0 | 0 | 0 | 0 | 0 | 0 | 0 | 0 | 0 | 0 | 0 | 0 | 0 | 0 | 0 | 0 |
| Q9C7Y2 | MORF5 | Psat6g105080.3 | AT1G32580.1 | 7090,8 | 0 | 0 | 0 | 12124 | 11179 | 0 | 0 | 0 | 12529 | 1717,9 | 0 | 12796 | 14173 | 0 | 0 | 22601 | 24804 | 1731,9 | 3993,7 | 6263,8 | 11479 | 0 | 0 | 0 | 0 | 0 | 0 |
| Q94AR8 | LEUC | Psat6g107560.1 | AT4G13430.1 | 47485 | 27539 | 22429 | 83920 | 34616 | 22427 | 68477 | 85888 | 0 | 78397 | 33862 | 96503 | 31928 | 45983 | 0 | 0 | 40005 | 53795 | 0 | 12607 | 0 | 0 | 0 | 23133 | 10762 | 0 | 0 | 0 |
| Q43117 | KPYA | Psat6g113960.1 | AT3G22960.1 | 2253900 | 3198100 | 1385100 | 1334600 | 3847900 | 4797500 | 2021500 | 2215700 | 4219700 | 4407400 | 1683700 | 1182300 | 2463400 | 2051100 | 1760300 | 2084600 | 1855600 | 1698200 | 1196200 | 1581000 | 3205600 | 1006400 | 1135800 | 806840 | 177240 | 260890 | 0 | 1070800 |
| Q9FNB0 | CHLH | Psat6g120040.1 | AT5G13630.1 | 5371,9 | 9072,3 | 196690 | 146610 | 8670,7 | 13891 | 235250 | 221060 | 7961,7 | 17681 | 174000 | 151190 | 9356,9 | 13273 | 57548 | 36936 | 3345,7 | 6301,4 | 87223 | 24559 | 0 | 0 | 0 | 0 | 10830 | 0 | 0 | 14378 |
| F4K410 | SVR3 | Psat6g127960.2 | AT5G13650.1 | 30347 | 18350 | 48902 | 8221,7 | 37325 | 25721 | 7018,7 | 0 | 34189 | 19587 | 0 | 13696 | 123860 | 31420 | 0 | 2893,9 | 0 | 49568 | 12019 | 4999,6 | 107790 | 0 | 0 | 0 | 0 | 3001,9 | 0 | 0 |
| Q9LVP0 | Y5639 | Psat6g141840.1 | AT5G63930.1 | 0 | 0 | 0 | 0 | 2035,5 | 0 | 0 | 0 | 2213,9 | 0 | 0 | 0 | 1741,5 | 0 | 0 | 0 | 0 | 2770,1 | 0 | 2090,2 | 1226,7 | 0 | 0 | 0 | 0 | 0 | 0 | 0 |
| Q9SR19 | RAF2 | Psat6g143080.1 | AT3G04550.1 | 807350 | 1306200 | 165290 | 234330 | 1197600 | 966780 | 156000 | 154580 | 1445700 | 888540 | 80767 | 98108 | 896520 | 1420900 | 86195 | 46139 | 1427900 | 1632800 | 34394 | 97030 | 1917400 | 2439800 | 6563,9 | 41526 | 56276 | 15165 | 0 | 35270 |
| O81983 | SYS | Psat6g149440.1 | At1g11870.1 | 0 | 0 | 23021 | 0 | 0 | 0 | 0 | 22438 | 0 | 0 | 33753 | 0 | 0 | 0 | 0 | 46959 | 5958,4 | 11205 | 44340 | 45349 | 0 | 0 | 0 | 0 | 91049 | 40275 | 0 | 279330 |
| I3SWS0 | I3SWS0 | Psat6g155960.2 | At5g53850 | 10898 | 4899,7 | 0 | 0 | 0 | 3363,7 | 0 | 0 | 0 | 3161,6 | 0 | 0 | 0 | 3033,6 | 0 | 0 | 0 | 3525,3 | 0 | 0 | 5222,1 | 4869,2 | 2153,2 | 0 | 0 | 0 | 0 | 0 |
| Q41651 | CYPB | Psat6g157800.1 | AT3G62030.1 | 42795 | 24794 | 0 | 0 | 38797 | 40875 | 0 | 0 | 18796 | 30942 | 0 | 9576,5 | 172740 | 85608 | 0 | 0 | 62281 | 51529 | 6577,5 | 0 | 25608 | 75931 | 0 | 0 | 0 | 0 | 0 | 0 |
| Q9S834 | CLPP5 | Psat6g161200.1 | AT1G02560.1 | 21816 | 32391 | 35447 | 17767 | 42173 | 34830 | 24363 | 36113 | 39347 | 52497 | 28720 | 24893 | 52126 | 51903 | 26844 | 0 | 28894 | 22323 | 14455 | 17319 | 25123 | 24988 | 7946,4 | 6352,9 | 10578 | 0 | 0 | 0 |
| O82261 | DEGP2 | Psat6g167960.1 | AT2G47940.1 | 14253 | 12493 | 0 | 0 | 10750 | 13798 | 5591,5 | 0 | 17629 | 12818 | 0 | 14325 | 17293 | 15367 | 0 | 0 | 9295,6 | 14101 | 0 | 0 | 12749 | 19514 | 0 | 0 | 0 | 0 | 0 | 0 |
| Q05046 | CH62 | Psat6g170600.1 | AT3G23990 | 1969000 | 2723900 | 450490 | 0 | 2420400 | 3229500 | 533420 | 0 | 2905800 | 3182700 | 481490 | 0 | 1925500 | 2526100 | 0 | 0 | 4842100 | 6599500 | 339610 | 1004900 | 3303200 | 3260400 | 95495 | 129590 | 0 | 213260 | 0 | 281700 |
| A4GGC9 | RR12 | Psat6g178680.1 | ATCG00065.1 | 20347 | 14238 | 0 | 0 | 13999 | 23838 | 2244,2 | 0 | 0 | 15473 | 3553,7 | 3698,5 | 48592 | 32076 | 0 | 0 | 44585 | 44473 | 37412 | 35628 | 30323 | 100110 | 3407,3 | 0 | 4129,1 | 5472,9 | 0 | 8806 |
| P51412 | RK21 | Psat6g187560.1 | AT1G35680.1 | 0 | 0 | 28503 | 25502 | 0 | 14875 | 36262 | 39998 | 11522 | 21803 | 60629 | 65090 | 6584,3 | 14424 | 34688 | 29556 | 0 | 41420 | 179120 | 85963 | 0 | 17378 | 57596 | 36563 | 34913 | 23642 | 0 | 74286 |
| Q9FLW9 | PKP2 | Psat6g194760.1 | AT5G52920.1 | 18283 | 15633 | 0 | 0 | 24969 | 22198 | 1625,6 | 3414,8 | 12394 | 9623 | 1981,8 | 0 | 18825 | 23511 | 0 | 0 | 51611 | 43672 | 1995,8 | 0 | 36768 | 35187 | 0 | 2843,3 | 0 | 0 | 0 | 0 |
| F6H7K5 | THI42 | Psat6g209120.1 | AT5G54770.1 | 22084 | 31160 | 68804 | 47078 | 34153 | 24638 | 115680 | 9676,3 | 46892 | 26547 | 42545 | 8761,1 | 34865 | 30931 | 0 | 0 | 49327 | 57789 | 0 | 7742,9 | 53795 | 30596 | 0 | 0 | 0 | 3861,4 | 0 | 0 |
| Q9XJ35 | CLPR1 | Psat6g215320.4 | AT1G49970.1 | 55671 | 44106 | 0 | 0 | 14933 | 32955 | 5324,6 | 0 | 21230 | 13720 | 8858,3 | 0 | 72610 | 22012 | 6303,7 | 0 | 23797 | 20408 | 8439,9 | 7367,7 | 39191 | 11102 | 0 | 0 | 10708 | 0 | 0 | 6087,8 |
| O49485 | SERA1 | Psat6g219520.1 | AT4G34200.1 | 0 | 11749 | 0 | 6929,6 | 13530 | 0 | 11160 | 8974,4 | 47461 | 16657 | 10480 | 8414,9 | 17918 | 20667 | 0 | 6871,8 | 0 | 0 | 0 | 4508 | 0 | 0 | 0 | 0 | 0 | 0 | 0 | 4616,5 |
| P43394 | K502 | Psat6g222520.1 | AT1G15140.1 | 25799 | 35464 | 69520 | 148720 | 50017 | 37829 | 169270 | 114010 | 0 | 17653 | 44988 | 29890 | 13913 | 30972 | 11120 | 28277 | 65431 | 31081 | 95976 | 70804 | 0 | 0 | 9069,8 | 0 | 9105,4 | 0 | 0 | 11083 |
| Q9ZU52 | ALFP3 | Psat6g223200.1 | AT2G01140.1 | 75518 | 45500 | 14065 | 0 | 87061 | 122090 | 0 | 0 | 86410 | 152470 | 17108 | 0 | 145310 | 116810 | 0 | 0 | 92112 | 101300 | 0 | 15739 | 84912 | 82205 | 0 | 0 | 0 | 0 | 0 | 0 |
| Q9C9F5 | C3H15 | Psat6g223920.1 | AT1G68200.1 | 3626,8 | 0 | 7888,2 | 2622,5 | 0 | 0 | 8845,2 | 12497 | 0 | 0 | 9185,3 | 0 | 0 | 0 | 9707,4 | 9555 | 0 | 0 | 6918,5 | 0 | 0 | 0 | 0 | 0 | 6542,7 | 0 | 0 | 4594,7 |
| P31165 | RK15 | Psat6g225000.1 | At3g25920.1 | 9420,3 | 10282 | 19878 | 21554 | 10195 | 11192 | 24455 | 7532,3 | 10451 | 9095,2 | 62117 | 53325 | 4397 | 64499 | 43521 | 33665 | 0 | 56492 | 140020 | 225640 | 28500 | 15718 | 126000 | 63542 | 64404 | 161980 | 0 | 154410 |
| Q9SQI8 | ODP24 | Psat6g228960.1 | AT3G25860.1 | 235520 | 151990 | 299000 | 236830 | 203990 | 150360 | 313140 | 237790 | 192620 | 142860 | 269390 | 159390 | 160540 | 134100 | 349090 | 204430 | 149310 | 177910 | 193330 | 109630 | 241890 | 126970 | 211650 | 195260 | 41231 | 5302,7 | 0 | 233210 |
| A0A2K3P005 | COBWD | Psat6g237480.1 | AT1G80480.1 | 24775 | 20979 | 12325 | 23715 | 34400 | 32252 | 40254 | 13678 | 26943 | 35906 | 0 | 0 | 19681 | 28154 | 0 | 0 | 43720 | 25557 | 0 | 14132 | 30837 | 22467 | 0 | 0 | 0 | 0 | 0 | 0 |
| F4IW47 | TKTC2 | Psat6g238800.1 | AT2G45290.2 | 1793400 | 2427300 | 101080 | 116240 | 1958400 | 1599500 | 162630 | 159420 | 2361100 | 2153300 | 129940 | 92875 | 5244300 | 4689100 | 93875 | 73776 | 3013100 | 2506500 | 85453 | 4577,3 | 1442300 | 730400 | 28751 | 49715 | 74442 | 65977 | 0 | 33248 |
| Q9C550 | LEU12 | Psat6g241040.1 | AT1G74040.1 | 29084 | 84489 | 25264 | 15373 | 39590 | 64546 | 30864 | 31158 | 64091 | 61677 | 30300 | 22530 | 60052 | 21031 | 20144 | 0 | 66987 | 37719 | 13997 | 343150 | 48491 | 58745 | 14816 | 0 | 49633 | 19637 | 0 | 10903 |
| Q6ZY51 | PWD | Psat6g244000.1 | AT5G26570.1 | 20517 | 12487 | 0 | 0 | 22644 | 21082 | 13179 | 0 | 19341 | 20814 | 0 | 0 | 20409 | 20130 | 0 | 0 | 16085 | 22050 | 0 | 6364,6 | 0 | 17579 | 0 | 0 | 0 | 0 | 0 | 0 |
| O82234 | IF32 | Psat7g006440.1 | AT2G24060.1 | 382110 | 452620 | 273970 | 265720 | 271590 | 303420 | 483040 | 293860 | 420110 | 343350 | 306110 | 271310 | 363910 | 129730 | 305420 | 220060 | 143910 | 150990 | 560460 | 265740 | 77841 | 179660 | 572580 | 622320 | 67332 | 98487 | 0 | 159310 |
| Q8VZ55 | RK35 | Psat7g006560.1 | AT2G24090.1 | 0 | 0 | 0 | 0 | 0 | 11501 | 9582,4 | 0 | 10967 | 12582 | 39621 | 37499 | 9931,2 | 15367 | 0 | 35174 | 0 | 29321 | 84093 | 48814 | 16685 | 19542 | 32935 | 56709 | 0 | 0 | 0 | 97147 |
| Q8VY91 | RR6 | Psat7g023960.1 | AT1G64510.1 | 5433,5 | 0 | 19798 | 14354 | 0 | 4832,5 | 0 | 0 | 0 | 0 | 33639 | 26330 | 12120 | 8143,8 | 0 | 40686 | 0 | 7737,6 | 53095 | 40276 | 0 | 9086,2 | 44467 | 75629 | 31545 | 22710 | 0 | 57560 |
| Q9SH69 | 6PGD1 | Psat7g028480.1 | AT1G64190.1 | 8194,6 | 15642 | 0 | 0 | 15109 | 18527 | 0 | 0 | 22481 | 25567 | 0 | 7683,6 | 25206 | 21065 | 0 | 0 | 31873 | 24898 | 0 | 0 | 15996 | 0 | 0 | 0 | 0 | 0 | 0 | 0 |
| P35100 | CLPC | Psat7g039080.1 | AT5G50920.1 | 1779900 | 1458700 | 379890 | 571280 | 1980200 | 2288600 | 559370 | 511210 | 1883200 | 2223400 | 305320 | 316360 | 1556000 | 1675200 | 109100 | 128990 | 2511400 | 1858900 | 215680 | 204100 | 2057200 | 1857300 | 46492 | 10550 | 62417 | 46105 | 0 | 135140 |
| Q9SW18 | CHLM | Psat7g039320.1 | AT4G25080.1 | 186170 | 121050 | 106480 | 144300 | 51432 | 41791 | 240540 | 265940 | 61176 | 55106 | 90160 | 110600 | 60735 | 29073 | 32527 | 38930 | 111760 | 135500 | 101440 | 113190 | 0 | 11647 | 14698 | 20739 | 11157 | 10833 | 0 | 22801 |
| P82248 | RK29 | Psat7g055240.1 | At5g65220.1 | 50948 | 46399 | 9979,7 | 16360 | 63474 | 109120 | 48854 | 37228 | 32740 | 37017 | 44474 | 94455 | 83430 | 92308 | 74193 | 101430 | 59540 | 68026 | 137610 | 182200 | 54441 | 124430 | 62137 | 65072 | 48149 | 48818 | 0 | 201350 |
| Q93YZ7 | ILVH2 | Psat7g063320.1 | AT2G31810.1 | 29267 | 25634 | 21666 | 19770 | 24035 | 23109 | 16859 | 30676 | 15136 | 19390 | 20791 | 20041 | 41806 | 23491 | 8128,9 | 5052,9 | 0 | 23966 | 9027,1 | 23339 | 67662 | 58690 | 0 | 0 | 2839,3 | 3197 | 0 | 12970 |
| P37142 | AKH | Psat7g075960.1 | AT4G19710.1 | 95998 | 94014 | 123670 | 161560 | 110490 | 135260 | 183970 | 344530 | 117430 | 79100 | 120020 | 91840 | 90647 | 52024 | 22627 | 34765 | 50935 | 21336 | 66622 | 97740 | 2942,8 | 70914 | 6804,3 | 0 | 13763 | 4855,6 | 0 | 18734 |
| Q93ZN9 | DAPAT | Psat7g104520.1 | AT4G33680.1 | 114810 | 41027 | 7269,8 | 2455 | 80948 | 30999 | 2396,3 | 8749,5 | 30293 | 85349 | 0 | 1544,5 | 44147 | 150320 | 0 | 0 | 204610 | 279020 | 57890 | 3853,2 | 62738 | 74358 | 0 | 0 | 3656,9 | 1112,4 | 0 | 0 |
| O81004 | Y2287 | Psat7g113880.1 | AT2G22870.1 | 0 | 0 | 10210 | 7470,4 | 0 | 0 | 6984,2 | 20774 | 0 | 0 | 10638 | 22780 | 0 | 0 | 9210,1 | 10164 | 0 | 0 | 17590 | 0 | 0 | 0 | 0 | 9879 | 6881 | 0 | 0 | 4415,1 |
| Q9ST69 | RR5 | Psat7g115360.1 | At2g33800.1 | 99914 | 95618 | 30244 | 0 | 37184 | 125290 | 36277 | 39553 | 74860 | 137310 | 80905 | 104600 | 43443 | 97253 | 87076 | 106390 | 182600 | 128260 | 110990 | 159880 | 91402 | 222880 | 57582 | 71781 | 60340 | 34320 | 0 | 168990 |
| Q42807 | STAD | Psat7g119680.1 | AT2G43710.1 | 8398,7 | 9333,2 | 2953,5 | 0 | 14310 | 21423 | 0 | 1238,3 | 22528 | 23150 | 1816,2 | 0 | 17069 | 43088 | 0 | 0 | 76393 | 16972 | 0 | 0 | 35731 | 25038 | 0 | 0 | 1994,8 | 1535,3 | 0 | 0 |
| Q01516 | ALFC1 | Psat7g139080.1 | AT2G21330 | 481250 | 565030 | 6879,6 | 0 | 455960 | 473340 | 13863 | 24967 | 1287800 | 867700 | 8980,2 | 0 | 1295500 | 941730 | 0 | 0 | 436330 | 613640 | 10745 | 0 | 312480 | 447580 | 5440,4 | 0 | 9216,4 | 0 | 0 | 5188,8 |
| Q01516 | ALFC1 | Psat7g139120.1 | AT2G21330 | 2293400 | 2177600 | 38932 | 78352 | 2522600 | 2410200 | 163100 | 152870 | 4863500 | 5478500 | 93164 | 21518 | 6739800 | 4537000 | 162350 | 158370 | 4710700 | 3878600 | 98428 | 17267 | 2463100 | 1782100 | 26626 | 67294 | 207030 | 266750 | 0 | 99434 |
| P08926 | RUBA | Psat7g144320.1 | AT2G28000.1 | 14698000 | 13420000 | 25213000 | 18517000 | 15859000 | 14382000 | 16148000 | 14054000 | 17267000 | 17747000 | 24645000 | 17081000 | 14456000 | 14996000 | 9216900 | 10522000 | 17766000 | 18355000 | 16472000 | 10284000 | 13747000 | 9558800 | 4198900 | 3772600 | 4995200 | 3427200 | 0 | 4846600 |
| Q43088 | RBCMT | Psat7g156240.1 | AT1G14030.1 | 0 | 14589 | 506820 | 169530 | 35849 | 0 | 534550 | 380480 | 17236 | 20815 | 400190 | 250090 | 3582,4 | 0 | 17888 | 72131 | 0 | 0 | 162520 | 46935 | 0 | 0 | 29338 | 0 | 38455 | 29136 | 0 | 38766 |
| O82043 | ILV5 | Psat7g165680.3 | AT3G58610.1 | 0 | 21412 | 421530 | 320480 | 22486 | 22019 | 692060 | 510600 | 20477 | 20542 | 397700 | 214290 | 116390 | 96566 | 98032 | 107290 | 29555 | 21618 | 221660 | 135960 | 27249 | 34196 | 0 | 35800 | 81316 | 48256 | 0 | 22319 |
| Q43082 | HEM3 | Psat7g172760.1 | AT5G08280.1 | 24659 | 14262 | 32198 | 7591,3 | 14373 | 25598 | 3345,2 | 3562,2 | 14138 | 26250 | 0 | 4013,6 | 195340 | 287160 | 0 | 0 | 52216 | 27753 | 12028 | 6452,2 | 19493 | 114700 | 0 | 0 | 2475,9 | 0 | 0 | 0 |
| O22265 | cpSRP43 | Psat7g178840.1 | AT2G47450.1 | 40779 | 145100 | 32868 | 42953 | 37219 | 77896 | 31328 | 24236 | 54045 | 20819 | 42778 | 13157 | 37815 | 36885 | 30521 | 0 | 60033 | 131370 | 36668 | 38195 | 37629 | 113690 | 53675 | 0 | 8056 | 0 | 0 | 0 |
| Q94B35 | ISPH | Psat7g185320.1 | AT4G34350.1 | 24894 | 16721 | 15381 | 12317 | 12687 | 12179 | 15518 | 20640 | 10779 | 7727,7 | 6108,3 | 10981 | 10876 | 20726 | 12394 | 9061,7 | 43064 | 0 | 0 | 14012 | 17765 | 0 | 0 | 0 | 7591,7 | 0 | 0 | 0 |
| Q8SKU2 | TIC62 | Psat7g185400.1 | AT3G18890.1 | 0 | 0 | 251590 | 311850 | 0 | 0 | 627310 | 360540 | 0 | 56080 | 364190 | 274830 | 0 | 13988 | 240870 | 297550 | 0 | 0 | 59415 | 249520 | 0 | 0 | 76228 | 262800 | 144550 | 220700 | 0 | 73647 |
| Q9LY66 | RK1 | Psat7g200640.1 | AT3G63490.1 | 25772 | 87606 | 131960 | 76715 | 39825 | 39228 | 105560 | 138860 | 172750 | 170660 | 273690 | 336420 | 44685 | 47475 | 300150 | 391340 | 91539 | 176790 | 331490 | 811590 | 42215 | 49351 | 91731 | 149700 | 261320 | 234800 | 0 | 819190 |
| Q03943 | IM30 | Psat7g208480.2 | AT1G65260.1 | 245480 | 194380 | 3097200 | 1499900 | 134010 | 144250 | 1450700 | 1134600 | 135490 | 241660 | 762290 | 930770 | 87521 | 128720 | 557770 | 465140 | 61409 | 71943 | 1140300 | 794490 | 295390 | 121770 | 1404200 | 2130000 | 131080 | 92018 | 63272 | 457700 |
| P11894 | RK9 | Psat7g223760.1 | At3g44890.1 | 123490 | 176660 | 123360 | 92267 | 91848 | 200390 | 118880 | 102480 | 101930 | 167130 | 279430 | 313940 | 264360 | 154160 | 499280 | 250420 | 64967 | 42235 | 702280 | 430180 | 40810 | 59514 | 380990 | 322450 | 127250 | 104040 | 0 | 693080 |
| G7JR97 | G7JR97 | Psat7g241800.1 | | 20180 | 32156 | 3604,9 | 0 | 27072 | 26233 | 4179,9 | 2944,7 | 43362 | 26769 | 0 | 0 | 13934 | 11230 | 0 | 3356,6 | 9505,3 | 6375,5 | 0 | 1801,1 | 13917 | 18013 | 0 | 0 | 0 | 3704 | 0 | 5136,1 |
| Q9M394 | SCKL1 | Psat7g243520.1 | AT3G54090.1 | 26729 | 12718 | 1373,3 | 0 | 60625 | 36583 | 20159 | 4868,8 | 16928 | 39153 | 1016,9 | 0 | 30890 | 13446 | 0 | 2507,1 | 67063 | 69963 | 1209,4 | 0 | 42812 | 4126,4 | 0 | 0 | 0 | 0 | 0 | 0 |
| A0A2K3NKT1 | A0A2K3NKT1 | Psat7g260440.1 | | 14060 | 13645 | 107020 | 60651 | 19139 | 17602 | 112270 | 138360 | 11121 | 28554 | 131850 | 69220 | 9059,5 | 8818,9 | 16921 | 27361 | 0 | 0 | 54795 | 23936 | 0 | 0 | 0 | 0 | 0 | 0 | 0 | 3974 |
